# Admixture Into and Within Sub-Saharan Africa

**DOI:** 10.1101/038406

**Authors:** George B.J. Busby, Gavin Band, Quang Si Le, Muminatou Jallow, Edith Bougama, Valentina Mangano, Lucas Amenga-Etego, Anthony Emimil, Tobias Apinjoh, Carolyne Ndila, Alphaxard Manjurano, Vysaul Nyirongo, Ogobara Doumbo, Kirk A. Rockett, Domnic P. Kwiatkowski, Chris C.A. Spencer, In Association with the Malaria Genomic Epidemiology Network

## Abstract

Understanding patterns of genetic diversity is a crucial component of medical research in Africa. Here we use haplotype-based population genetics inference to describe gene-flow and admixture in a collection of 48 African groups with a focus on the major populations of the sub-Sahara. Our analysis presents a framework for interpreting haplotype diversity within and between population groups and provides a demographic foundation for genetic epidemiology in Africa. We show that coastal African populations have experienced an influx of Eurasian haplotypes as a series of admixture events over the last 7,000 years, and that Niger-Congo speaking groups from East and Southern Africa share ancestry with Central West Africans as a result of recent population expansions associated with the adoption of new agricultural technologies. We demonstrate that most sub-Saharan populations share ancestry with groups from outside of their current geographic region as a result of large-scale population movements over the last 4,000 years. Our in-depth analysis of admixture provides an insight into haplotype sharing across different geographic groups and the recent movement of alleles into new climatic and pathogenic environments, both of which will aid the interpretation of genetic studies of disease in sub-Saharan Africa.

## Introduction

Genetic epidemiological studies aim to uncover novel relationships between genes, the environment, and disease [Malaria Genomic Epidemiology Network, 2015]. A critical component of this work is a description of patterns of genetic variation and an understanding of the genetic structure of populations. Whilst tens of thousands of genetic variants have been associated with different diseases in populations of European descent [Welter et al., 2014], despite the high burden of disease in Africa, medical genetic research on the continent has lagged behind [Need and Goldstein, 2009]. To alleviate this, several broad consortia are beginning to focus on understanding the genetic basis of infectious and non-communicable disease specifically in Africa [Malaria Genomic Epidemiology Network, 2008; H3Africa Consortium, 2014; Gurdasani et al., 2014; Malaria Genomic Epidemiology Network, 2015], and a number of recent studies have sought to describe patterns of genetic variation across the continent [Campbell and Tishkoff, 2008; Tishkoff et al., 2009; Gurdasani et al., 2014].

Genome-wide analyses of African populations are refining previous models of the continent’s genetic history. One such emerging insight is the identification of clear, but complex, evidence for the movement of Eurasian ancestry back into the continent as a result of admixture over a variety of timescales [Pagani et al., 2012; Pickrell et al., 2014; Gurdasani et al., 2014; Hodgson et al., 2014a; Llorente et al., 2015]. Admixture occurs when genetically differentiated ancestral groups come together and mix, a process which is increasingly regarded as a common feature of human populations across the globe [Patterson et al., 2012; Hellenthal et al., 2014; Busby et al., 2015]. In a broad sample of 18 ethnic groups from eight countries, the African Genome Variation Project (AGVP) [Gurdasani et al., 2014] recreated previously published results to identify recent Eurasian admixture within the last 1.5 thousand years (ky) in the Fulani of West Africa [Tishkoff et al., 2009; Henn et al., 2012] and several East African groups from Kenya, older Eurasian ancestry (2-5 ky) in Ethiopian groups, consistent with previous studies of similar populations [Pagani et al., 2012; Pickrell et al., 2014], and a novel signal of ancient (>7.5 ky) Eurasian admixture in the Yoruba of Central West Africa [Gurdasani et al., 2014]. Comparisons of contemporary sub-Saharan African populations with the first ancient genome from within Africa, a 4.5 ky Ethiopian individual [Llorente et al., 2015], provide additional support for limited migration of Eurasian ancestry back into East Africa within the last 3,000 years.

Within this timescale, the major demographic change within Africa was the transition from hunting and gathering to pastoralist and agricultural lifestyles [Diamond and Bellwood, 2003; Smith, 2005; Barham and Mitchell, 2008; Li et al., 2014]. This shift was long and complex and occurred at different speeds, instigating contrasting interactions between the agriculturalist pioneers and the inhabitant people [Mitchell, 2002; Marks et al., 2014]. The change was initialised by the spread of pastoralism (i.e. the raising and herding of livestock) across Africa and the subsequent movement east and south from Central West Africa of agricultural technology together with the branch of Niger-Congo languages known as Bantu [Mitchell, 2002; Barham and Mitchell, 2008]. The AGVP also found evidence of widespread hunter-gatherer ancestry in African populations, including ancient (9 ky) Khoesan ancestry in the Igbo from Nigeria, and more recent hunter-gatherer ancestry in eastern (2.5-4.5 ky) and southern (0.9-4 ky) African populations [Gurdasani et al., 2014]. The identification of hunter-gatherer ancestry in non-hunter-gatherer populations together with the timing of these latter events is consistent with the known expansion of Bantu languages across African within the last 3 ky [Mitchell, 2002; Diamond and Bellwood, 2003; Smith, 2005; Barham and Mitchell, 2008; Marks et al., 2014; Li et al., 2014].

These studies have described the novel and important influence of both Eurasian and hunter-gatherer ancestry on the population genetic history of sub-Saharan Africa. Nevertheless, there is still more to understand about the nature and timing of admixture across Africa. For example, do we observe Eurasian ancestry in additional African populations? Do alternative dating methods recreate a similar timescale for these admixtures? Can we understand more about how the Eurasian ancestry entered African populations? Uncertainty about the relationships between populations within Africa also remains. For example, can we use genetics to refine the origins, interactions, and timings of the Bantu expansion? What did the ancestry of pre-Bantu populations of east and southern Africa look like? What other historical relationships can be characterised between different African ethno-linguistic groups?

Here we review these questions using the same methods as previous authors, and provide additional novel inference by using methods that utilise haplotype information. To gain a detailed understanding of the population structure and history of Africa, we analyse genome-wide data from 14 Eurasian and 46 sub-Saharan African groups. Half (23) of the African groups represent subsets of samples collected from nine countries as part of the MalariaGEN consortium. Details on the recruitment of samples in relation to studying malaria genetics are published elsewhere [Malaria Genomic Epidemiology Network, 2014, 2015]. The remaining 23 groups are from publicly available datasets from a further eight sub-Saharan African countries [Pagani et al., 2012; Schlebusch et al., 2012; Petersen et al., 2013] and the 1000 Genomes Project (1KGP), with Eurasian groups from the latter included to help understand the genetic contribution from outside of the continent (Figure 1-figure supplement 1). Although we have representative groups from all four major African linguistic macro-families (Supplementary File 1), our sample represents a significant proportion of the sub-Saharan population in terms of number, but not does not equate to a complete picture of African ethnic diversity.

To study patterns of genetic diversity we created an integrated dataset at a common set of over 328,000 high-quality SNP genotypes and use established approaches for comparing population allele frequencies across groups to provide a baseline view of historical gene flow. We apply statistical approaches to phasing genotypes to obtain haplotypes for each individual, and use previously published methods to represent the haplotypes that an individual carries as a mosaic of other haplotypes in the sample (so-called chromosome painting [Li and Stephens, 2003]). We use these data to demonstrate that haplotype-based methods have the potential to tease apart subtle relationships between closely related populations. We present a detailed picture of haplotype sharing across sub-Saharan Africa using a model-based clustering approach that groups individuals using haplotype information alone. The inferred groups reflect broad-scale geographic patterns. At finer scales, our analysis reveals smaller groups, and often differentiates closely related populations consistent with self-reported ancestry [Tishkoff et al., 2009; Bryc et al., 2010; Hodgson et al., 2014a]. We describe these patterns by measuring gene flow between populations and relate them to potential historical movements of people into and within sub-Saharan Africa. Understanding the extent to which individuals share haplotypes (which we call coancestry), rather than independent markers, can provide a rich description of ancestral relationships and population history [Lawson et al., 2012; Leslie et al., 2015]. For each group we use the latest analytical tools to characterise the populations as mixtures of haplotypes and provide estimates for the date of admixture events [Lawson et al., 2012; Hellenthal et al., 2014; Leslie et al., 2015; Montinaro et al., 2015]. As well as providing a quantitative measure of the coancestry between groups, we identify the detailed dominant events which have shaped current genetic diversity in sub-Saharan Africa. We discuss the relevance of these observations to studying genotype-phenotype associations in Africa, particularly in the context of infectious disease.

## Results

### Broad-scale population structure reflects geography and language

Throughout this article we use shorthand current-day geographical and ethno-linguistic labels to describe ancestry. For example we write “Eurasian ancestry in East African Niger-Congo speakers”, where the more precise definition would be “ancestry originating from groups currently living in Eurasia in groups currently living in East Africa that speak Niger-Congo languages” [Pickrell et al., 2014]. We also stress that the use of Khoesan in the current setting refers to groups with shared linguistic characteristics which does not necessarily imply shared close genealogical relationships [Güuldemann and Fehn, 2014]. Our combined dataset included 3,283 individuals from 46 sub-Saharan African ethnic groups and 12 non-African populations (Figure 1A and Figure 1-figure supplement 1).

**Figure 1.**
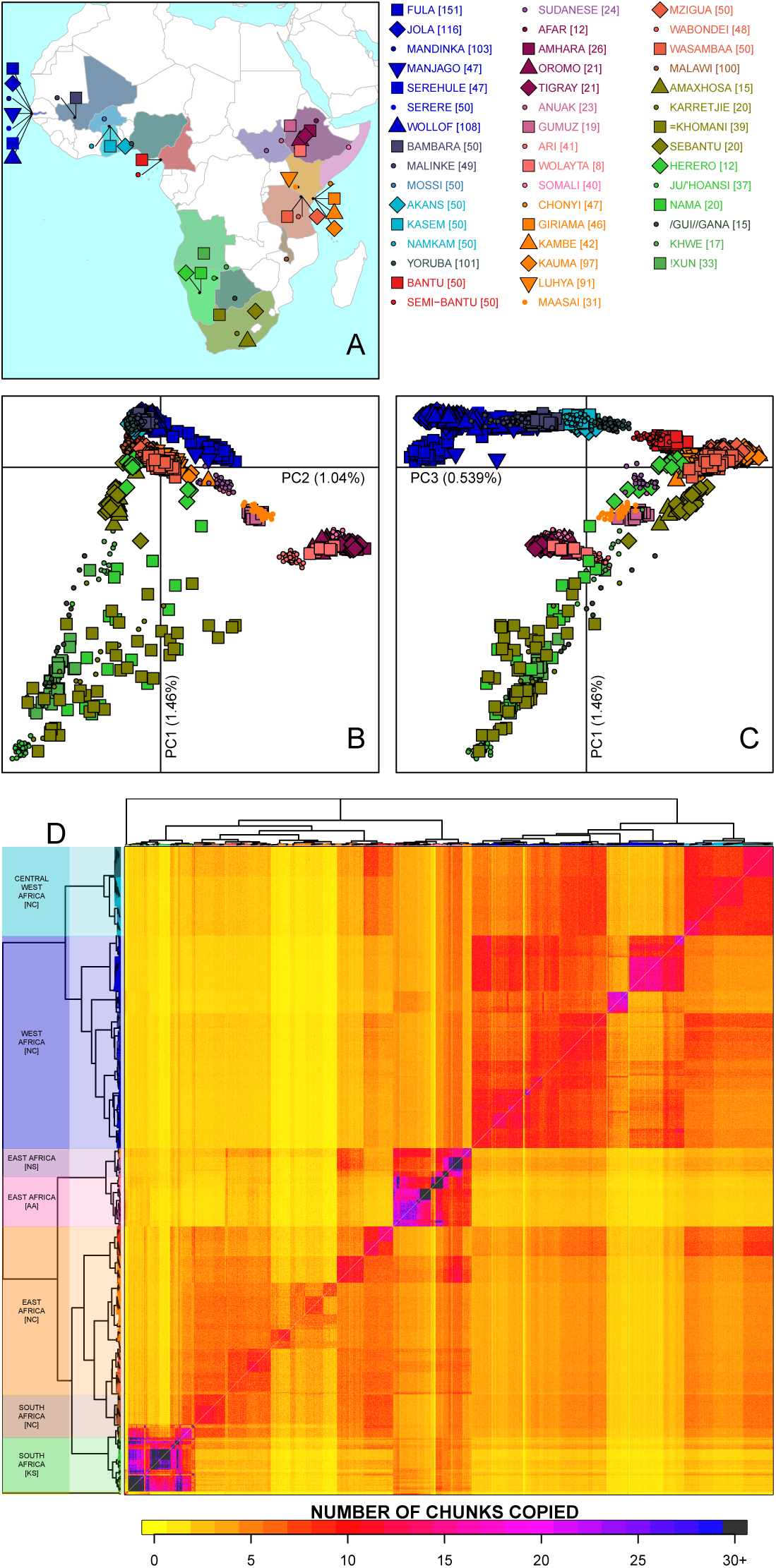
Sub-Saharan African genetic variation is shaped by ethno-linguistic and geographical similarity. (A) the origin of the 46 African ethnic groups used in the analysis; ethnic groups from the same country are given the same colour, but different shapes; the legend describes the identity of each point. Figure 1-figure supplement 1 and Figure 1-Source Data 1 provide further detail on the provenance of these samples. (B) PC1vPC2 shows that the first major axis of variation in Africa splits southern groups from the rest of Africa, each symbol represents an individual; PC2 reflects ethno-linguistic differences, with Niger-Congo speakers split from Afroasiatic and Nilo-Saharan speakers. (C) PC1vPC3 shows that the third principle component represents geographical separation of Niger-Congo speakers, forming a cline from west to east Africans. (D) results of the fineSTRUCTURE clustering analysis using copying vectors generated from chromosome painting; each row of the heatmap is a recipient copying vector showing the number of chunks shared between the recipient and every individual as a donor (columns);the tree clusters individuals with similar copying vectors together, such that block-like patterns are observed on the heat map; darker colours on the heatmap represent more haplotype sharing (see text for details); individual tips of the tree are coloured by country of origin, and the seven ancestry regions are identified and labelled to the left of the tree; labels in parentheses describe the major linguistic type of the ethnic groups within: AA = Afroasiatic, KS = Khoesan, NC = Niger-Congo, NS = Nilo-Saharan.

As in other regions of the world [Novembre et al., 2008; Reich et al., 2009; Behar et al., 2010], analyses of population structure using principal component analysis (PCA) show that genetic relationships are broadly defined by geographical and ethno-linguistic similarity (Figure 1B,C). The first two principal components (PCs) reflect ethno-linguistic divides: PC1 splits southern Khoesan speaking populations from the rest of Africa, and PC2 splits the Afroasiatic and Nilo-Saharan speakers from East Africa from sub-Saharan African Niger-Congo speakers. The third axis of variation defines east versus west Africa, suggesting that in general, population structure in Africa largely mirrors linguistic and geographic similarity [Tishkoff et al., 2009].

Using a previously published implementation of chromosome painting (CHROMOPAINTER [Lawson et al., 2012]), we reconstructed the genomes of each study individual as mosaics of haplotype segments (or chunks) from all other individuals. We used the clustering algorithm implemented in fineSTRUCTURE [Lawson et al., 2012] to cluster individuals into groups based purely on the similarity of these paintings (Figure 1 and Figure 1-figure supplement 3). More specifically, for a given recipient individual, we can estimate the amount of their genome that is shared with (or copied from) each of a set of donor individuals based on the paintings, which are summarised as copying vectors. These vectors are clustered hierarchically to form a tree which describes the inferred relationship between different groups (Figure 1-figure supplement 3). As such, this method uses chromosome painting to describe the coancestry between groups, which can then be visualised as a heatmap, as in Figure 1D.

Consistent with PCA, African populations tend to share more DNA with geographically proximate populations (dark colours on the diagonal; Figure 1D). Block structures on this diagonal indicate higher levels of haplotype sharing within groups, which is indicative of close genealogical relationships with other members of your group. These patterns can be seen in some of the Khoesan speaking individuals (eg. the Ju/’hoansi), several groups from the East Africa (Sudanese, Ari, and Somali groups), and the Fulani and Jola from the Gambia, each of which provides clues to the ancestral connections between the groups in our analysis. The heatmap also shows evidence for coancestry across regions (dark colours away from the diagonal), which can be informative of historical connections between modern-day groups. For example, east Africans from Kenya, Malawi and Tanzania tend to share more DNA with west Africans (lower right) which suggests that more haplotypes may have spread from west to east Africa (upper left) than vice versa. Using the results of the PCA and fineSTRUCTURE analyses together with ethno-linguistic classifications and geography, we defined seven groups of populations within Africa (Supplementary File 1), which we refer to as ancestry regions when describing gene-flow across Africa. Under this definition, there is widespread sharing of haplotypes within and between ancestry regions (Figure 1).

### Haplotypes reveal subtle population structure

To quantify the extent of the genetic difference between groups we used two different metrics. First, we used the classical measure *F_ST_* [Hudson et al., 1992; Bhatia et al., 2013] which measures the differentiation in allele frequencies between populations. It can be thought of as measuring the proportion of the heterozygosity at SNPs explained by the group labels. The second metric uses the similarity in copying patterns between two groups to estimate the total variation difference (TVD) at the haplotypic level. TVD takes advantage of the fact that recombination rates are faster than mutation rates, and so is expected to capture more variability than genotype counts. Figure 2A shows these two metrics side by side in the upper and lower diagonal. When compared to non-African populations, *F_ST_* measured at our integrated set of SNPs is relatively low between many groups from West, Central, and East Africa (yellows on the upper right triangle), whereas TVD in the same populations can reveal haplotypic differences as strong as between Europe and Asia (pink and purples in lower left triangle). For example, the Chonyi from Kenya have relatively low *F_ST_* but high TVD with West African groups, like the Jola (Chonyi-Jola *F_ST_* = 0.019; Chonyi-Jola TVD = 0.803) suggesting that, whilst allele frequency differences between the two populations are relatively low, when we compare the populations’ ancestry vectors, the haplotypic differences are some the strongest between sub-Saharan groups. In fact, whilst pairwise TVD tends to increase with pairwise *F_ST_* (Pearson’s correlation *R*^2^ = 0.79) the relationship is not linear (Figure 2B) demonstrating that this discrepancy is not simply a function of TVD generating larger statistics. We note that high values of TVD are not sufficient to infer specific demographic events [van Dorp et al., 2015] and we therefore do not interpret these differences as necessarily resulting from admixture, but as motivation to use haplotype-based methods to characterise population relationships.

**Figure 2.**
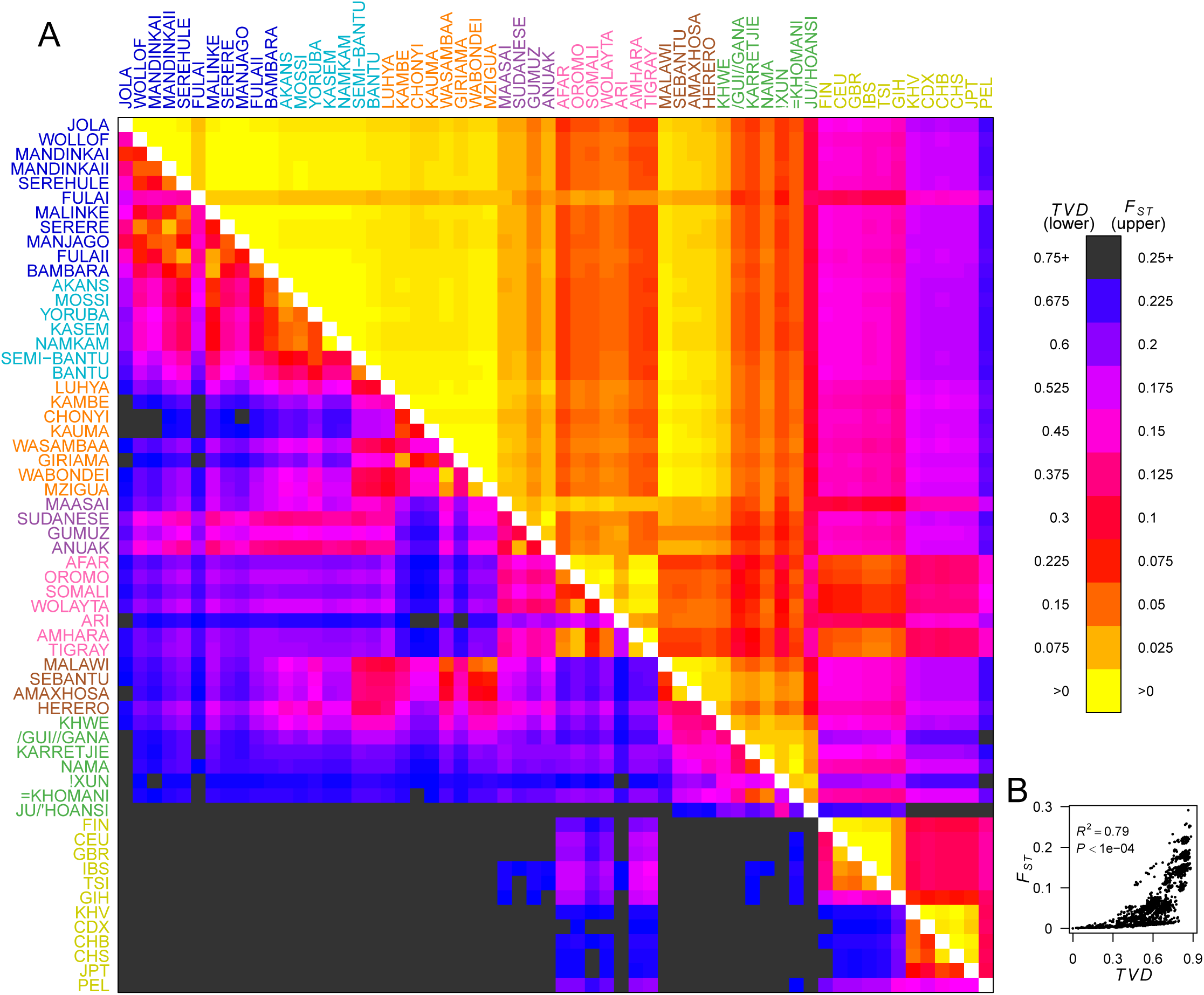
Haplotypes capture more population structure than independent loci. (A) For each population pair, we estimated pairwise *F_ST_* (upper right triangle) using 328,000 independent SNPs, and *TV D* (lower left triangle) using population averaged copying vectors from CHROMOPAINTER. *TV D* measures the difference between two copying vectors. (B) Comparison of pairwise *F_ST_* and *TV D* shows that they are not linearly related: some population pairs have low *F_ST_* and high *TV D*. [Source data is detailed in Figure 2-Source Data 1 to Figure 2-Source Data 4]

### Widespread evidence for admixture

To check that our dataset was consistent with those used in other analyses of African variation, we next applied similar approaches to previous authors to infer admixture between populations using correlations in allele frequencies within and between populations [Pickrell et al., 2014; Gurdasani et al., 2014]. The first approach, the three-population test (*f*_3_ statistic [Reich et al., 2009]), uses correlations in allele frequencies between a target population and two potential source populations to identify significant departures from the null model of no admixture. Negative values are indicative of canonical admixture events where the allele frequencies in the target population are intermediate between the two source populations. Consistent with recent research [Pickrell et al., 2014; Pickrell and Reich, 2014; Gurdasani et al., 2014; Llorente et al., 2015], the majority (80%, 40/48), but not all, of the African groups surveyed showed evidence of admixture (*f*_3_<-5), with the most negative *f*_3_ statistic tending to involve either a Eurasian or African hunter-gatherer group [Gurdasani et al., 2014] (Supplementary Table 1).

The second approach, ALDER [Loh et al., 2013; Pickrell et al., 2014] (Supplementary Table 1) exploits the fact that correlations between allele frequencies along the genome decay over time as a result of recombination. Linkage disequilibrium (LD) can be generated by admixture events, and leaves detectable signals in the genome that can be used to infer historical processes from allele frequency data [Loh et al., 2013]. Following Pickrell et al. [2014] and the AGVP [Gurdasani et al., 2014], we computed weighted admixture LD curves using the ALDER [Loh et al., 2013] package to characterise the sources and timing of gene flow events. Specifically, we estimated the y-axis intercept (amplitude) of weighted LD curves for each target population using curves from an analysis where one of the sources was the target population (self reference) and the other was, separately, each of the other (non-self reference) populations. Theory predicts that the amplitude of these “one-reference” curves becomes larger the more similar the non-self reference population is to the true admixing source [Loh et al., 2013]. As with the *f*_3_ analysis outlined above, for many of the sub-Saharan African populations, Eurasian and hunter-gatherer groups produced the largest amplitudes (Figure 3-figure supplement 1 and Figure 3-figure supplement 2), reinforcing the contribution of these ancestries to our broad set of African populations.

We investigated the evidence for more complex admixture using MALDER [Pickrell et al., 2014], an implementation of ALDER which fits a mixture of exponentials to weighted LD curves to infer multiple admixture events (Figure 3 and Table Figure 3-Source Data 1). In Figure 3A, for each target population, we show the ancestry region of the two populations involved in generating the MALDER curves with the greatest amplitudes, together with the date of admixture for at most two events. Throughout, we convert time since admixture in generations to a date by assuming a generation time of 29 years [Fenner, 2005]. In general, we find that groups from similar ancestry regions tend to have inferred events at similar times and between similar groups (Figure 3), which suggests that genetic variation in groups has been shaped by shared historical events. For every event, the curves with the greatest amplitudes involved a population from a (normally non-Khoesan) African population on the one side, and either a Eurasia or Khoesan population on the other. To provide more detail on the composition of the admixture sources, we compared MALDER curve amplitudes between curves involving populations from different ancestry regions (central panel Figure 3). In general, this analysis showed that, with a few exceptions, we were unable to precisely define the ancestry of the African source of admixture, as curves involving populations from multiple different regions were not statistically different from each other (*Z* < 2; SOURCE 1). Conversely, comparisons of MALDER curves when the second source of admixture was Eurasian (yellow) or Khoesan (green), showed that these groups were ususally the single best surrogate for the second source of admixture (SOURCE 2).

**Figure 3.**
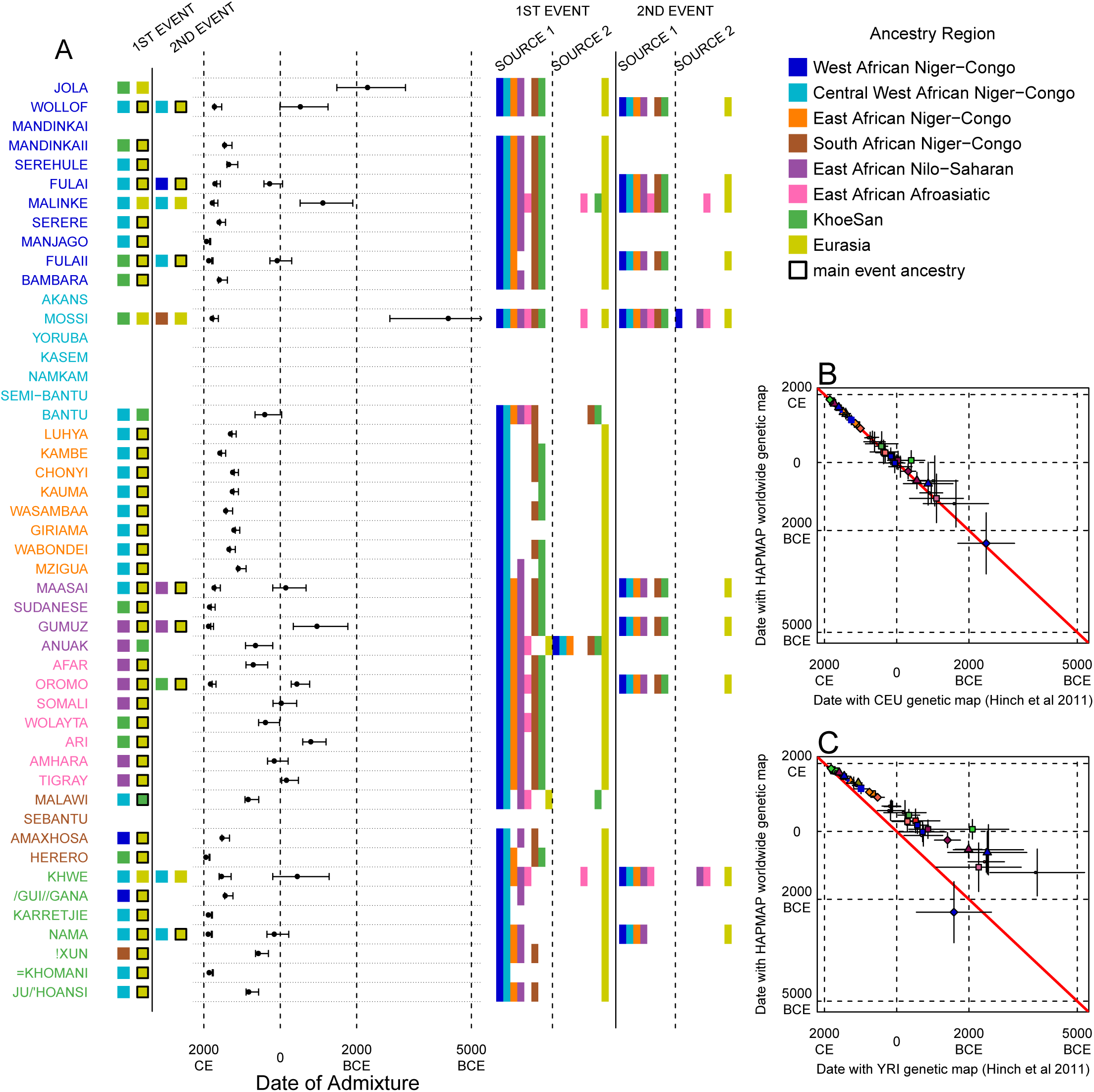
Inference of admixture in sub-Saharan Africa using MALDER. We used MALDER to identify the evidence for multiple waves of admixture in each population. (A) For each population, we show the ancestry region identity of the two populations involved in generating the MALDER curves with the greatest amplitudes (coloured blocks) for at most two events. The major contributing sources are highlighted with a black box. Populations are ordered by ancestry of the admixture sources and dates estimates which are shown ± 1 s.e. For each event we compared the MALDER curves with the greatest amplitude to other curves involving populations from different ancestry regions. In the central panel, for each source, we highlight the ancestry regions providing curves that are not significantly different from the best curves. In the Jola, for example, this analysis shows that, although the curve with the greatest amplitude is given by Khoesan (green) and Eurasian (yellow) populations, curves containing populations from any other African group (apart from Afroasiatic) in place of a Khoesan population are not significantly smaller than this best curve (SOURCE 1). Conversely, when comparing curves where a Eurasian population is substituted with a population from another group, all curve amplitudes are significantly smaller (*Z* < 2). (B) Comparison of dates of admixture ± 1 s.e. for MALDER dates inferred using the HAPMAP recombination map and a recombination map inferred from European (CEU) individuals from [Hinch et al., 2011]. We only show comparisons for dates where the same number of events were inferred using both methods. Point symbols refer to populations and are as in Figure 1. (C) as (B) but comparison uses an African (YRI) map. Source data can be found in Figure 3-Source Data 1

We performed multiple MALDER analyses, varying the input parameters and the genetic map used. We observed a large amount of shared LD at short genetic distances between different African populations (Figure 3-figure supplement 3 and Figure 3-figure supplement 4). Such patterns may result from population genetic processes other than admixture, such as shared demographic history and population bottlenecks [Loh et al., 2013]. In the main MALDER analysis we present, we have removed the effect of this short-range LD by generating curves only after ignoring SNPs at short genetic distances where frequencies of markers at such distances are correlated between target and reference population – which can be thought of as conservative – but provide supplementary analyses where this parameter was relaxed. The main difference between the two types of analysis is that we were only able to identify ancient events in West Africa if we force the MALDER algorithm to start computing LD decay curves from a genetic distance of 0.5cM, irrespective of any short-range correlations in LD between populations.

Many West African groups show evidence of recent (within the last 4 ky) admixture involving African and Eurasian sources. The Mossi from Burkina Faso have the oldest inferred date of admixture, at roughly 5000BCE, but we were unable to recreate previously reported ancient admixture events in other Central West African groups [Gurdasani et al., 2014] (although see Figure 3-figure supplement 5 for results where we use different MALDER parameters). Across East Africa Niger-Congo speakers (orange) we infer admixture within the last 4 ky (and often within the last 1 ky) involving Eurasian sources on the one hand, and African sources containing ancestry from other Niger-Congo speaking African groups from the west on the other. Despite events between African and Eurasian sources appearing older in the Nilo-Saharan and Afroasiatic speakers from East Africa, we see a similar signal of very recent Central West African ancestry in a number of Khoesan groups from Southern Africa, such as the Khwe, /Gui //Gana, and !Xun, together with Malawi-like (brown) sources of ancestry in recent admixture events in East African Niger-Congo speakers.

Inference of older events relies on modelling the decay of LD over short genetic distances because recombination has had more time to break down correlations in allele frequencies between neighbouring SNPs. We investigated the effect of using European (CEU) and Central West African (YRI) specific recombination maps [Hinch et al., 2011] on the dating inference. Whilst dates inferred using the CEU map were consistent with those using the HAPMAP recombination map (Figure 3B), when using the African map dates were consistently older (Figure 3C), although still generally still within the last 7ky. There was also variability in the number of inferred admixture events for some populations between the different map analyses (Figure 3-figure supplement 6 and Figure 3-figure supplement 7).

Most events involved sources where Eurasian (dark yellow in Figure 3A) groups gave the largest amplitudes. In considering this observation, it is important to note that the amplitude of LD curves will partly be determined by the extent to which a reference population has differentiated from the target. Due to the genetic drift associated with the out-of-Africa bottleneck and subsequent expansion, Eurasian groups will tend to generate the largest curve amplitudes even if the proportion of this ancestry in the true admixing source is small [Pickrell et al., 2014] (in our dataset, the mean pairwise *F_ST_* between Eurasian and African populations is 0.157; Figure 2A and Figure 2-Source Data 1). To some extent this also applies to Khoesan groups (green in Figure 3A), who are also relatively differentiated from other African groups (mean pairwise *F_ST_* between Ju/’hoansi and all other African populations in our dataset is 0.095; Figure 2A and Figure 2-Source Data 1). In light of this, and the observation that curves involving groups from different ancestry regions are often no different from each other, it is therefore difficult to infer the proportion or nature of the African, Khoesan, or Eurasian admixing sources, only that the sources themselves contained African, Khoesan, or Eurasian ancestry. Moreover, given uncertainty in the dating of admixture when using different maps and MALDER parameters, these results should be taken as a guide to the general genealogical relationships between African groups, rather than a precise description of the gene-flow events that have shaped Africa.

### Modelling gene flow with haplotypes

So far, our analyses have largely recapitulated recent studies of the genetic history of African populations, albeit across a broader set of sub-Saharan populations. However, we were interested in gaining a more detailed characterisation of historical gene-flow to provide an understanding and framework with which to inform genetic epidemiological studies in Africa. The chromosome painting methodology described above provides an alternative approach to inferring admixture events which directly models the similarity in haplotypes between pairs of individuals. Evidence of recent haplotype sharing suggests that the ancestors of two individuals must have been geographically proximal at some point in the past. More generally, the distance over which haplotype sharing extends along chromosomes is inverse to how far in the past coancestry events have occurred. We can use copying vectors inferred through chromosome painting to help identify those populations that share ancestry with a recipient group by fitting each vector as a mixture of all other population vectors (Figure 4A) [Leslie et al., 2015; Montinaro et al., 2015; van Dorp et al., 2015]. Figure 4A shows the contribution that each ancestry region makes to these mixtures (MIXTURE MODEL column). Almost all groups can best be described as mixtures of ancestry from different regions. For example, the copying vector of the Bantu ethnic group from Cameroon is best described as a combination of 40% Central West African Niger-Congo (sky blue), 30% Eastern Niger-Congo (orange), 25% Southern Niger-Congo (brown), and the remaining 5% coming from West African Niger-Congo (dark blue) and Khoesan-speaking (green) groups. The key insight from this analysis is that many African groups share fragments of haplotypes with groups from outside of their own ancestry region.

**Figure 4.**
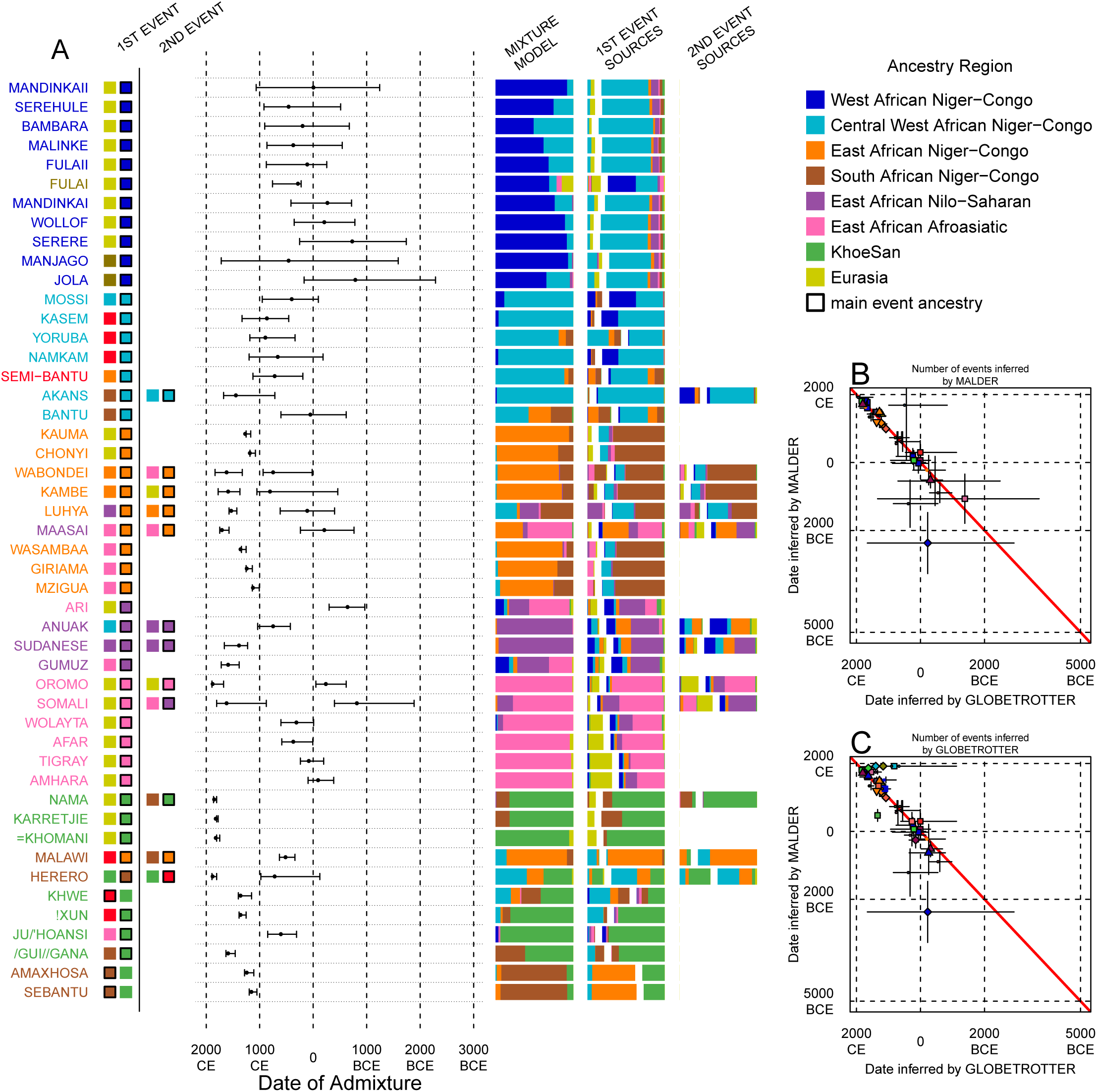
Inference of admixture in sub-Saharan African using GLOBETROTTER. (A) For each group we show the ancestry region identity of the best matching source for the first and, if applicable, second events. Events involving sources that most closely match FULAI and SEMI-BANTU are highlighted by golden and red colours, respectively. Second events can be either multiway, in which case there is a single date estimate, or two-date in which case the 2ND EVENT refers to the earlier event. The point estimate of the admixture date is shown as a black point, with 95% CI shown with lines. MIXTURE MODEL: We infer the ancestry composition of each African group by fitting its copying vector as a mixture of all other population copying vectors. The coefficients of this regression sum to 1 and are coloured by broad ethno-linguistic origin. 1ST EVENT and 2ND EVENT SOURCES shows the ancestry breakdown of the admixture sources inferred by GLOBETROTTER, coloured by ancestry region as in the key top right. (B) and (C) Comparisons of dates inferred by MALDER and GLOBETROTTER. Because the two methods sometimes inferred different numbers of events, in (B) we show the comparison for based on the inferred number of events in the MALDER analysis, and in (C) for the number of events inferred by GLOBETROTTER. Point symbols refer to populations and are as in Figure 1 and source data can be found in Figure 4-Source Data 1.

The mixture model approach is useful for describing and summarising the copying vectors of populations, but such summaries can result from both admixture and shared evolutionary history. We thus used GLOBETROTTER [Hellenthal et al., 2014], a method that explicitly tests for and characterises admixture as an extension of the mixture model approach described above, to gain a more detailed understanding of recent ancestry. Admixture inference can be challenging for a number of reasons: the true admixing source population is often not well represented by a single sampled population; admixture could have occurred in several bursts, or over a sustained period of time; and multiple groups may have come together as complex convolution of admixture events. GLOBETROTTER aims to overcome some of these challenges, in part by using painted chromosomes to explicitly model the correlation structure among nearby SNPs, but also by allowing the sources of admixture themselves to be mixed [Hellenthal et al., 2014]. In addition, the approach has been shown to be relatively insensitive to the genetic map used [Hellenthal et al., 2014], and therefore potentially provides a more robust inference of admixture events, the ancestries involved, and their dates. GLOBETROTTER uses the distance between chromosomal chunks of the same ancestry to infer the time since historical admixture has occurred.

Throughout the following discussion, we refer to target populations as *recipients*, any other sampled populations used to describe the recipient population’s admixture event(s) as surrogates, and populations used to paint both target and surrogate populations as donors. Including closely related individuals in chromosome painting analyses can cause the resulting painted chromosomes to be dominated by donors from these close genealogical relationships, which can mask signals of admixture in the genome [Hellenthal et al., 2014; van Dorp et al., 2015]. To help ameliorate this, we painted chromosomes for the GLOBETROTTER analysis by performing a fresh run of CHROMOPAINTER where we painted each individual from a recipient group with a set of donors which did not include individuals from within their own ancestry region. We additionally painted all (59) other surrogate populations with the same set of non-local donors, and used these copying vectors, together with the non-local painted chromosomes, to infer admixture. Using this approach, we found evidence of recent admixture in all African populations (Figure 4A). To summarise these events, we show the composition of the admixing source groups as barplots for each population coloured by the contribution from each African ancestry regions and Eurasia, alongside the inferred date with confidence interval determined by bootstrapping, and the estimated proportion of admixture (Figure 4). For each event we also identify the best matching donor population to the admixture sources. We describe observations from these analyses below, and note that the dates of admixture we infer indicate when gene-flow occurred between source populations and not the arrival of groups into an area, which may often be several generations earlier.

### Direct and indirect gene flow from Eurasia back into Africa

We did not find evidence for recent Eurasian admixture in every African population (Figure 4). In particular, in several groups from South Africa and all from the Central West African ancestry region, which includes populations from Ghana, Nigeria, and Cameroon, we infer admixture between groups that are best represented by contemporary populations residing in Africa. As GLOBETROTTER is designed to identify the most recent admixture event(s) [Hellenthal et al., 2014], this observation does not rule out gene-flow from Eurasia back into these groups, but does suggest that subsequent movements between African groups were important in generating the contemporary ancestry of Central West and Southern African Niger-Congo speaking groups. With some exceptions that we describe below, we also do not observe Eurasian ancestry in all East African Niger-Congo speakers, instead finding more evidence for coancestry with Afroasiatic speaking groups. As we show later, Afroasiatic populations have a significant amount of genetic ancestry from outside of Africa, so the observation of this ancestry in several African groups identifies a route by which Eurasian ancestry may have indirectly entered the continent [Pickrell et al., 2014].

In fact, characterising admixture sources as mixtures allows us to infer whether Eurasian haplotypes are likely to have come directly into sub-Saharan Africa – in which case the admixture source will contain only Eurasian surrogates – or whether Eurasian haplotypes were brought indirectly together with sub-Saharan groups. In West African Niger-Congo speakers from The Gambia and Mali, we infer admixture involving minor admixture sources which contain mostly Eurasian (dark yellow) and Central West African (sky blue) ancestry, which most closely match the contemporary copying vectors of northern European populations (CEU and GBR) or the Fulani (FULAI, highlighted in gold in Figure 4A). The Fulani, a nomadic pastoralist group found across West Africa, were sampled in The Gambia, at the very western edge of their current range, and have previously reported genetic affinities with Niger-Congo speaking, Sudanic, Saharan, and Eurasian populations [Tishkoff et al., 2009; Henn et al., 2012], consistent with the results of our mixture model analysis (Figure 4A). Admixture in the Fulani differs from other populations from this region, with sources containing greater amounts of Eurasian and Afroasiatic ancestry, but appears to have occurred during roughly the same period (c. 0CE; Figure 5).

**Figure 5.**
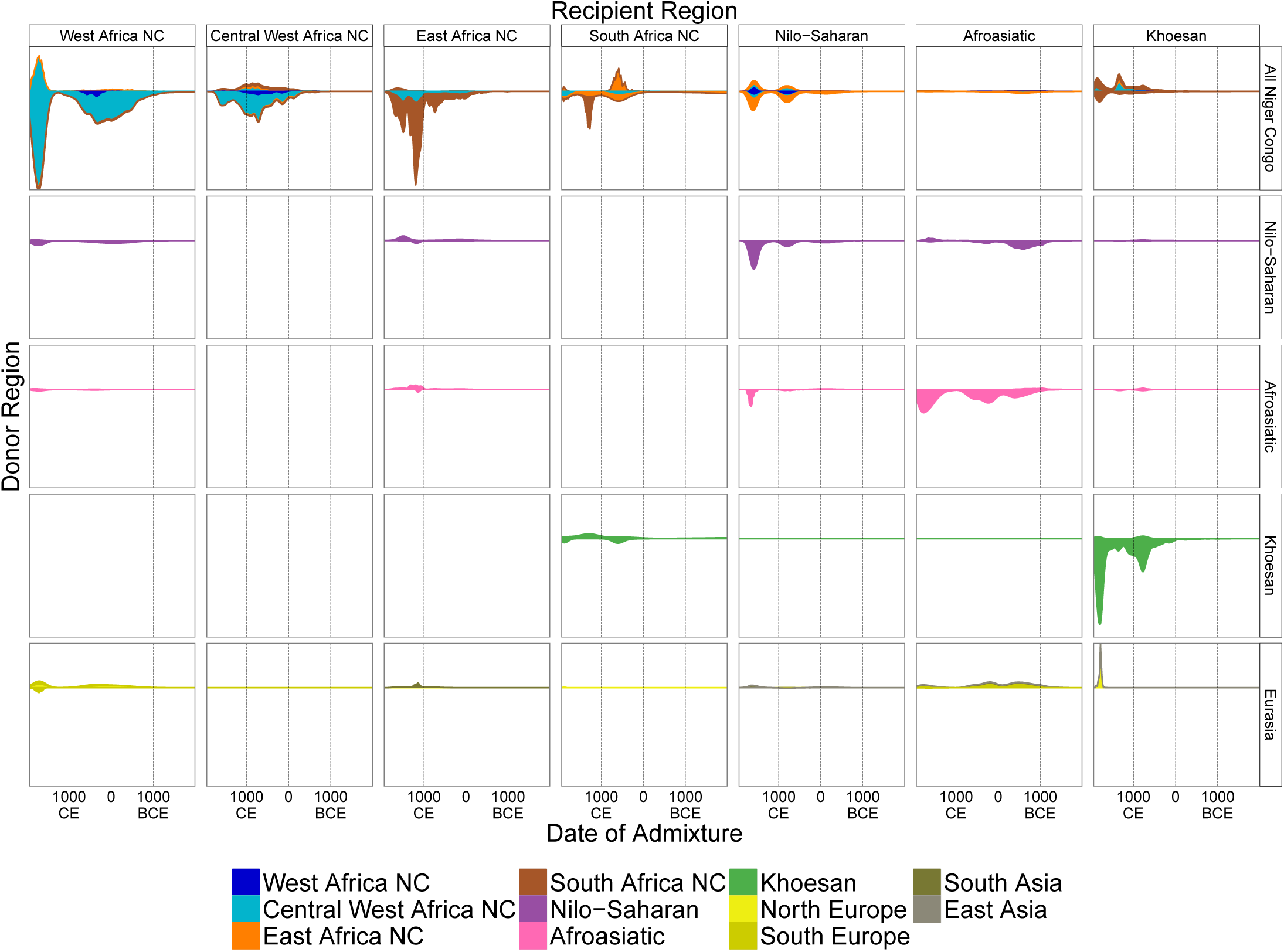
A timeline of recent admixture in sub-Saharan Africa. For all events involving recipient groups from each ancestry region (columns) we combine all date bootstrap estimates generated by GLOBETROTTER and show the densities of these dates separately for the minor (above line) and major (below line) sources of admixture. Dates are additionally stratified by the ancestry region of the surrogate populations (rows), with all dates involving Niger Congo speaking regions combined together (All Niger Congo). Within each panel, the densities are coloured by the geographic (country) origin of the surrogates and in proportion to the components of admixture involved in the admixture event. The integrals of the densities are proportional to the admixture proportions of the events contributing to them.

The Fulani represent the best-matching surrogate to the minor source of recent admixture in the Jola and Manjago, which we interpret as resulting not from specific admixture from them into these groups, but because the mix of African and Eurasian ancestries in contemporary Fulani is the best proxy for the minor sources of admixture in this region. With the exception of the Fulani themselves, the major admixture source in groups across this region is a similar mixture of African ancestries that most closely matches contemporary Gambian and Malian surrogates (Jola, Serere, Serehule, and Malinke), suggesting ancestry from a common West African group within the last 3,000 years. The Ghana Empire flourished in West Africa between 300 and 1200CE, and is one of the earliest recorded African states [Roberts, 2007]. Whilst its origins are uncertain, it is clear that trade in gold, salt, and slaves across the Sahara, perhaps from as early as the Roman Period, as well as evolving agricultural technologies, were the driving forces behind its development [Oliver and Fagan, 1975; Roberts, 2007]. The observation of Eurasian admixture in our analysis is at least consistent with this moment in African history, and suggests that ancestry in groups from across this region of West Africa is the result of interactions through North Africa that were catalysed by trade across the Sahara.

We infer direct admixture from Eurasian sources in two populations from Kenya, where specifically South Asian populations (GIH, KHV) are the most closely matched surrogates to the minor sources of admixture (Figure 5). Interestingly, the Chonyi (1138CE: 1080-1182CE) and Kauma (1225CE: 1167-1254CE) are located on the so-called Swahili Coast, a region where Medieval trade across the Indian Ocean is historically documented [Allen, 1993]. In the Kambe, the third group from coastal Kenya, we infer two events, the more recent one involving local groups, and the earlier event involving a European-like source (GBR, 761CE: 461BCE-1053CE). In Tanzanian groups from the same ancestry region, we infer admixture during the same period, this time involving minor admixture sources with different, Afroasiatic ancestry: in the Giriama (1196CE: 1138-1254CE), Wasambaa (1312CE: 1254-1341CE), and Mzigua (1080: 1007-1138CE). Although the proportions of admixture from these sources differ, when we consider the major sources of admixture in East African Niger-Congo speakers, they are again similar, containing a mix of mainly local Southern Niger-Congo (Malawi), Central West African, Afroasiatic, and Nilo-Saharan ancestries.

In the Afroasiatic speaking populations of East Africa we infer admixture involving sources containing mostly Eurasian ancestry, which most closely matches the Tuscans (TSI, Figure 4). This ancestry appears to have entered the Horn of Africa in three distinct waves (Figure 5). We infer admixture involving Eurasian sources in the Afar (326CE: 7-587CE), Wolayta (268CE: 8BCE-602CE), Tigray (36CE: 196BCE-240CE), and Ari (689BCE:965-297BCE). There are no Middle Eastern groups in our analysis, and this latter group of events may represent previously observed migrations from the Arabian peninsular from the same time [Pagani et al., 2012; Hodgson et al., 2014a]. Considering Afroasiatic and Nilo-Saharan speakers separately, the ancestry of the major sources of admixture of the former are predominantly local (purple), indicative of less historical interaction with Niger-Congo speakers due to their previously reported Middle Eastern ancestry [Pagani et al., 2012]. In Nilo-Saharan speaking groups, the Sudanese (1341CE: 1225-1660), Gumuz (1544CE: 1384-1718), Anuak (703: 427-1037CE), and Maasai (1646CE: 1584-1743CE), we infer greater proportions of West (blue) and East (orange) African Niger-Congo speaking surrogates in the major sources of admixture, indicating both that the Eurasian admixture occurred into groups with mixed Niger-Congo and Nilo-Saharan / Afroasiatic ancestry, and a clear recent link with Central and West African groups.

In two Khoesan speaking groups from South Africa, the ≠Khomani and Karretjie, we infer very recent direct admixture involving Eurasian groups most similar to Northern European populations, with dates aligning to European colonial period settlement in Southern Africa (c. 5 generations or 225 years ago; Figure 5) [Hellenthal et al., 2014]. Taken together, and in addition the MALDER analysis above, these observations suggest that gene flow back into Africa from Eurasia has been common around the edges of the continent, has been sustained over the last 3,000 years, and can often be attributed to specific and different historical time periods.

### Population movements within Africa and the Bantu expansion

Admixture events involving sources that best match populations from within Africa tend to involve local groups. Even so, there are several long range admixture events of note. In the Ju/’hoansi, a San group from Namibia, we infer admixture involving a source that closely matches a local southern African Khoesan group, the Karretjie, and an East African Afroasiatic, specifically Somali, source at 558CE (311-851CE). We infer further events with minor sources most similar to present day Afroasiatic speakers in the Maasai (1660CE: 1573-1747CE and 254BCE: 764-239BCE) and, as mentioned, all four Tanzanian populations (Wabondei, Wasambaa, Giriama, and Mzigua). In contrast, the recent event inferred in the Luhya (1486: 1428-1573CE) involves a Nilo-Saharan-like minor source. The recent dates of these events imply not only that Eastern Niger-Congo speaking groups have been interacting with nearby Nilo-Saharan and Afroasiatic speakers – who themselves have ancestry from more northerly regions – after the putative arrival of Bantu-speaking groups to Eastern Africa, but also that the major sources of admixture contained both Central West and a majority of Southern Niger-Congo ancestry.

With the exception of Austronesian in Madagascar, African languages can be broadly classified into four major macro-families: Afroasiatic, Nilo-Saharan, Niger-Congo, and Khoesan [Blench, 2006]. Most of the sampled groups in this study, and indeed most sub-Saharan Africans, speak a language belonging to the Niger Congo lingustic phylum [Greenberg, 1972; Nurse and Philippson, 2003]. A sub-branch of this group are the so-called “Bantu” languages – a group of approximately 500 very closely related languages – that are of particular interest because they are spoken by the vast majority of Africans south of the line between Southern Nigeria/Cameroon and Somalia [Pakendorf et al., 2011]. Given the their high similarity and broad geographic range, it is likely that Bantu languages spread across Africa quickly. Whether this cultural expansion was accompanied by people is an active research question, but an increasing number of molecular studies, mostly using uni-parental genetic markers, indicate that the expanson of languages was accompanied by the diffusion of people [Beleza et al., 2005; Berniell-Lee et al., 2009; Pakendorf et al., 2011; de Filippo et al., 2012; Ansari Pour et al., 2013; Li et al., 2014; González-Santos et al., 2015]. Bantu languages can themselves be divided into three major groups: northwestern, which are spoken by groups near to the proto-Bantu heartland of Nigeria / Cameroon; western Bantu languages, spoken by groups situated down the west coast of Africa; and eastern, which are spoken across East and Central Africa [Li et al., 2014]. One particular debate concerns whether eastern languages are a primary branch that split off before the western groups began to spread south (the early-split hypothesis) or whether this occurred after the migrants had begun to spread south (the late-split hypothesis) [Pakendorf et al., 2011]. Recent linguistic [Holden, 2002; Currie et al., 2013; Grollemund et al., 2015], and genetic analyses [Li et al., 2014] support the latter.

Whilst the current dataset does not cover all of Africa, and importantly contains no hunter-gather groups outside of southern Africa, we explored whether our admixture approach could be used to gain insight into the Bantu expansion. Specifically, we wanted to see whether the dates of admixture and composition of admixture sources were consistent with either of the two major models of the Bantu expansion. The major sources of admixture in East African Niger-Congo speakers tend to have both Central West and Southern Niger-Congo ancestry, although it is predominantly the latter (Figure 4). If Eastern Niger-Congo speakers derived all of their ancestry directly from Central West Africa (i.e. the early-split hypothesis) then we would not observe any ancestry from more southerly groups, but this is not what we see. That all East African Niger-Congo speakers that we sampled have admixture ancestry from a Southern group (Malawi), suggests that this group is more closely related to their Bantu ancestors than Central West Africans on their own. To explore this further, we performed additional GLOBETROTTER analyses where we restricted the surrogate populations used to infer admixture. For example, we re-ran the admixture inference process without allowing any groups from within the same ancestry region to contribute to the admixture event. As in the main analysis, when we disallow any East African Niger-Congo speakers from forming the admixture, Southern African Niger-Congo speakers, specifically Malawi, contribute most to the major source of admixture. In Southern African Niger-Congo groups, restricting admixture surrogates to those outside of their region leads to East African groups contributing most to the sources of admixture. The closer affinity between East and Southern Niger-Congo Bantu components is consistent with a common origin of these groups after the split from the Western Bantu. Moreover, this observation provides additional support for the hypothesis based on linguistic phylogenetics [Currie et al., 2013; Grollemund et al., 2015] that the main Bantu migration moved south and then east around the Congo rainforest. We performed a further restriction, disallowing any Southern or Eastern Niger-Congo speaking groups from being involved in admixture across both the South and East African Niger-Congo regions, which led to the vast majority of the ancestry of the major sources of admixture originating from Central West Africa, and almost exclusively from Cameroon populations (Figure 4-figure supplement 1 and Figure 4-figure supplement 2).

We inferred recent admixture involving major sources most similar to East or Southern African Niger-Congo speakers in the SEBantu (1109:1051-1196CE) and AmaXhosa (1196CE: 1109-1283CE) from South Africa. We infer two admixture dates in a Bantu-speaking group from Namibia, the Herero (1834CE: 1805-1892CE and 674CE: 124BCE-979CE), and a single date in the Khoesan-speaking Khwe (1312; 1152-1399CE), involving sources that more clearly contain ancestry from the Semi-Bantu. (Semi-Bantu and Bantu refer to two contemporary ethnic groups that currently reside in Cameroon.) In a third south-west African group, the !Xun from Angola, we infer admixture from a similar Cameroon-like source at around the same time as the Khwe (1312CE: 1254-1385CE). Assuming that Cameroon populations are the best proxy for Bantu ancestry in our dataset, these results suggest either a separate, more recent, arrival for Niger-Congo (Bantu) ancestry in south-west compared to south-east Africa, with the former coming recently directly down the west coast of Africa, specifically from Cameroon, and the latter deriving their ancestry from earlier interactions via an eastern route [de Filippo et al., 2012; Li et al., 2014], or that there was limited detectable gene-flow in south-east Africa from the Bantus that eventually moved into south-west Africa.

Interestingly, in individuals from Malawi we infer a multi-way event with an older date (471: 340-631CE) involving a minor source which mostly contains ancestry from Cameroon, an event that is also seen in the Herero from Namibia. This Bantu admixture appears to have preceded that in some of the other South Africans by a few hundred years, suggesting either input from both western and eastern migrating Bantu speakers after the two branches split c.1,500 years ago, or that present day Malawi harbour ancestry from groups intermediate between the two branches [de Filippo et al., 2012]. Within this group we also see an admixture source with a significant proportion of non-Bantu (green) ancestry (2nd event, minor source), ancestry which we do not observe in the mixture model analysis, but which is also evident in other south-east African Niger-Congo speakers, the AmaXhosa and SEBantu, indicating that gene-flow must have occurred between the expanding Bantus and the resident hunter-gatherer groups [Marks et al., 2014]. The complex ancestry composition of the sources of these events, together with the dates of admixture, suggest large-scale interactions between groups during the Bantu expansion, and also hint that the expansion may have generated multiple waves of movement south. These events have resulted in recent haplotype sharing between groups across sub-Saharan Africa.

### A haplotype-based model of gene flow in sub-Saharan Africa

Our haplotype-based analyses support a complex and dynamic picture of recent historical gene flow in Africa. We next attempted to summarise these events by producing a demographic model of African genetic history (Figure 6). Using genetics to infer historical demography will always depend somewhat on the available samples and population genetics methods used to infer population relationships. The model we present here is therefore unlikely to recapitulate the full history of the continent and will be refined in the future with additional data and methodology. We again caution that we do not have an exhaustive sample of African populations, and in particular lack significant representation from extant hunter-gatherer groups outside of southern Africa. Nevertheless, there are signals in the data that our haplotype-based approach allows us to pick up, and it is therefore possible to highlight the key gene-flow events and sources of coancestry in Africa, which we visualise in Figure 6:

**Figure 6.**
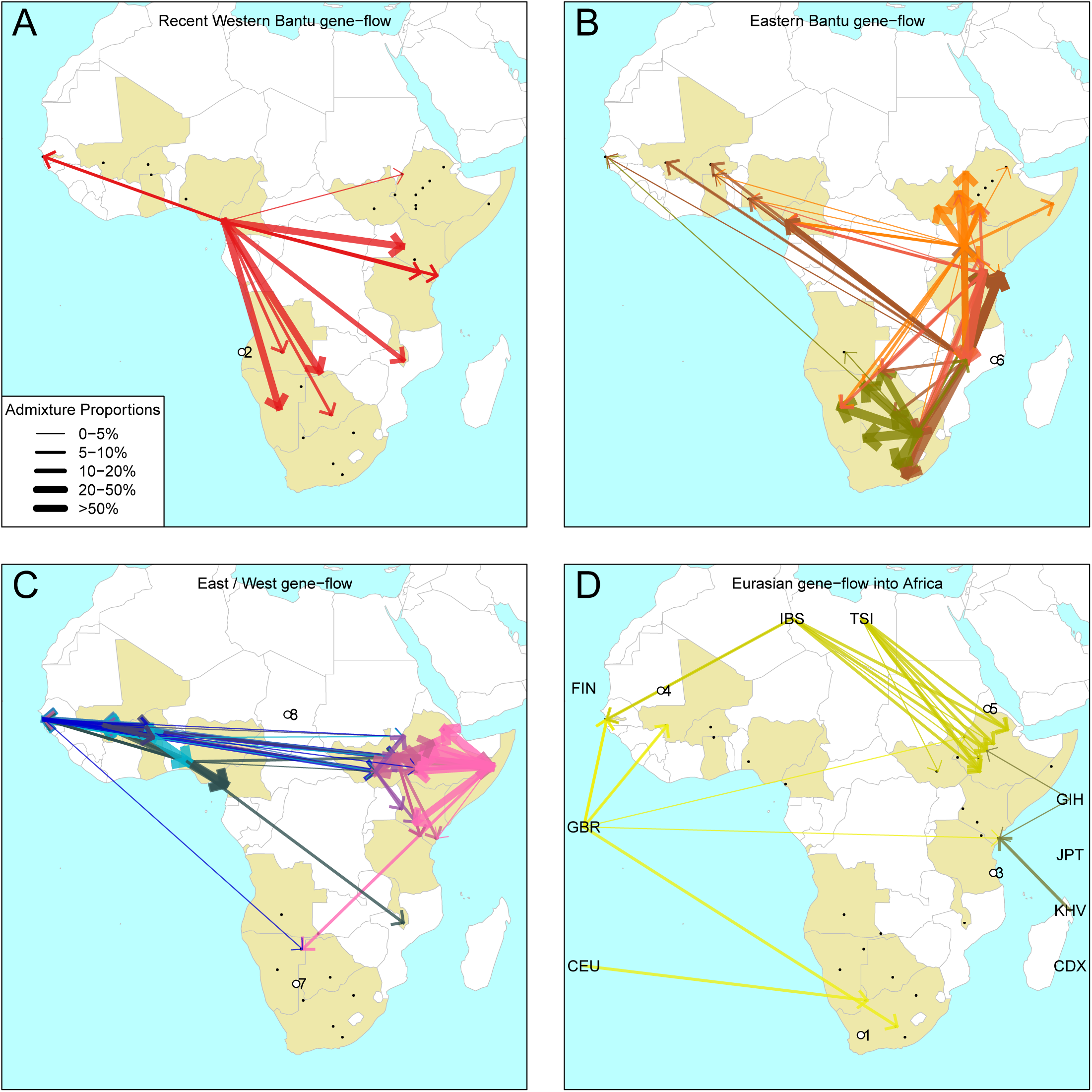
The geography of recent gene-flow in Africa. We summarise gene-flow events in Africa using the results of the GLOBETROTTER analysis. For each ethnic group, we inferred the composition of the admixture sources, and link recipient population to surrogates using arrows, the size of which is proportional to the amount it contributes to the admixture event. We separately plot (A) all events involving admixture source components from the Bantu and Semi-Bantu ethnic groups in Cameroon; (B) all events involving admixture sources from East and South African Niger-Congo speaking groups; (C) events involving admixture sources from West African Niger-Congo and East African Nilo-Saharan / Afroasiatic groups; (D) all events involving components from Eurasia. in (D) arrows are linked to the labelled 1KGP Eurasian groups. Arrows are coloured by country of origin, as in Figure 5. Numbers 1-8 in circles represent the events highlighted in section *A haplotype-based model of gene flow in sub-Saharan Africa*. An alternative version of this plot, stratified by date, is shown in Figure 6-figure supplement 1.

1. **Colonial Era European admixture in the Khoesan**. In two southern African Khoesan groups we see very recent admixture involving northern European ancestry which likely resulted from Colonial Era movements from the UK, Germany, and the Netherlands into South Africa [Thompson, 2001].
2. **The recent arrival of the Western Bantu expansion in southern Africa**. Central West African, and in particular ancestry from Cameroon (red ancestry in Figure 6A), is seen in Southern African Niger-Congo and Khoesan speaking groups, the Herero, Khwe and !Xun, indicating that the gradual diffusion of Bantu ancestry reached the south of the continent only within the last 750 years. Central West African ancestry in Malawi appears to have appeared prior to this event.
3. **Medieval contact between Asia and the East African Swahili Coast**. Specific Asian geneflow is observed into two coastal Kenyan groups, the Kauma and Chonyi, which represents a distinct route of Eurasian, in this case Asian, ancestry into Africa, perhaps as a result of Medieval trade networks between Asia and the Swahili Coast around 1200CE.
4. **Gene-flow across the Sahara**. Over the last 3,000 years, admixture involving sources containing northern European ancestry is seen on the Western periphery of Africa, in The Gambia and Mali. This ancestry in West Africa is likely to be the result of more gradual diffusion of DNA across the Sahara from northern Africa and across the Iberian peninsular, and not via the Middle East, as in the latter scenario we would expect to see Spanish (IBS) and Italian (TSI) in the admixture sources. We do see limited southern European ancestry in West Africa (Figs. 5 and 6D) in the Fulani, suggesting that some Eurasian ancestry may also have entered West Africa via North East Africa [Henn et al., 2012].
5. **Several waves of Mediterranean / Middle Eastern ancestry into north-east Africa**. We observe southern European gene flow into East African Afroasiatic speakers over a more prolonged time period over the last 3,000 years, with a major wave 2,000 years ago (Figs. 5 and 6D). We do not have Middle-Eastern groups in our analysis, so the observed Italian ancestry in the minor sources of admixture – the Tuscans are the closest Eurasian group to the Middle East – is consistent with previous results using the same samples [Pagani et al., 2012; Hodgson et al., 2014a], indicating this region as a major route for the back migration of Eurasian DNA into sub-Saharan Africa [Pagani et al., 2012; Pickrell et al., 2014].
6. **The late split of the Eastern Bantus**. The major source of admixture in East Africa Niger-Congo speakers is consistently a mixture of Central West Africa and Southern Niger-Congo speaking groups, in particular Malawi. This result best fits a model where Bantu speakers initially spread south along the western side of the Congo rainforest before splitting off eastwards, and interacting with local groups in central south Africa – for which Malawi is our best proxy – and then moving further north-east and south (Figure 6B).
7. **Pre-Bantu pastoralist movements from East to South Africa**. In the Ju/’hoansi we infer an admixture event involving an East African Afroasiatic source. This event precedes the arrival of Bantu-speaking groups in southern Africa, and is consistent with several recent results linking Eastern and Southern Africa and the limited spread of cattle pastoralism prior to the Bantu expansion (Figs. 5 and 6C) [Pickrell et al., 2014; Ranciaro et al., 2014; Macholdt et al., 2015; Barham and Mitchell, 2008].
8. **Ancestral connections between West Africans and the Sudan**. Concentrating on older events, we observe old “Sudanese” (Nilotic) components in very small proportions in the Gambia (Figure 4-figure supplement 1 and Figure 5) which may represent ancient expansion relationships between East and West Africa. When we infer admixture in West and Central West African groups without allowing any West Africans to contribute to the inference, we observe a clear signal of Nilo-Saharan ancestry in these groups, consistent with bidirectional movements across the Sahel [Tishkoff et al., 2009] and coancestry with (unsampled) Nilo-Saharan groups in Central West Africa. Indeed, if we look again at the PCA in Figure 1C, we observe that the Nilo-Saharan speakers are between West and East African Niger-Congo speaking individuals on PC3, an affinity which is supported by the presence of West African components in non-Niger-Congo speaking East Africans (Fig 6C).
9. **Ancient Eurasian gene-flow back into Africa and shared hunter-gatherer ancestry**. The MALDER analysis and *f*_3_ statistics show the general presence of ancient Eurasian and/or Khoesan ancestry across much of sub-Saharan Africa. We tentatively interpret these results as being consistent with recent research suggesting very old (>10 kya) migrations back into Africa from Eurasia [Hodgson et al., 2014a], with the ubiquitous hunter-gatherer ancestry across the continent possibly related to the inhabitant populations present across Africa prior to these more recent movements. Future research involving ancient DNA from multiple African populations will help to further characterise these observations.

## Discussion

Here we present an in-depth analysis of the genetic history of sub-Saharan Africa in order to characterise its impact on present day diversity. We show that gene-flow has taken place over a variety of different time scales which suggests that, rather than being static, populations have been sharing DNA, particularly over the last 3,000 years. An unanswered question in African history is how contemporary populations relate to those present in Africa before the transition to pastoralism that began some 2,000 years ago. Whilst the *f*_3_ and MALDER analyses show evidence for deep Eurasian and some hunter-gatherer ancestry across Africa, our GLOBETROTTER analysis provides a greater precision on the admixture sources and a timeline of events and their impact on groups in our analysis (Figure 6). The transition from foraging to pastoralism and agriculture in Africa was likely to be complex, with its impact on existing populations varying substantially. However, the similarity of language and domesticates in different parts of Africa implies that this transition spread; there are few cereals or domesticated animals that are unique to particular parts of Africa. Whilst herding has likely been going on for several thousand years in some form in Africa, 2,000 years ago much of Africa still remained the domain of hunter-gatherers [Barham and Mitchell, 2008]. Our analysis provides an attempt at timing the spread of Niger-Congo speakers east from Central West Africa. Although we do not have representative forager (hunter-gatherer) groups from all parts of sub-Saharan Africa, we observe that in addition to their local region, many of the sampled groups share ancestry with Central West African groups, which is likely the result of genes spreading with the Bantu agriculturalists as their farming technology spread across Africa.

After 0CE we begin to see admixture events shared between East and West Africa, with predominantly West to East direction, which appear to be most extensive around 1,000 years ago. During this time we see evidence of admixture from West, and West Central Africa into southern Khoesan speaking groups as well as down the Eastern side of the continent. Below, we outline some of the important technical challenges in using genetic data to interpret historical events exposed by our analyes. Nonetheless, the study presented here shows that patterns of haplotype sharing in sub-Saharan African are largely determined by historical gene-flow events involving groups with ancestry from across and outside of the continent.

### Interpreting haplotype similarity as historical admixture

Analyses that rely on correlations in allele frequencies (such as those performed here in the *Widespread evidence for admixture* section) provided initial evidence that the presence of Eurasian DNA across sub-Saharan Africa is the result of gene flow back into the continent within the last 10,000 years [Gurdasani et al., 2014; Pickrell et al., 2014; Hodgson et al., 2014a]. In addition, some groups have ancient (over 5 kya) shared ancestry with hunter-gather groups (Figure 3) [Gurdasani et al., 2014]. Whilst the weighted admixture LD decay curves between pairs of populations suggests that this admixture involved particular groups, for two main reasons, the interpretation of such events is difficult. Firstly, because our dataset includes closely related groups, in many populations, multiple pairs of reference populations generate significant admixture curves. Although the MALDER analysis helps us to identify which of these groups is closest to the true admixing source, it is not always possible to identify a single best matching reference, implying that sub-Saharan African groups share some ancestry with many different extant groups. So, as confirmed with the GLOBETROTTER analysis, it is unlikely that any single contemporary population adequately represents the true admixing source. On this basis of these analyses alone, it is not possible to characterise the composition of admixture sources.

Secondly, when we infer ancient events with ALDER, such as in the Mossi from Burkina Faso, where we estimate admixture around 5,000 years ago between a Eurasian (GBR) and a Khoesan speaking group (/Gui //Gana), we know that modern haplotypes are likely to only be an approximation of ancestral diversity [Pickrell and Reich, 2014]. Even the Ju/’hoansi, a San group from southern Africa traditionally thought to have undergone limited recent admixture, has experienced gene flow from non-Khoesan groups within this timeframe (Figure 4) [Pickrell et al., 2012, 2014]. Our result hints that the Mossi share deep ancestry with Eurasian and Khoesan groups, but any description of the historical event leading to this observation is potentially biased by the discontinuity between extant populations and those present in Africa in the past. In fact, this is one motivation for grouping populations into ancestry regions and defining admixture sources in this way. In this example we can (as we do) refer to Eurasian ancestry in general moving back into Africa, rather than British DNA in particular. Using contemporary populations as proxies for ancient groups is not the perfect approach, but it is the best that we have in parts of the world from which we do not have (for the moment) DNA from significant numbers of ancient human individuals, at sufficient quality, with which to calibrate temporal changes in population genetics.

An alternative approach is to characterise admixture as occurring between sources that have mixed ancestry themselves, to account for the fact that whilst groups in the past were not intact in the same way as they are now, DNA from such groups is nevertheless likely to be present in extant groups today. Haplotype-based methods allow for this type of analysis [Hellenthal et al., 2014; Leslie et al., 2015], and have the additional benefit of potentially identifying hidden structure and relationships (Figure 2). Whilst this approach still requires one to label groups by their present-day geographic or ethno-linguistic identity, it at least allows for the construction of admixture sources containing ancestry from multiple different contemporary groups, which is likely to be closer to the truth. It also provides a framework for describing historical admixture in terms of gene flow networks, which taps into the increasingly supported notion that the genetic variation present in the vast majority of modern human groups is the result of mixture events in the past [Pickrell and Reich, 2014; Hellenthal et al., 2014]. One drawback of such an approach is that it is not always possible to assign a specific historical population label to mixed admixture sources, but it nevertheless accommodates the need to express admixture sources as containing multiple different ancestries.

### Admixture in the context of infectious disease studies

Whilst inference of historical events are interesting in there own right, they also provide an important background for conducting large-scale genetic epidemiology in sub-Saharan Africa. Two studies designs are of particular relevance: genetics association studies and inference of historical natural selection. Principally, the haplotype analysis described here suggests that while recent gene-flow events mean that absolute allele frequency differences across the sub-Sahara are small relative to between-Eurasian groups, there remains substantial haplotype diversity. Genetic studies often rely on assumptions about the similarity of haplotypes between two groups, either as part of testing for association, or in order to impute genetic variation that is not assayed directly. Extensive recent gene-flow can help in this regard by reducing the differences between groups. However, the clear signals of admixture suggest that novel and divergent haplotypes are likely to be pocketed with subtle variation occurring both ethno-geographically, due to differences in ancestry which pre-dates recent movements, and along the genome, either by chance or by the differential effects of natural selection.

When new haplotypes are introduced into a population their fate will be partly be determined by the selective advantage they confer, as well as the action of genetic drift. These selective forces could include response to changes in infectious diseases, climate, and the cultural environmental [Coop et al., 2009; Fumagalli et al., 2015]. Additionally, important causes of morbidity and ill-health in Africa are due to highly polymorphic parasites like *Plasmodium falciparum*, where moving into new environments might lead to exposure to new strains. An implication of widespread gene-flow is that it can provide a route for potentially beneficial novel mutations to enter populations allowing them to adapt to such change. Recent examples of this phenomena include the presence of altitude adaptations in Tibetans from admixture with Sherpa [Jeong et al., 2014] and Denisovans [Huerta-Sánchez et al., 2014]; higher than expected frequencies of the Duffy-null mutation in populations from Madagascar as a result of admixture with African Bantu speaking groups [Hodgson et al., 2014b]; and the observation of shared haplotype(s) in humans and Neanderthals immunity genes belonging to the Toll-like Receptor (*TLR*) gene cluster as a result of archaic admixture [Deschamps et al., 2016; Dannemann et al., 2016].

In the context of malaria, the spread of the Duffy-null allele, an ancient mutation which arose at least 30,000 years ago [Hamblin and Di Rienzo, 2000; Hamblin et al., 2002] and which confers resistance to *P. vivax* malaria, throughout Africa is only possible through contact and gene flow between populations right across the sub-Sahara. Conversely, the same sickle-cell causing mutation appears to have recently occurred five times independently in Africa, causing multiple distinct haplotypes to be observed [Hedrick, 2011]. These mutations are young, within the order of 250-1750 years old [Currat et al., 2002; Modiano et al., 2008], so will have had limited opportunity to have been moved around by the gene-flow events that we describe.

We observed that some of this gene-flow may have coincided with the spread of Bantu languages and the concomitant introduction of new agricultural practices, which can jointly introduce both new genes and novel selective pressures into existing populations, such as specific agricultural products or unknown communicable disease. As an example, our analysis also sheds light on the spread of pastoralism into southern Africa prior to the Bantu expansion [Ehret, 1967; Guldemann, 2008; Henn et al., 2008]. Several recent resequencing studies have described the the distribution of mutations in and around the lactase gene (LCT) in African populations which are associated with the ability to digest lactase in adult life, lactase persistence (LP) [Ranciaro et al., 2014; Macholdt et al., 2015; Breton et al., 2014]. The C-14010 LP-associated variant, thought to have originated in East African Niger-Congo speaking groups [Tishkoff et al., 2007], was observed in the the San and ¡Xhosa, with the haplotype in the latter matching that of east African Bantu speaking populations [Ranciaro et al., 2014]. The same LCT variant was also observed at appreciable frequencies in the Nama [Breton et al., 2014]. We infer an admixture event in the Ju/’hoansi involving an Afroasiatic source, which at 732CE (616-993CE), is earlier than most of the Bantu admixture we observe in other southern African populations. In the Nama we infer a very recent admixture event (c. 1800CE) involving a source containing Niger-Congo (Malawi) ancestry. Whilst the event in the Nama is unlikely to have delivered the East African mutation to this group, the event in the Ju/’hoansi is consistent with pre-Bantu interactions between east African and southern African groups. Conversely, in the AmaXhosa the Bantu source of admixture derives most of its ancestry from East Africa, so the introduction the C-14010 mutation into this part of Africa may well have been the result of the Bantu expansion.

These examples demonstrate the utility of detailed inference exploiting the complex information that is captured by large-scale genome-wide studies of genetic diversity. Africa has an exciting opportunity to be part of bringing the genetic revolution to bear in understanding and treating both infectious and non-communicable disease. Further exploration and interpretation the the rich genetic diversity of the continent will help in this endeavour.

## Materials and Methods

### 1 Overview of the dataset

The dataset comprises a mixture of 2,504 previously published individuals from Africa and elsewhere (see below) plus novel genotypes on 1,712 sampled by the Malaria Genomic Epidemiology Network (MalariaGEN) Figure 1-Source Data 1. The MalariaGEN samples were a subset of those collected at 8 locations in Africa as part of a consortial project on genetic resistance to severe malaria: details of the study sites and investigators involved are described elsewhere [Malaria Genomic Epidemiology Network, 2014]. Samples were genotyped on the Illumina Omni 2.5M chip in order to perform a multicentre genome-wide association study (GWAS) of severe malaria: initial GWAS findings from The Gambia, Kenya and Malawi have already been reported [Band et al., 2013; Malaria Genomic Epidemiology Network, 2015] and a manuscript describing findings at all 8 locations is in preparation

The MalariaGEN samples used in the present analysis were selected to be representative of the main ethnic groups present at each of the 8 African study sites. We screened the samples collected at each study site (typically >1000 individuals) to select individuals whose reported parental ethnicity matched their own ethnicity. This process identified 23 ethnic groups for which we had samples for approximately 50 unrelated individuals or more. For ethnic groups with more than 50 samples available, we performed a cluster analysis on cohort-wide principle components, generated as part of the GWAS, with the R statistical programming language [R Development Core Team, 2011] using the *MClust* package [Fraley et al., 2012], choosing individuals from the cluster containing the largest number of individuals, to avoid any accidental inclusion of outlying individuals and to ensure that the 50 individuals chosen were, when possible, relatively genetically homogeneous. We note that in several ethnic groups (Malawi, the Kambe from Kenya, and the Mandinka and Fula from The Gambia; Fig. 1-figure supplement 2) PCA of the genotype data showed a large amount of population structure. In these cases we chose two sub-groups of individuals from a given ethnic group, selected to represent the diversity of ancestry depicted by the PCs. In several other cases, following GWAS quality control (see below), genotype data for fewer than 50 control individuals was available, and in these cases we chose as many individuals as possible, regardless of the PC-based clustering or case/control status.

We additionally included further individuals from each of four Gambian ethnic groups: the Fula, Mandinka, Jola, Wollof. The genotype data from these individuals were included as the same individuals are also being sequenced as part of the **Gambian Genome Variation Project**^1^. These subsets included ˜30 trios from each ethnic group, information on which was used in phasing (see below). Full genome data from these individuals will be made available in the future.

#### 1.1 Quality Control

Detailed quality control (QC) for the MalariaGEN dataset was performed for a genome-wide association study of severe malaria in Africa and is outlined in detail elsewhere [Malaria Genomic Epidemiology Network, 2015]. Briefly, genotype calls were formed by taking a consensus across three different calling algorithms (Illuminus, Gencall in Illumina’s BeadStudio software, and GenoSNP) [Band et al., 2013] and were aligned to the forward strand. Using the data from each country separately, SNPs with a minor allele frequency of <1% and missingness <5% were excluded, and additional QC to account for batch effects and SNPs not in Hardy-Weinberg equilibrium was also performed.

#### 1.2 Combining the MalariaGEN populations with additional populations

The post-QC MalariaGEN data was combined with published data typed on the same Illumina 2.5M Omni chip from 21 populations typed for the 1000 Genomes Project (1KGP)^2^, including and accounting for duos and trios, and with publicly available data from individuals from several populations from Southern Africa (Figure 1-Source Data 1) [Consortium, 2012; Schlebusch et al., 2012]. We merged samples to the forward strand, removing any ambiguous SNPs (A to T or C to G). Merging was checked by plotting allele frequencies between populations from both datasets, which should be generally correlated (data not shown). To describe population structure across sub-Saharan Africa we combined the above dataset with further publicly available samples typed on different Illumina (Omni 1M) chips, containing individuals from southern Africa [Petersen et al., 2013] and the Horn of Africa (Somalia/Ethiopia/Sudan)[Pagani et al., 2012] to generate a final dataset containing 4,216 individuals typed on 328,176 high quality common SNPs. To obtain the final set of analysis individuals we performed additional sample QC after phasing and removed American 1KG populations (Figure 1-Source Data 1).

#### 1.3 Phasing

We used SHAPEITv2 [Delaneau et al., 2012] to generate haplotypically phased chromosomes for each individual. SHAPEITv2 conditions the underlying hidden Markov model (HMM) from Li and Stephens [2003] on all available haplotypes to quickly estimate haplotypic phase from genotype data. We split our dataset by chromosome and phased all individuals simultaneously, and used the most likely pairs of haplotypes (using the –*output-max* option) for each individual for downstream applications. We performed 30 iterations of the MCMC and used default values for all other parameters. As mentioned, we used known pedigree relationships to improve the phasing, using family data from both the 1KG and the Gambia Sequencing Projects.

#### 1.4 Removing non-founders and cryptically related individuals

Our dataset included individuals who were known to be closely related (1KG duos and trios; Gambia Sequencing Project trios) and, because we took multiple samples from some population groups, there was also the potential to include cryptically related individuals. After phasing we therefore performed an additional step where we first removed all non-founders from the analysis and then identified individuals with high identity by descent (IBD), which is a measure of relatedness. Using an LD pruned set of SNPs generated by recursively removing SNPs with an *R*^2^ > 0.2 using a 50kb sliding window, we calculated the proportion of loci that are IBD for each pair of individuals in the dataset using the R package *SNPRelate* [Zheng et al., 2012] and estimated kinship using the pi-hat statistic (the proportion of loci that are identical for both alleles (IBD=2) plus 0.5* the proportion of loci where one allele matches (IBD=1); i.e. PI HAT=P(IBD=2)+0.5*P(IBD=1)). For any pair of individuals where IBD > 0.2, we randomly removed one of the individuals. 327 individuals were removed during this step.

#### 1.5 1KG American populations and Native American Ancestry in 1KG Peruvians

With the exception of Peru, post-phasing we dropped all 1KG American populations from the analysis (97 ASW, 102 ACB, 107 CLM, 100 MXL and 111 PUR). We used a subset of the 107 Peruvian individuals that showed a large amount of putative Native American ancestry, with little apparent admixture from non-Amerindians (data not shown). Although Amerindians are not central to this study, and it is unlikely that there has been any recurrent admixture from the New World into Africa, we nevertheless generated a subset of 16 Peruvians to represent Amerindian admixture components in downstream analyses. When this subset was used, we refer to the population as PELII. The removal of these 606 American individuals left a final analysis dataset comprising 3,283 individuals from 60 different population groups (Figure 1-Source Data 1).

### 2 Analysis of population structure in sub-Saharan Africa

#### 2.1 Principal Components Analysis

We performed Principal Components Analysis (PCA) using the SNPRelate package in R. We removed SNPs in LD by recursively removing SNPs with an *R*^2^ > 0.2, using a 50kb sliding window, resulting in a subset of 162,322 SNPs.

#### 2.2 Painting chromosomes with CHROMOPAINTER

We used fineSTRUCTURE [Lawson et al., 2012] to identify finescale population structure and to identify high level relationships between ethnic groups. The initial step of a fineSTRUCTURE analysis involves “painting” haplotypically phased chromosomes sequentially using an updated implementation of a model initially introduced by Li and Stephens [2003] and which is exploited by the CHROMOPAINTER package [Lawson et al., 2012]. The Li and Stephens copying model explicitly relates linkage disequilibrium to the underlying recombination process and CHROMOPAINTER uses an approximate method to reconstruct each “recipient” individual’s haplotypic genome as a series of recombination “chunks” from a set of sample “donor” individuals. The aim of this approach is to identify, at each SNP as we move along the genome, the closest relative genome among the members of the donor sample. Because of recombination, the identity of the closest relative will change depending on the admixture history between individual genomes. Even distantly related populations share some genetic ancestry since most human genetic variation is shared [The International HapMap Consortium, 2010; Ralph and Coop, 2013], but the amount of shared ancestry can differ widely. We use the term “painting” here to refer to the application of a different label to each of the donors, such that – conceptually – each donor is represented by a different colour. Donors may be coloured individually, or in groups based on a priori defined labels, such as the geographic population that they come from. By recovering the changing identity of the closest ancestor along chromosomes we can understand the varying contributions of different donor groups to a given population, and by understanding the distribution of these chunks we can begin to uncover the historical relationships between groups.

#### 2.3 Using painted chromosomes with fineSTRUCTURE

We used CHROMOPAINTER with 10 Expectation-Maximisation (E-M) steps to jointly estimate the program’s parameters Ne and *θ*, repeating this separately for chromosomes 1, 4, 10, and 15 and weight-averaging (using centimorgan sizes) the Ne and *θ* from the final E-M step across the four chromosomes. We performed E-M on 5 individuals from every population in the analysis and used a weighted average of the values across all pops to arrive at final values of 190.82 for Ne and 0.00045 for *θ*. We ran each chromosome from each population separately and combined the output to generate a final coancestry matrix to be used for fineSTRUCTURE.

As the focus of our analysis is population structure within Africa, we used a “continental force file” to combine all non-African individuals into single populations. The processing time of the algorithm is directly related to the number of individuals included in the analysis, so reducing the number of individuals speeds the analysis up. Furthermore, fineSTRUCTURE initially uses a prior that assumes that all individuals are equally distant from each other, which in the case of worldwide populations is likely to be untrue: African populations are likely to be more closely related to each other than to non-Africa populations, for example. The result is that not all of the substructure is identified in one run.

We therefore combined all individuals from each of the non-Africa 1KG populations into “continents”, which has the effect of combining all of the copying vectors from the individuals within them to look like (re-weighted) normal individuals but cannot be split and do not contribute to parameter inference, and can thus be considered as copying vectors that contain the average of the individuals within them. They are then included in the algorithm at minimal extra computational cost and exist primarily to provide chunks to (and from) the remaining groups. We combined all individuals from a labelled population (e.g. all IBS individuals were now contained in a “continent” grouping called IBS), with the exception of the three Chinese population CHB, CHD, and CHS where we combined all individuals into a single CHN continent.

#### 2.4 Using fineSTRUCTURE to inform population groupings

The fineSTRUCTURE analysis identified 154 clusters of individuals, grouped on the basis of copying-vector similarity (Fig. 1-figure supplement 3). Some ethnic groups, such as Yoruba, Mossi, Jola and Ju/’hoansi form clusters containing only individuals from their own ethnic groups. In other cases, individuals from several different ethnic groups are shared across different clusters. We see this particularly in groups from The Gambia and Kenya. These are the two countries where the most ethnic groups were sampled, seven and four, respectively, and this putative mixing could be a result of sampling several different ethnic groups from a limited geographic area. The reason that we do not see the same extent of haplotype sharing in Cameroon, for example, may be because we only have two ethnic groups sampled from that area. The mixed nature of the clusters in this region could also be the results of real ancestral relationships: some of the ethnic groups in The Gambia, for example, may have history of choosing partners from outside their own ethnic group.

As mentioned above, we combined data from different sources, and in some groups (e.g. the Fula) we specifically chose different groups of individuals in an attempt to cover the broad spectrum of ancestry present in that group. We used the fineSTRUCTURE tree to visually group individuals based on their ancestry. Our aim here is to try to maximise the number of individuals that we can include within an ethnic group, without merging together individuals that are distant on the tree. We also decided *not to* use the fineSTRUCTURE clusters themselves as analytical groups because of difficulties with the interpretation of the history of such clusters. We were interested in identifying the major admixture events that have occurred in the history of different populations, and it is not clear what an analytical group that is defined as, for example, a mixture of Manjago, Mandinka and Serere individuals, would mean in our admixture analyses. In practice, this meant that we used the original geographic population labelled groups for all populations except in the Fula and Mandinka from The Gambia, where individuals fell into two distinct groups of clusters. Here we defined two clusters for each group, with the the two groups suffixed with an “I” or “II’ (Figure 1-figure supplement 3).

#### 2.5 Defining ancestry regions

We used a combination of genetic and ethno-linguistic information (see Supplementary File 1 below) to define seven ancestry regions in sub-Saharan Africa. The ancestry regions are reported in Figure 1-Source Data 1 and closely match the high level groupings we observed in the fineSTRUCTURE tree, with the following exceptions:

1. East African Niger-Congo speakers

a. The two ethnic groups from Cameroon – Bantu and Semi-Bantu – were included in the Central West African Niger-Congo ancestry region despite clustering more closely with East African groups from Kenya and Tanzania in Figure 1-figure supplement 3. In a preliminary fineSTRUCTURE analysis based on the MalariaGEN and 1KG populations only, using c. 1 million SNPs, the Cameroon populations clustered with other Central West African groups, and not East Africans (data not shown).
b. Malawi was included in the South African Niger-Congo ancestry region, despite being an outlying cluster in a clade with East African Niger-Congo speaking groups. A preliminary fineSTRUCTURE analysis based on the MalariaGEN, 1KG and Schlebusch populations, clustered Malawi with the Herero and SEBantu speakers (data not shown).
2. Southern Africa

a. We treated Southern African individuals slightly differently: even though the fineSTRUCTURE analysis did not split them into two separate clades of Khoesan and Niger-Congo speaking individuals, we nevertheless did. Schlebusch et al. [2012] showed that these populations were inter-related and admixed, two properties in the data we were hoping to uncover. The final ancestry region assignments are outlined in Figure 1-Source Data 1.

#### 2.6 Estimating pairwise *F_ST_*

We used *smartpca* in the EIGENSOFT [Patterson et al., 2006] package to estimate pairwise *F_ST_* between all populations. This implementation uses the Hudson estimator recently recommended by Bhatia et al. [2013]. Results are shown in Figure 2, Figure 2-Source Data 1 and Figure 2-Source Data 2.

#### 2.7 Comparing sets of copying vectors

We used Total Variation Distance (TVD) to compare copying vectors [Leslie et al., 2015; van Dorp et al., 2015]. As the copying vectors are discrete probability distributions over the same set of donors, TVD is a natural metric for quantifying the difference between them. For a given pair of groups *A* and *B* with copying vectors describing the copying from *n* donors, *a* and *b*, we can compute TVD with the following equation:

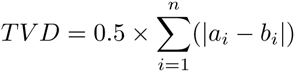

### 3 Analysis of admixture in sub-Saharan African populations

We used a combination of approaches to explore admixture across Africa. Initially, we employed commonly used methods that utilise correlations in allele frequencies to infer historical relationships between populations. To understand ancient relationships between African groups we used the *f*_3_ statistic [Reich et al., 2009] to look for shared drift components between a test population and two reference groups. We next used ALDER [Loh et al., 2013] and MALDER [Pickrell et al., 2014] – an updated implementation of ALDER that attempts to identify multiple admixture events – to identify admixture events through explicit modelling of admixture LD by generating weighted LD curves. The weightings of these curves are based on allele frequency differences, at varying genetic distances, between a test population and two putative admixing groups.

To identify more recent events we used two methods which aim to more fully model the mixed ancestry in a population by utilising the distribution and length of shared tracts of ancestry as identified with the CHROMOPAINTER algorithm [Lawson et al., 2012; Hellenthal et al., 2014]. We outline the details of this analysis below, but note here that, because this approach is based on the comparison and analysis of painted chromosomes, it offers a different perspective from approaches based on comparisons of allele frequencies.

#### 3.1 Inferring admixture with the *f*_3_ statistic and ALDER

We computed the *f*_3_ statistic, introduced by Reich et al. [2009], as implemented in the TREEMIX package [Pickrell and Pritchard, 2012]. These tests are a 3-population generalization of *F_ST_*, equal to the inner product of the frequency differences between a group X and two other groups, A and B. The statistic, commonly denoted *f*_3_(X:A,B) is proportional to the correlated genetic drift between A and X and A and B. If X is related in a simple way to the common ancestor with A and B, we expect this quantity to be positive. Significantly negative values of *f*_3_ suggest that X has arisen as a mixture of A and B, which is thus an unambiguous signal of mixture. Standard errors are computed using a block jackknife procedure in blocks of 500 SNPs (Supplementary Table 1).

This analysis shows that sub-Saharan African populations tend to have either a hunter-gatherer or Eurasian source contributing to their most significant *f*_3_ statistics (Supplementary Table 1). That is, there is clear evidence of admixture involving both Eurasian and hunter-gatherer groups across sub-Saharan Africa. Whilst we do not infer admixture using this statistic in the Jola, Kasem, Sudanese, Gumuz, and Ju/’hoansi, in most other groups the most significant *f*_3_ statistic includes either the Ju/’hoansi or a 1KGP European source (GBR, CEU, FIN, or TSI). Niger-Congo speaking groups from Central West and Southern Africa tend to show most significant statistics involving the Ju/’hoansi, where as West and East African and Southern Khoesan speaking groups tended to show most significant statistics involving European sources, consistent with an recent analysis on a similar (albeit smaller) set of African populations [Gurdasani et al., 2014].

We used ALDER [Patterson et al., 2012; Loh et al., 2013] to test for the presence of admixture LD in different populations. This approach works by generating weighted admixture curves for pairs of populations and tests for admixture. As noted in Loh et al. [2013] the use of *f*_3_ statistics and weighted LD curves are somewhat complementary, and there are several reasons why *f*_3_ statistics might pick up signals of admixture when ALDER does not. In particular, admixture identified using *f*_3_ statistics but not by ALDER is potentially related to more ancient events because whilst shared drift signals will still be present, admixture LD will have been broken over (potentially) millennia of recombination.

As previously shown by Loh et al. [2013] and Pickrell et al. [2014], weighted LD curves can be used to identify the source of the gene flow by comparing curves computed using different reference populations. This is possible because theory predicts that the amplitude (i.e. the y-axis intercept) of these curves becomes larger as one uses reference populations that are closer to the true mixing populations. Loh et al. [2013] demonstrated that this theory holds even when using the admixed population itself as one of the reference populations. Pickrell et al. [2014] used this concept to identify west Eurasian ancestry in a number of East African and Khoesan speaking groups from southern Africa.

We thus initially ran ALDER in “one-reference” mode, where for each focal population, we generated curves involving itself with every other reference population in turn. We used the average amplitude of the curves generated in this way to identify the groups important in describing admixture in the history of the focal group. Figure 3-figure supplement 1 shows comparative plots to those by Pickrell et al. [2014] for a selection of African populations, including the Ju/’hoansi, who we also infer to have largest curve amplitudes with Eurasian groups, consistent with that previous analysis. We summarise the results of the analysis of curve amplitudes across all populations in Figure 3-figure supplement 2, which shows the rank of the curve amplitudes for each reference population across different sub-Saharan African groups. Across much of Africa, amplitudes from non-African (European) groups are amongst the highest, indicative of admixture from Eurasian-like sources.

Next, for each focal ethnic group in turn, we used ALDER to characterise admixture using all other ethnic groups as potential reference groups (i.e. in two-population mode). In effect, this approach compares every pair of reference groups, identifying those pairs that show evidence of shared admixture LD (P <0.05 after multiple-hypothesis testing). As many of the groups are closely related, we often observed more than one pair of ethnic groups as displaying admixture in a given focal population, the results of which are highly correlated. In Supplementary Table 1, we show the evidence for admixture only for the pair of groups with the lowest P-value for each focal group. Dates for admixture events were generated using a generation time of 29 years [Fenner, 2005] and the following equation:

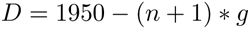

where *D* is the inferred date of admixture, n is the inferred number of generations since admixture, and *g* is the generation time in years.

#### 3.2 Inferring multiple waves of admixture in African populations using weighted LD curves

We used MALDER [Pickrell et al., 2014], an implementation of ALDER designed to fit multiple exponentials to LD decay curves and therefore characterise multiple admixture events to allele frequency data. For each event we recorded (a) the curve, *C*, with the largest overall amplitude 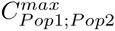, and (b) the curves which gave the largest amplitude where each of the two reference populations came from a different ancestry region, and for which a significant signal of admixture was inferred. To identify the source of an admixture event we compared curves involving populations from the same ancestry region as the two populations involved in generating 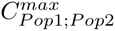. For example, in the Jola, the population pair that gave *C^max^* were the Ju/’hoansi and GBR. Substituting these populations for their ancestry regions we get 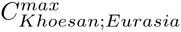. To understand whether this event represents a specific admixture involving the Khoesan in the history of the Jola, we identified the amplitude of the curves from (b) of the form 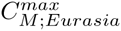, where *M* represents a population from any ancestry region other than Eurasia that gave a significant MALDER curve. We generated a Z-score for this curve comparison using the following formula [Pickrell et al., 2014]:

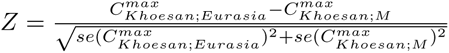

The purpose of this was to determine, for a given event, whether the sources of admixture could be represented by a single ancestry region, in which case the overall 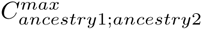 will be significantly greater than curves involving other regions, or whether populations from multiple ancestry regions can generate admixture curves with similar amplitudes, in which case there will be a number of ancestry regions that best represent the admixing source. We combined all values of *M* where the Z-score computed from the above test gave a value of < 2, and define the sources of admixture in this way.

To identify the major source of admixture, we performed a similar test. We determined the regional identity of the two populations used to generate *C^max^*. In the example above, these are Khoesan and Eurasia. Separately for each region, we identify the curve, *C*, with the maximum amplitude where either of the two reference populations was from the Khoesan region, 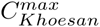 as well as the curve where neither of the reference populations was Khoesan, 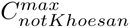. We compute a Z-score as follows:

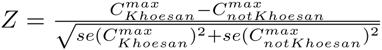

This test generates two Z-scores, in this example, one for the Khoesan/not-Khoesan comparison, and one for the Eurasia/not-Eurasia comparison. We assign the main ancestry of an event to be the region(s) that generate Z > 2. If neither region generates a Z-score > 2, then we do not assign a major ancestry to the event.

#### 3.3 Comparisons of MALDER dating using the HAPMAP worldwide and African-specific recombination maps

Recombination maps inferred from different populations are correlated on a broad scale, but differ in the fine-scale characterisation of recombination rates [Hinch et al., 2011]. Having observed significantly older dates in many West and Central West African groups, we investigated the effect of recombination map choice by recomputing MALDER results with an African specific genetic map which was inferred through patterns of LD from the HAPMAP Yoruba (YRI) sample (Fig. 3C).

We next re-inferred admixture parameters with MALDER using all populations with the African (YRI) and additionally with a European (CEU) map [Hinch et al., 2011]. We show comparison of the dates inferred with these different maps in the main paper, and here we shows the equivalent figures to Figure 3 for events inferred using the African (Fig. 3-figure supplement 6) and European (Fig. 3-figure supplement 7) maps.

#### 3.4 Analysis of the minimum genetic distance over which to start curve fitting when using ALDER/MALDER

A key consideration when using weighted LD to infer admixture parameters is the minimum genetic distance over which to begin computing admixture curves. Short-range LD correlations between two reference populations and a target may not only be the result of admixture, but may also be due to demography unrelated to admixture, such as shared recent bottlenecks between the target population and one of the references, or from an extended period of low population size [Loh et al., 2013]. Indeed, the authors of the ALDER algorithm specifically incorporate checks into the default ALDER analysis pipeline that define the threshold at which a test population shares short-range LD with with either of the two reference populations. Subsequent curve analyses then ignore data from pairs of SNPs at smaller distances than this correlation threshold [Loh et al., 2013].

The authors nevertheless provide the option of over-riding this LD correlation threshold, allowing the user to define the minimum genetic distance over which the algorithm will begin to compute curves and therefore infer admixture. So there are (at least) two different approaches that can be used to infer admixture using weighted LD. The first is to infer the minimum distance to start building admixture curves from the data (the default), and the second is to assume that any short-range correlations that we observe in the data result from true admixture, and prescribe a minimum distance over which to infer admixture.

#### 3.5 All African populations share correlated LD at short genetic distances

We tested these two approaches by inferring admixture using MALDER/ALDER using a minimum distance defined by the data on the one hand, and a prescribed minimum distance of 0.5cM on the other. This value is commonly used in MALDER analyses, for example by the African Genome Variation Project [Gurdasani et al., 2014]. For each of the 48 African populations as a target, we used ALDER to infer the genetic distance over which LD correlations are shared with every other population as a reference. In Figure 3-figure supplement 3 we show the distribution of these values across all targets for each reference population.

Across all African populations we observe LD correlations with other African populations at genetic distances > 0.5cM, with median values ranging between 0.7cM when GUMUZ is used as a reference to 1.4cM when FULAII is used as a reference. In fact, when we further explore the range of these values across each region separately (Fig. 3-figure supplement 3), we note that, as expected, these distances are greater between more closely related groups.

When we plot the inverse of these distributions, that is, the range of these values for each target population separately, we see that the median minimum distance that all sub-Saharan African populations is always greater than 0.5cM (Fig. 3-figure supplement 4). Taken together, these results suggest that all African populations share some LD over short genetic distances, that may be the result of shared demography or admixture. In order to reduce the confounding effect of demography, with the exception of the next section, all ALDER/MALDER analyses presented in the paper were performed after accounting for this short range shared LD.

#### 3.6 Results of MALDER analysis using a fixed minimum genetic distance of 0.5cM

In order to compare our MALDER analysis to previously published studies, we show the results of the MALDER analysis where we fixed the minimum genetic distance to 0.5cM (Figure 3-figure supplement 5). The main differences between this analysis and that presented in the main part of the paper are:

1. Ancient (>5ky) admixture in Central West African populations where the main analysis found no signal of admixture
2. A second ancient admixture in Malawi c.10ky
3. More of the events appear to involve Eurasian and Khoesan groups mixing.

#### 3.7 Chromosome painting for mixture model and GLOBETROTTER

For the mixture model and GLOBETROTTER analyses, we generated painted samples where we disallowed closely related groups from being painting donors. In practice, this meant removing all populations from the same ancestry region as a given population from the painting analysis. The exception to this are populations from the Nilo-Saharan and Afroasiatic ancestry regions. In these groups, no population from either ancestry region was used as painting donors. We refer to this as the “non-local” painting analysis.

#### 3.8 Modelling populations as mixtures of each other using linear regression

Copying vector summaries generated from painted chromosomes describe how populations relate to one another in terms of the relative time to a common shared ancestor, subsequent recent admixture, and population-specific drift [Hellenthal et al., 2014; Leslie et al., 2015]. For the following analysis, we used the GLOBETROTTER package to generate the mixing coefficients used in Figure 4.

Given a number of potential admixing donor populations, a key step in assessing the extent of admixture in a given population *k* is to identify which of these donors is relevant; that is, we want to identify the set *T\** containing all populations *l* ≠ *k* ∈ [1,…, *K*] believed to be involved in any admixture generative to population *k*. Using copying vectors from the non-local painting analysis, we generate an initial estimate of the mixing coefficients that describe the copying vector of population *k* by fitting *f^k^* as a mixture of *f^l^* where *l* ≠ *k* ∈ [1,…, *K*]. The purpose of this step is to assess the evidence for putative admixture in our populations, as described by Hellenthal et al. [2014] and Leslie et al. [2015]. In practice, we remove the self-copying (drift) element from these vectors, i.e. we set 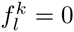, and rescale each population’s copying vector such that 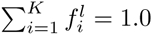 for all *l* = *k* ∈ [1,…, *K*].

We assume a standard linear model form for the relationship between *f^k^* and terms *f^l^* for *l* ≈ *k* ∈ [1,…, *K*]:

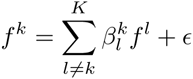

where ε is a vector of errors which we seek to choose the *β* terms to minimise using non negative least squares regression with the R “nnls” package. Here, 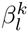 is the coefficient for *f^l^* under the mixture model, and we estimate the 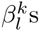 under the constraints that all 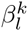 ≥ 0 and 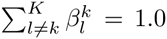. We refer to the estimated coefficient for the *l^th^* population as 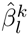; to avoid over-fitting we exclude all populations for which 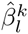 < 0.001 and rescale so that 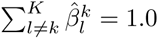. *T\** is the set of all populations whose 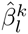 > 0.001.

The 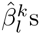 represent the mixing coefficients that describe a recipient population’s DNA as a linear combination of the set *T\** donor populations. This process identifies donor populations whose copying vectors match the copying vector of the recipient, as inferred by the painting algorithm.

#### 3.9 Overview of GLOBETROTTER analysis pipeline

In the current setting we are interesting in identifying the general historical relationships between the different African and non-African groups in our dataset. We used GLOBETROTTER [Hellenthal et al., 2014] to characterise patterns of ancestral gene flow and admixture. Individuals tend to share longer stretches of DNA with more closely related individuals, so we used a focused approach where we disallowed copying from local populations.

GLOBETROTTER was originally described by Hellenthal et al. [2014] and a detailed description of the algorithm and the extensive validation of the method is presented in that paper. Here we run over the general framework as used in the current study, with the key difference between our approach and the default use of the algorithm being that we do not allow any groups from within the same ancestry region as a target group to be donors in the painting analysis. Throughout we use GLOBETROTTERv2.

For a given test population *k*:

1. We define the set of populations present in the same broad ancestry region as *k* as *m*, with the caveats outlined below. Using CHROMOPAINTER, we generate painting samples using a reduced set of donors *T\**, that included only those populations not present in the same region of Africa as *k*, i.e. *T\** = *l* ≠ *m* ∈ [1,…, *K*]. The effect of this is to generate mosaic painted chromosomes whose ancestral chunks do not come from closely related individuals or groups, which can mask more subtle signals of admixture. For each group in turn, prior to this final painting, we ran 10 iterations of CHROMOPAINTER’s EM algorithm to infer the population-specific prior copying probabilities (using the -*ip* flag), and use these for the final sampled paintings.
2. For each population, *l* in *T\**, we generate a copying-vector, *f^l^*, allowing all individuals from *l* to copy from every individual in *T\**; i.e. we paint every population *l* with the same set of restricted donors as *k* in (1). For each recipient and surrogate group in turn, we sum the chunklengths donated by all individuals within all of our final donor groups (i.e. all 59 groups: including the recipient’s own group) and average across all recipients to generate a single 59 element copying vector for each recipient group.
3. To account for noise due to haplotype sharing among groups, we perform a non-negative-leastsquares regression (mixture model; outlined above) that takes the copying vector of the recipient group as the response and the copying vectors for each donor group as the predictors. We take the coefficients of this regression, which are restricted to be ≥ 0 and to sum to 1 across donors, as our initial estimates of mixing coefficients describing the genetic make-up of the recipient population as a mixture of other sampled groups.
4. Within and between every pairing of 10 painting samples generated for each haploid of a recipient individual, we consider every pair of chunks (i.e. contiguous segments of DNA copied from a single donor haploid) separated by genetic distance *g*. For every two donor populations, we tabulate the number of chunk pairs where the two chunks come from the two populations. This is done in a manner to account for phasing switch errors, a common source of error when inferring haplotypes.
5. An appropriate weighting and rescaling of the curves calculated in step 4 gives us the observed coancestry curves illustrating the decay in ancestry linkage disequilibrium versus genetic distance. There is one such curve for each pair of donor populations.
6. We find the maximum likelihood estimate (MLE) of rate parameter λ of an exponential distribution fit to all coancestry curves simultaneously. Specifically, we perform a set of linear regressions that takes each curve in turn as a response and the exponential distribution with parameter λ as a predictor, finding the λ that minimizes the mean-squared residuals of these regressions. This value of λ is our estimated date of admixture. We take the coefficients from each regression. (In the case of 2 dates, we fit two independent exponential distributions with separate rate parameters to all curves simultaneously and take the MLEs of these two rate parameters as our estimates of the two respective admixture dates. We hence get two sets of coefficients, with each set representing the coefficients for one of the two exponential distributions.)
7. We perform an eigen decomposition of a matrix of values formed using the coefficients inferred in step 6. (In the case of 2 dates, we perform an eigen decomposition of each of the two matrices of coefficients, one for each inferred date.)
8. We use the eigen decomposition from step 7 and the copying vectors to infer both the proportion of admixture *α* and the mixing coefficients that describe each of the admixing source groups as a linear combination of the donor populations. (In the case of 2 dates, we perform separate fits on each of the two eigen decompositions described in step 7 to describe each admixture event separately.)
9. We re-estimate the mixing coefficients of step 3 to be 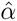 times the inferred mixing coefficients of the first source plus 1 – 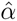 times the inferred mixing coefficients of the second source.
10. We repeat steps 5-9 for five iterations.
11. We repeat steps 4-5 using a new set of coancestry curves that should eliminate any putative signal of admixture (by taking into account the background distribution of chunks, the so-called **null procedure**), normalize our previous curves using these new ones, and repeat steps 6-10 to reestimate dates using these normalized curves. We generate 101 date estimates via bootstrapping and assess the proportion of inferred dates that are = 1 or ≥ 400, setting this proportion as our empirical *p*-value for showing *any* evidence of admixture.
12. Using values calculated in the final iteration of step 10, we classify the admixture event into one of five categories as: (A) ‘no admixture’, (B) ‘uncertain’, (C) ‘one date’, (D) ‘multiple dates’ and (E) ‘one date, multiway’.

#### 3.10 Inferring admixture with GLOBETROTTER

We use the painting samples from (1) and the copying-vectors from (2) detailed in Section 3.9 to implement GLOBETROTTER, characterising admixture in group *k;* the intuition being that any admixture observed is likely to be representative of gene-flow from across larger geographic scales.

We report the results of this analysis in Figure 4-Source Data 1 as well as Figures 4, 5, and 6. We generate date estimates by simultaneously fitting an exponential curve to the coancestry curves output by GLOBETROTTER and generate confidence intervals based on 100 bootstrap replicates of the GLOBETROTTER procedure, each time bootstrapping across chromosomes. Because it is unlikely that the true admixing group is present in our set of donor groups, GLOBETROTTER infers the sources of admixture as mixtures of donor groups, which are in some sense equivalent to the *β* coefficients described above, but are inferred using the additional information present in the coancestry curves. We infer the composition of the admixing sources by using the *β*s output by GLOBETROTTER from the two (or more) sources of admixture to arrive at an understanding of the genetic basis of the the admixing source groups. These contrasts show us the contribution of each population - which we sum together into regions - to the admixture event and thus provide further intuition into historical gene flow.

#### 3.11 Defining GLOBETROTTER admixture events

The GLOBETROTTER algorithm provides multiple metrics as evidence that admixture has taken place which are combined to arrive at an understanding of the nature of the observed admixture event. In particular, as the authors suggest, to generate an admixture *P* value, we ran GLOBETROTTER’s “NULL” procedure, which estimates admixture parameters accounting for unusual patterns of LD, and then inferred 100 date bootstraps using this inference, identifying the proportion of inferred dates(s) that are ≤ 1 or ≥ 400.

Although the algorithm provides a “best-guess” for observed admixture event, we performed the following post-GLOBETROTTER filtering to arrive at our final characterisation of events. We outline the full GLOBETROTTER output in Figure 4-Source Data 2.

1. **Southern African populations** In all Southern African groups we present the results of the GLOBETROTTER runs where results are standardised by using the “NULL” individual see Hellenthal et al. [2014] for further details. We also note that in both the AmaXhosa and SEBantu GLOBETROTTER found evidence for two admixture events but on running the date bootstrap inference process, in both populations the most recent date confidence interval contained 1 generation, suggesting that the dating is not reliable. Inspection of the coancestry curves in this case showed that evidence for a single date of admixture.
2. **East Africa Afroasiatic speaking populations** In all Afroasiatic groups we present the results of the GLOBETROTTER runs where results are standardised by using a “NULL” individual see Hellenthal et al. [2014] for further details.
3. **West African Niger-Congo speaking populations** In all West African Niger-Congo speaking groups, with the exception of the Jola, GLOBETROTTER found evidence for two dates of admixture. In all cases the most recent event was young (1-10 generations) and the date bootstrap confidence interval often contained very small values. Inspection of the coancestry curves showed a sharp decrease at short genetic distances – consistent with the old inferred event – but there was little evidence of a more recent event based on these curves. In groups from this region we therefore show inference of a single date, which we take to be the older of the two dates inferred by GLOBETROTTER.

In all other cases we used the result output by GLOBETROTTER using the default approach.

#### 3.12 Comparison of weighted LD curve dates with GLOBETROTTER dates

Noting that there were differences between the dates inferred by the two dating methods we employed, we compared the dates generated by ALDER/MALDER with those inferred from GLOBETROTTER. Figure 3B shows a comparison of dates using the MALDER event inference; that is, for each population, we used the MALDER inference (either one or two dates) and used the corresponding GLOBETROTTER date inference (either one or two dates) irrespective of whether GLOBETROTTER’s inferred event was different to that of MALDER. Figure 3C is the opposite: we use GLOBETROTTER’s event inference to define whether we select one or two dates, and then use MALDER’s two date inferences if two dates are inferred, or ALDER’s inference if MALDER infers two dates and GLOBETROTTER infers one. Each point represents a comparison of dates for a single ethnic group, with the symbol and colour defining the ethnic group as in previous plots. In general there appears to be an inflation in dates inferred from West African groups (blues) with MALDER/ALDER compared to GLOBETROTTER.

#### 3.13 Analysis of admixture using sets of restricted surrogates

Recall that for GLOBETROTTER analyses two painting steps are required. One needs to (a) paint target individuals with a set of “donor” individuals to generate mosaic painted chromosomes, and (b) paint all potential “surrogate” groups with the same set of painting donors, such that we then describe admixture in the target individuals with this particular set of surrogate groups. One major benefit of GLOBETROTTER is its ability to represent admixing source groups as mixtures of surrogates.

To infer admixture in sub-Saharan African groups, we painted individuals with a set of painting donors which did not include other individuals from the same ancestry region. We refer to these as “non-local” donors. We then painted individuals from all ethnic groups with the same reduced set of painting donors, allowing us to include all ethnic groups as surrogates in the inference process. This allows us to infer sources of admixture for which the components can potentially come from any ethnic group. For example, when inferring admixture in the Jola from the West African Niger-Congo ancestry region, we painted all Jola individuals without including individuals from any other ethnic group from the West African Niger-Congo region. We then painted all ethnic groups with the same set of non-West African Niger-Congo donors, allowing us to model admixture using information from all ethnic groups which have been painted with the same set of painting donors.

The purpose of using this approach was to “mask” the effects of including many closely-related individuals in the painting process: including such individuals in a painting analysis causes the resultant mosaic chromosomes to be dominated by chunklengths from these local groups. As mentioned, this allows us to concentrate on identifying the ancestral processes that have occurred at broader spatio-temporal scales. In effect, removing closely-related individuals from the painting step means that we identify the changing identity of the non-local donor along chromosomes and use these identities to infer historical admixture processes.

#### 3.14 Removing non-local surrogates

In the main analysis we inferred admixture in each of the 48 target sub-Saharan African ethnic groups using all other 47 sub-Saharan African and 12 Eurasian groups as surrogates. We were interested in seeing how the admixture inference changed as we removed surrogate groups from the analysis. Masking surrogates like this provides further insight into the historical relationships between groups. By removing non-local surrogates, we can infer admixture parameters and characterize admixture sources as mixtures of this reduced set of surrogates. Given that, by definition, local groups are more closely related to the target of interest, this approach effectively asks who, outside of the targets region is next best at describing the sources of admixture.

We performed several “restricted surrogate” analyses, for different sets of targets, where we infer admixture using sub-sets of surrogates. One aim of the this analysis was to track the spread of Niger-Congo ancestry in the four Niger-Congo ancestry regions. For example, in the full analysis, the major sources of admixture in East African groups tended to be dominated by Southern African Niger-Congo (specifically Malawi) components. If we remove South African Niger-Congo groups from the admixture inference, how is the admixture source now composed?

We performed the following restricted surrogate analyses:

1. **No local region**: for all 48 African groups, we re-ran GLOBETROTTER without allowing any surrogates from the same ancestry region.
2. **No local, east or south**: for groups from the East African and South African Niger-Congo ancestry regions, we disallowed groups from both East and South African Niger-Congo regions from being surrogates. In effect, this asks where in West/Central African is their Niger-Congo ancestry likely to come from.
3. **No local or west**: For West and Central West African groups, we disallowed both West and Central West African Niger-Congo groups from being admixture surrogates. In effect, this asks where in East/South African their ancestry comes from.
4. **No local or Malawi**: As previously noted (Materials and Methods Section 2.5), Malawi was included in the South Africa Niger-Congo ancestry region. There is some evidence, for example from the fineSTRUCTURE analysis, that Malawi is closely related to the East African groups. We therefore wanted to assess whether the inference of a large amount of South African Niger-Congo ancestry in the major sources of admixture in East African Niger-Congo groups was a function of the genetic proximity of Malawi to East Africa. We removed East African Niger-Congo and Malawi as surrogates, and re-inferred admixture parameters.

#### 3.15 Summary of inferred gene-flow from West to East and South Africa

In Figure 4-figure supplement 1 we show the changing composition of GLOBETROTTER’s inferred sources of admixture. Figure 4-figure supplement 2 shows these same results with Niger-Congo surrogates only coloured, and with Malawi and Cameroon groups additionally emphasized. This analysis highlights several key insights:

1. **West and Central West Africa Niger-Congo**. Removing local surrogates from the West African Niger-Congo ancestry region does not substantially change the admixture source inference as admixture in groups from this region already involved Central West African donors. Sources of admixture in Central West African are replaced by West African Niger-Congo components in the non-local analysis. In both regions, admixture source components contain limited ancestry from East and South Africa regions. Interestingly, when we remove all West and Central West African surrogates, we observe similar major sources of admixture across groups from this region, which contains a significant amount of Nilo-Saharan / Afroasiatic ancestry (specifically Sudanese). We do not have Nilo-Saharan populations from Central and West Africa, although these exist. This observation may represent gene-flow between Nilo-Saharans populations across sub-Saharan Africa.
2. **East African Niger-Congo**. The major sources of admixture inferred in groups from this region tend to contain a majority of Southern African (Malawi) ancestry (Fig. 4-figure supplement 2) whether or not local surrogates are removed from the analysis. When we remove East African Niger-Congo groups and Malawi from the analysis, other South African Niger-Congo speaking groups (SEBantu, Herero, Amaxhosa) are involved in the admixture source. Subsequent removal of these groups leads to an admixture source that is almost exclusively made up of groups from Cameroon, suggesting that Niger-Congo ancestry in East and Southern Africa is most closely related to populations from this part of Central West Africa. Intriguingly, the remainder of major admixture source is made up of a small amount of Khoesan ancestry. Whilst we do not have autochtonous groups from East Africa, this component likely represents ancestry from groups that were present in the area before the major West to East population migrations.
3. **East African Nilo-Saharan and Afroasiatic**. In the full analysis the major source of admixture tends to contain local (purple) groups (Fig. 4-figure supplement 1). When we remove local surrogates from the analysis we see an increase in West and East African donors in the major sources of admixture.
4. **South African Niger-Congo**. We similarly observe mostly East African Niger-Congo ancestry in South African Niger-Congo speakers whether or not we remove non-local surrogates, although there is also a significant Malawi component in the SEBantu and Amaxhosa in the full analysis, which is replaced by East African Niger-Congo groups in the non-local analysis. As in East African Niger-Congo speakers, when we disallow any East or South African Niger-Congo surrogates, the major sources of admixture are almost exclusively from Cameroon donors.
5. **South African Khoesan**. Several Khoesan groups show evidence of specific admixture events involving Central West African donors, in the Khwe, /Gui//Gana, and Xun. When we remove Khoesan groups from being surrogates, interestingly we see that the major sources of admixture now tend to contain ancestry from Southern African Niger-Congo populations, which specifically are not Malawi-like (Fig. 4-figure supplement 2). We also observe a small amount of West African (dark blue) ancestry in these admixture sources, and very little Central West African ancestry (and none from Cameroon). This suggests that the non-Khoesan ancestry in groups from this region is unlikely to be associated with the migrations that spread the Cameroon (Bantu) ancestry into East and South Africa. We also observe a small amount of East African Nilo-Saharan / Afroasiatic ancestry in the major sources of admixture in the non-local analysis.

A key insight from these analyses is that a large amount of the ancestry in East and Southern Africa Niger-Congo speaking groups comes from populations currently present in Central West Africa. The opposite is not true for Western and Central West African groups, who share DNA amongst themselves, but when local surrogates are removed, they instead choose mainly non-Niger-Congo speaking groups as admixture sources.

### 4 Plotting date densities

For each admixture event we split the admixture sources into their constituent components (i.e. we used the *β* coefficients inferred by GLOBETROTTER) at the appropriate admixture proportions. For a given event, these components sum to 1. We multiplied these components by 100 to estimate the percentage of ancestry from a given event that originates from each donor group. We then assigned each of the components the set of date bootstraps associated with the event. For example, in the Kauma we infer an admixture event with an admixture proportion α of 6% involving a minor source containing the following coefficients: Massai 0.02 Afar 0.15 GBR 0.26 GIH 0.57. We multiply each of these coefficients by α to obtain a final proportion that each group gives to the admixture event: Massai 0.5 Afar 1 GBR 1.5 GIH 3. We assign all the inferred date bootstraps for the Kauma to each of the populations in these proportions. In this example, GIH has twice the density of GBR. We then additionally sum components across the same country to finally arrive at the density plots in Figures 5.

### 5 Gene-flow maps

We generated maps with the *rworldmaps* package in R. To generate arrows, we combined the inferred ancestral components (i.e. 1ST and 2ND EVENT SOURCES in Figure 3) for each population and estimated the proportion of a group’s ancestry coming from each component, summed across all surrogates from a particular country. For example, if an admixture contained source contains components from both the Jola and Wollof (both from The Gambia), then these components were added together. As such, the arrows point from the country of component origin to the country of the recipient. We then plot only those arrows which relate to events pertaining to the different broad gene-flow events. For each map, we used plot arrows for any event involving the following:

A. **Recent Western Bantu gene-flow**: any admixture source which has a component from either of the two Cameroon ethnic groups, Bantu and Semi-Bantu.
B. **Eastern Bantu gene-flow**: any admixture source which has a component from Kenya, Tanzania, Malawi, or South Africa (Niger-Congo speakers).
C. **East / West gene-flow**: any admixture event which has a component from Gambia, Burkina Faso, Ghana, Mali, Nigeria, Ethiopia, Sudan or Somalia.
D. **Eurasian gene-flow into Africa**: any admixture event which has a component from any Eurasian population.

An alternative map stratified by time window, rather than admixture component is shown in Figure 6-figure supplement 1.

### 6 Analysis and plotting code

Code used for analyses and plotting will be made available at https://github.com/georgebusby/popgen

## Acknowledgements

We thank all the MalariaGEN study sites that contributed samples to this analysis: a list of researchers involved at each study site can be found at https://www.malariagen.net/projects/host/consortium-members.

MalariaGEN is funded by the Wellcome Trust (WT077383/Z/05/Z, 090770/Z/09/Z) and the Bill and Melinda Gates Foundation through the Foundation for the National Institutes of Health (566). Genotyping was performed at the Wellcome Trust Sanger Institute, partly funded by its core award from the Wellcome Trust (098051/Z/05/Z). This research was also supported by Centre grants from the Wellcome Trust (090532/Z/09/Z) and the Medical Research Council (G0600718). C.C.A.S. was supported by a Wellcome Trust Career Development Fellowship (097364/Z/11/Z).

The Malaria Research and Training Center–Bandiagara Malaria Project (MRTC-BMP) in Mali group is supported by an Interagency Committee on Disability Research (ICDR) grant from the National Institute of Allergy and Infectious Diseases/US National Institutes of Health (NIAID/NIH) to the University of Maryland and the University of Bamako (USTTB) and by the Mali-NIAID/NIH International Centers for Excellence in Research (ICER) at USTTB. The Kenya Medical Research Institute (KEMRI)–Wellcome Trust Programme is funded through core support from the Wellcome Trust. This paper is published with the permission of the director of KEMRI. C.M.N. is supported through a strategic award to the KEMRI–Wellcome Trust Programme from the Wellcome Trust (084538). The Joint Malaria Programme, Kilimanjaro Christian Medical Centre in Tanzania received funding from a UK MRC grant (G9901439).

We thank Clare Bycroft, Lucy van Dorp, and Cristian Capelli for critically evaluating the manuscript and Francesco Montinaro for insightful discussions on interpretation of MALDER analyses. Genotype data for the MalariaGen samples included in this paper will be made available at the European Nucleotide Archive (accession number TBC).

## Figures and Figure Supplements

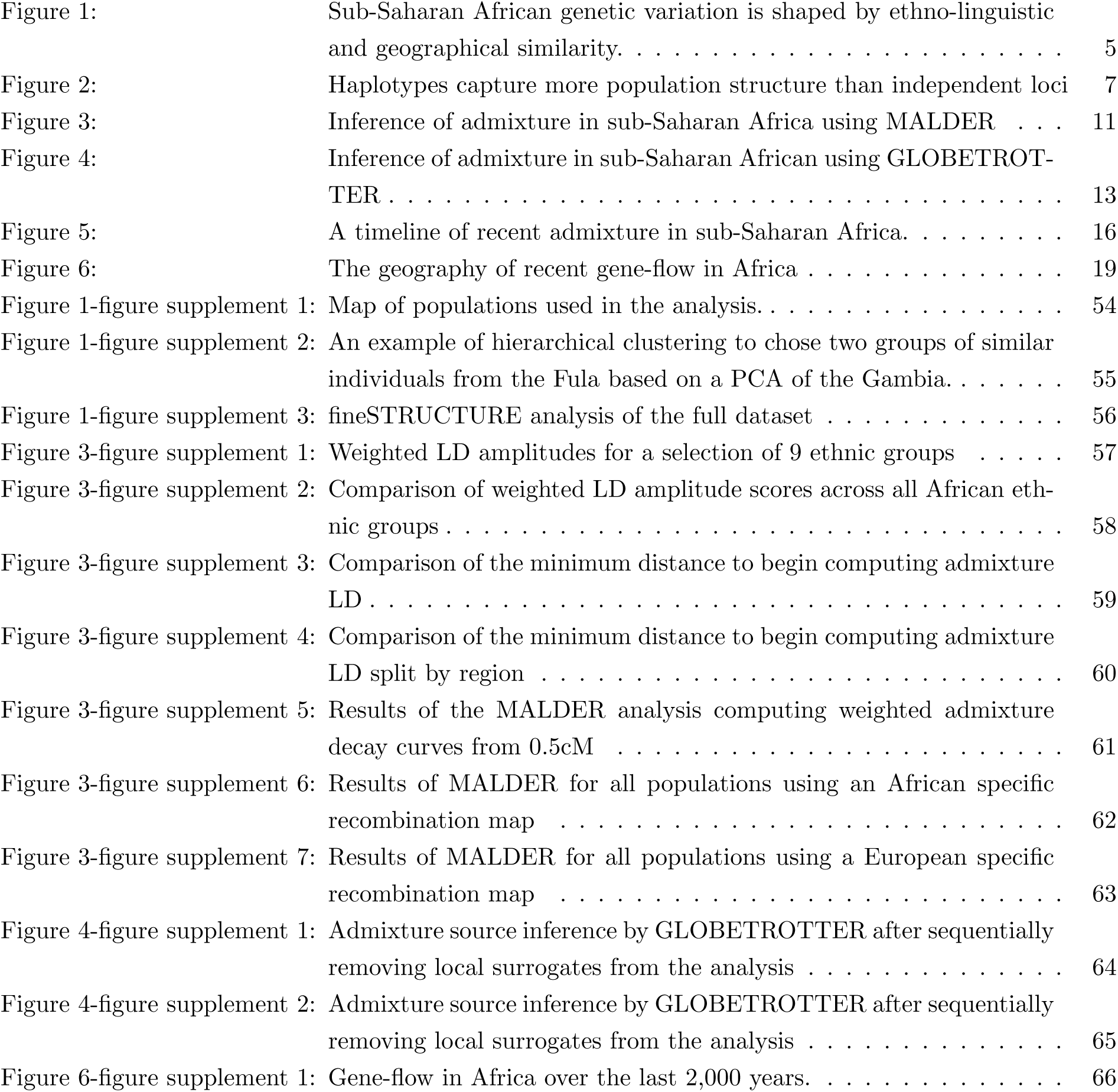

### Supplementary Files

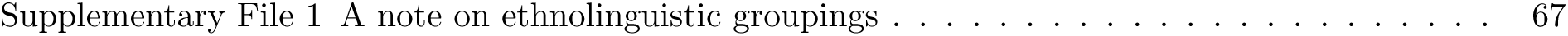

|  |  |
| --- | --- |
| Supplementary File 1 A note on ethnolinguistic groupings . . . . . | 67 |

### Source Data Tables

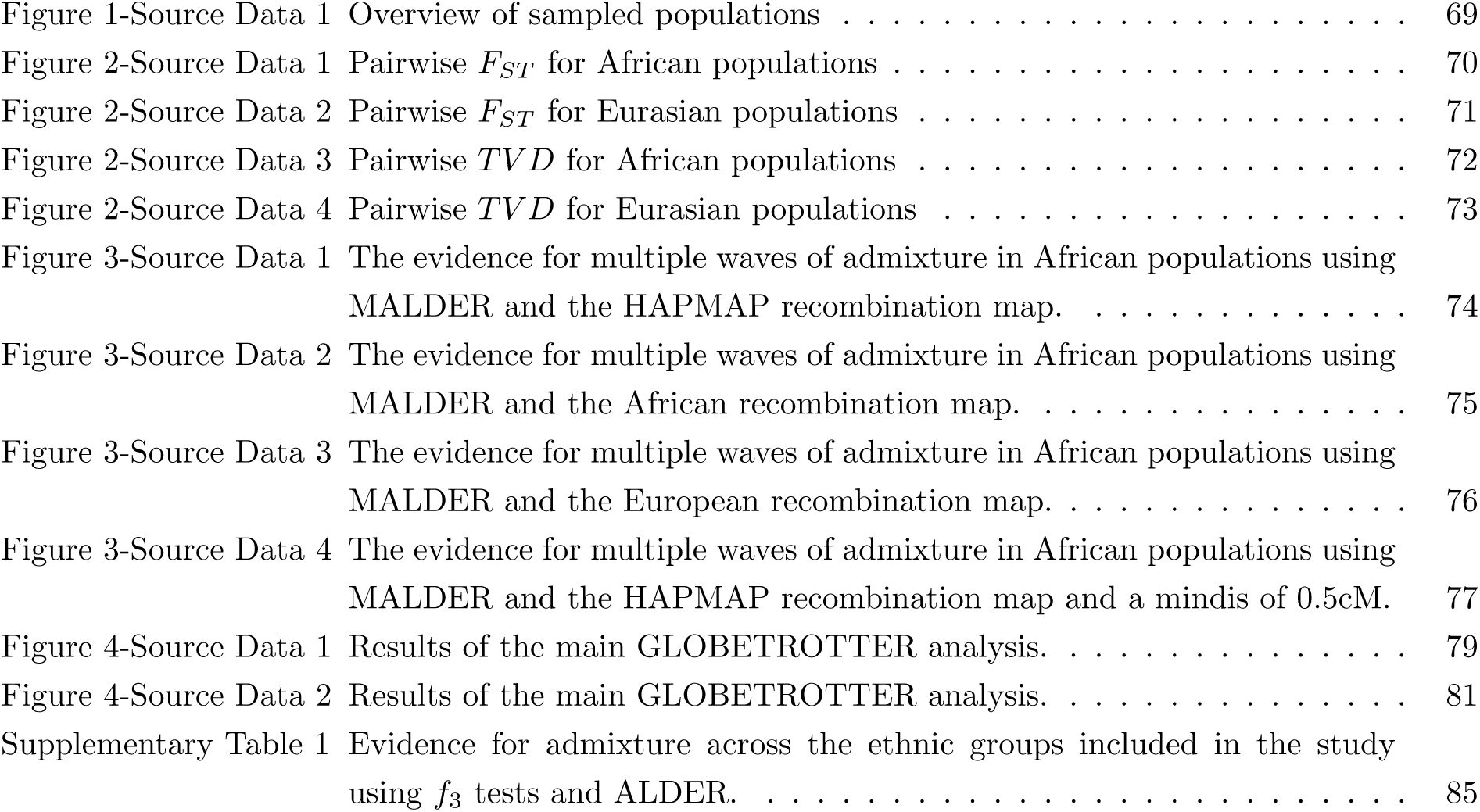

### Figure Supplements

**Figure 1-figure supplement 1.**
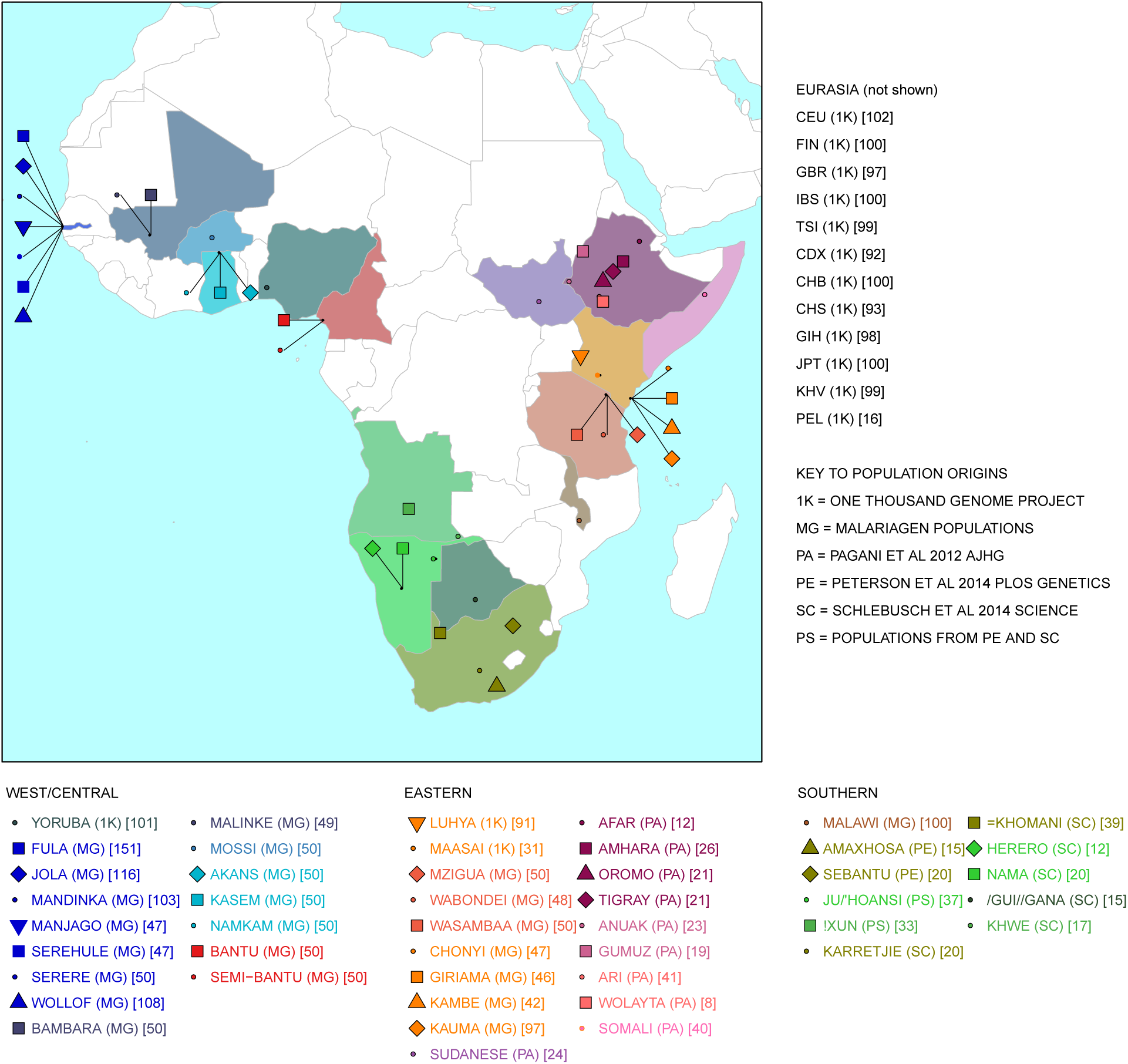
Map of populations used in the analysis. Population names are coloured by the country of origin; positions of the countries are shown on the map. Individual point labels, which are used throughout this paper, are shown for each population in the legend. Sample provenance is shown immediately after the population name in circular parentheses and final number of individuals is shown in square parentheses.

**Figure 1-figure supplement 2.**
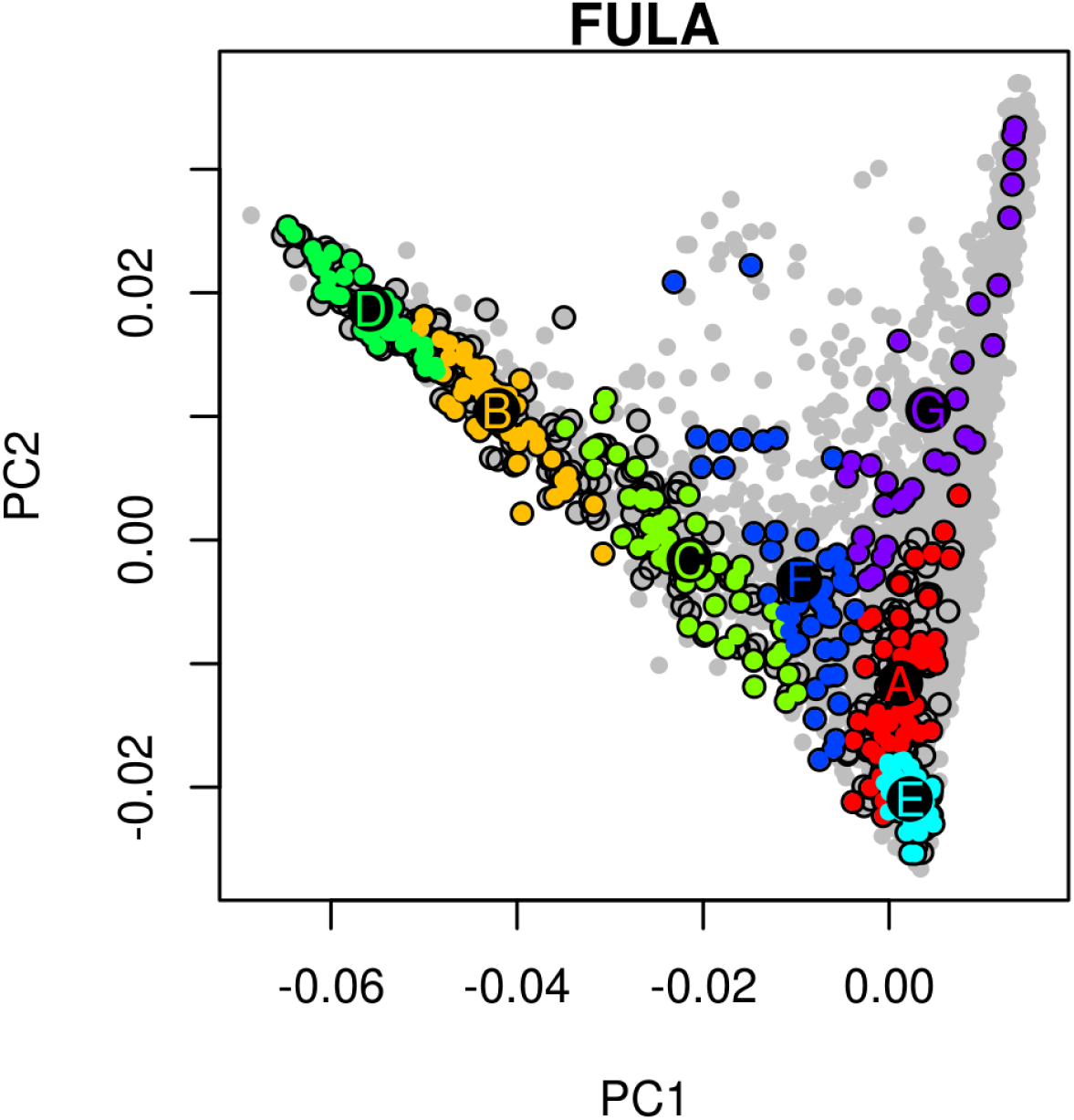
An example of hierarchical clustering to chose two groups of similar individuals from the Fula based on a PCA of the Gambia. Projected onto a PCA of Gambian genetic variation where each point represents an individual, all Fula individuals are coloured, with the colour depicting their cluster assignment, based on the MClust clustering algorithm. We chose individuals from the green (D) and light blue (E) clusters to maximise the representation of Fula genetic variation. Note that the majority of the individuals from the other 6 Gambian ethnic groups occur in the right arm of the PCA. An analogous process was preformed for all ethnic groups from the MalariaGEN dataset where more than 50 individuals were available.

**Figure 1-figure supplement 3.**
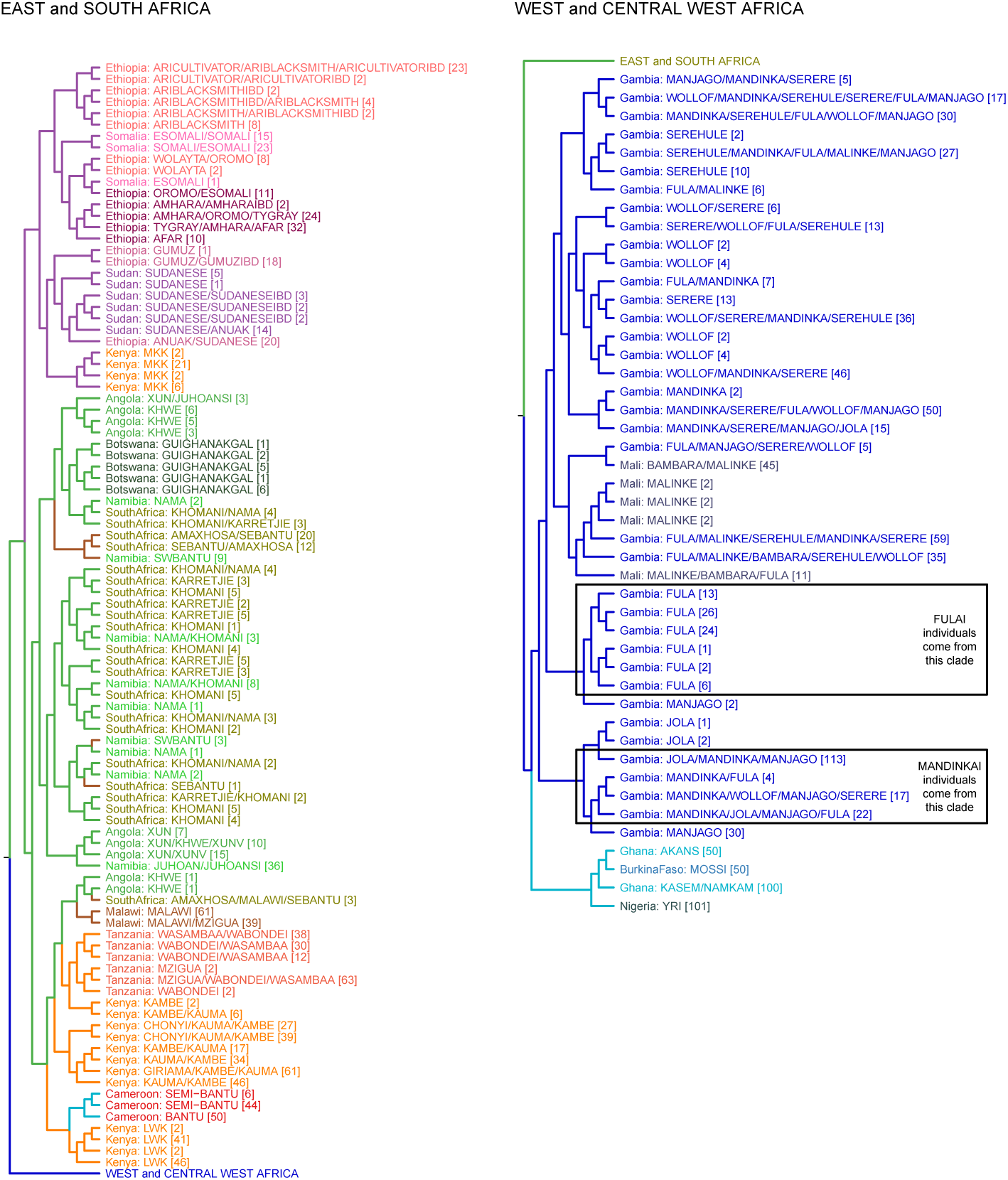
fineSTRUCTURE analysis of the full dataset. We show the tree output from a single run of the fineSTRUCTURE algorithm. To aid reading, the tree has been split in two, East and Southern African groups are on the left, West and Central West African groups are on the right. Leaves are labelled by the identity of the individuals within them, with the total number of individuals in the clusters shown in parentheses. Leaves are coloured by the country of origin (as in Fig. 1-figure supplement 1) and branches are coloured by the final ancestry region that the clusters were assigned to. Note that although Malawi and Cameroon individuals were located in a clade with mostly East African individuals, they were assigned to Southern and Central West African ancestry regions, respectively. Clades containing outlying individuals from the Fula and Mandinka are also shown.

**Figure 3-figure supplement 1.**
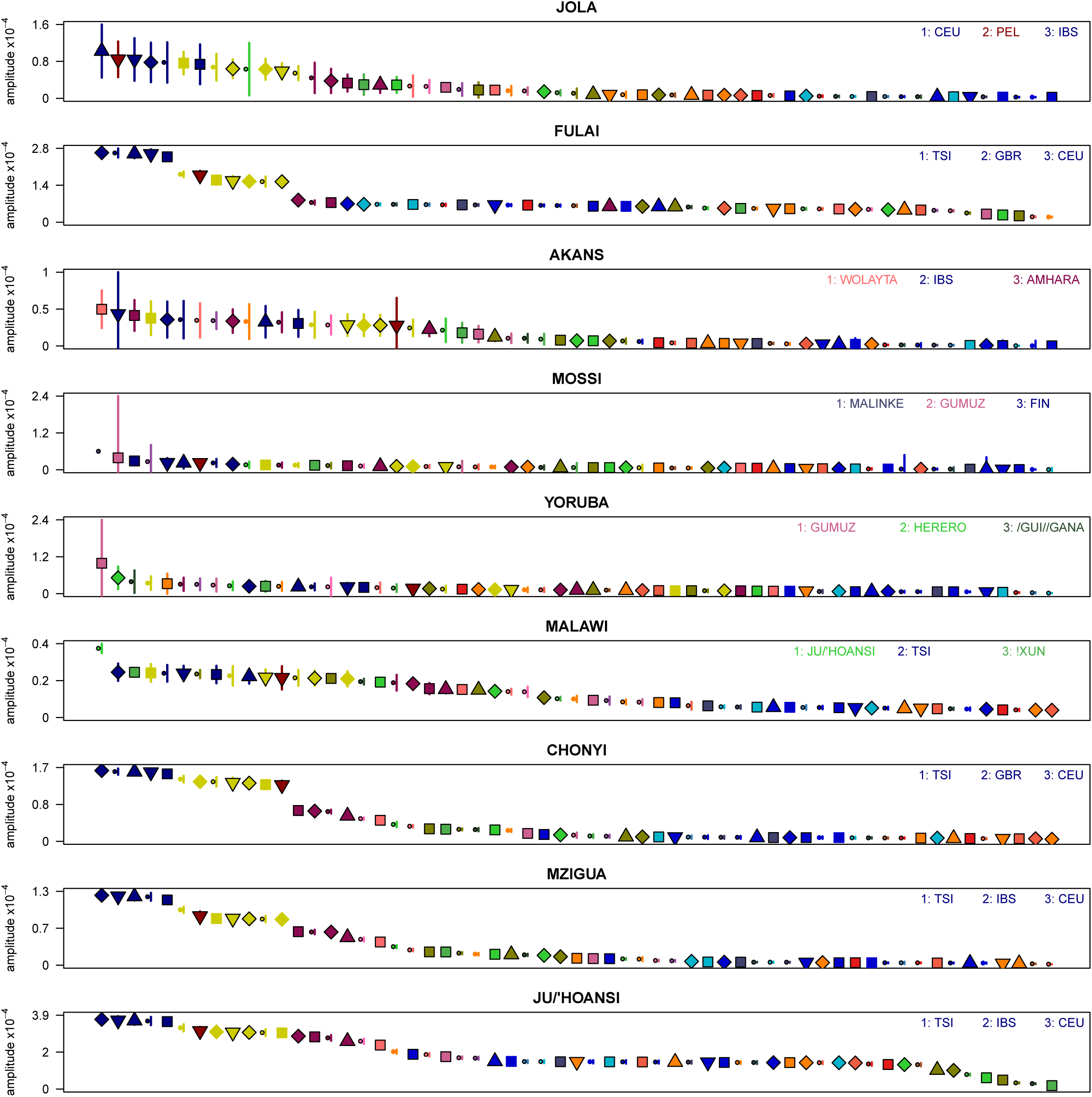
Weighted LD amplitudes for a selection of 9 ethnic groups. For a given test population we show the amplitude (± 1 s.e.) computed using a test population and every other population as the second reference. Plotted are the fitted amplitudes for each set of curves with the population used labelled beneath, with populations ordered by amplitude. A large number of population showed a similar profile to (A), that is with Eurasian populations showing the highest amplitudes. Other populations, e.g. Malawi, obtained the largest amplitudes from an African population.

**Figure 3-figure supplement 2.**
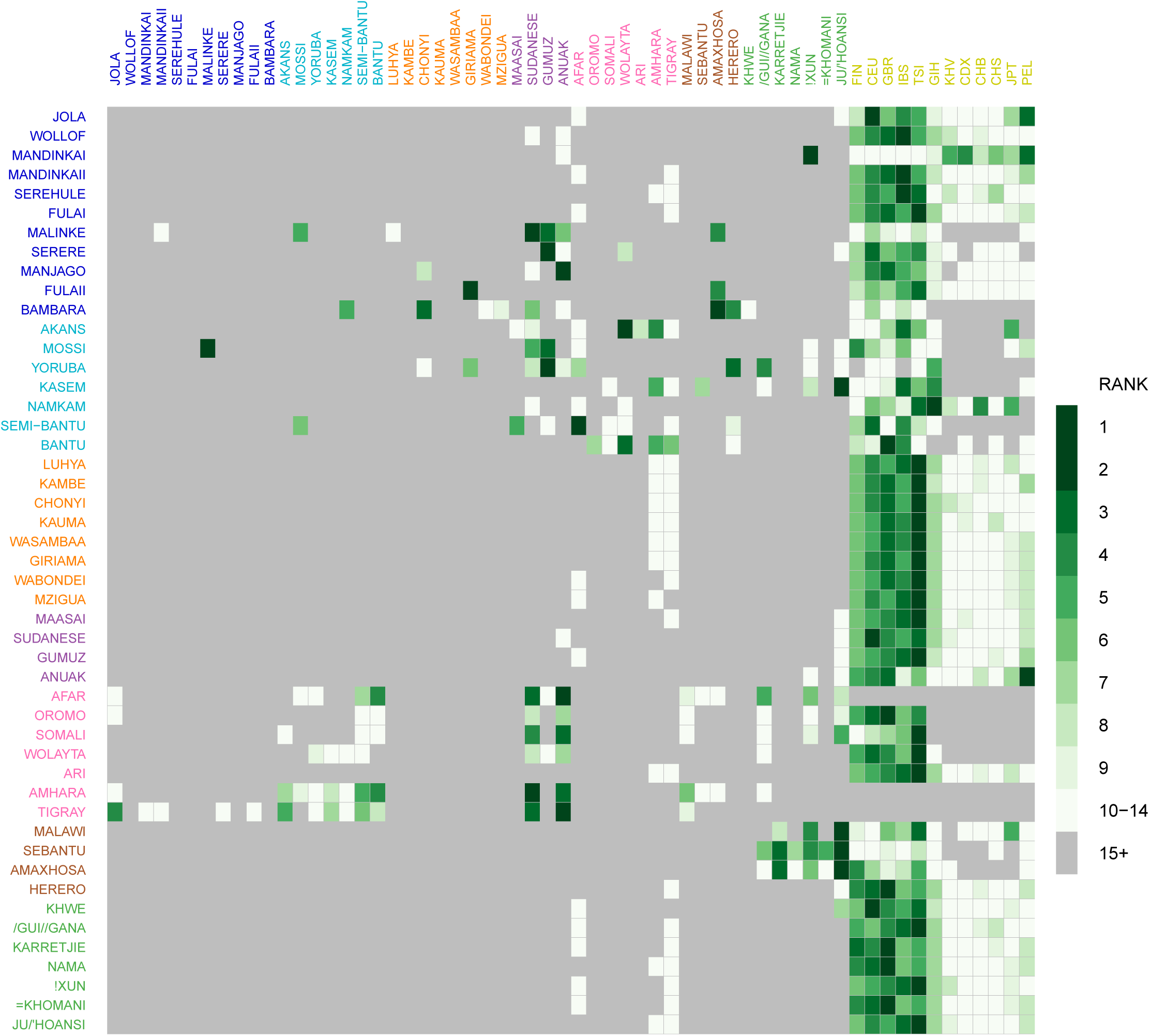
Comparison of weighted LD amplitude scores across all African ethnic groups. For a given test population we computed the ALDER amplitude (y-axis intercept) using the test population and every other population as the second reference. We then ranked the amplitudes across a given test population: populations who gave the top-ranked (i.e. largest) amplitude are in green, with those beneath a rank 15 shown in grey. This analysis shows that for many populations the reference populations giving the largest amplitudes (i.e. have the highest rank) are often non-African groups.

**Figure 3-figure supplement 3.**
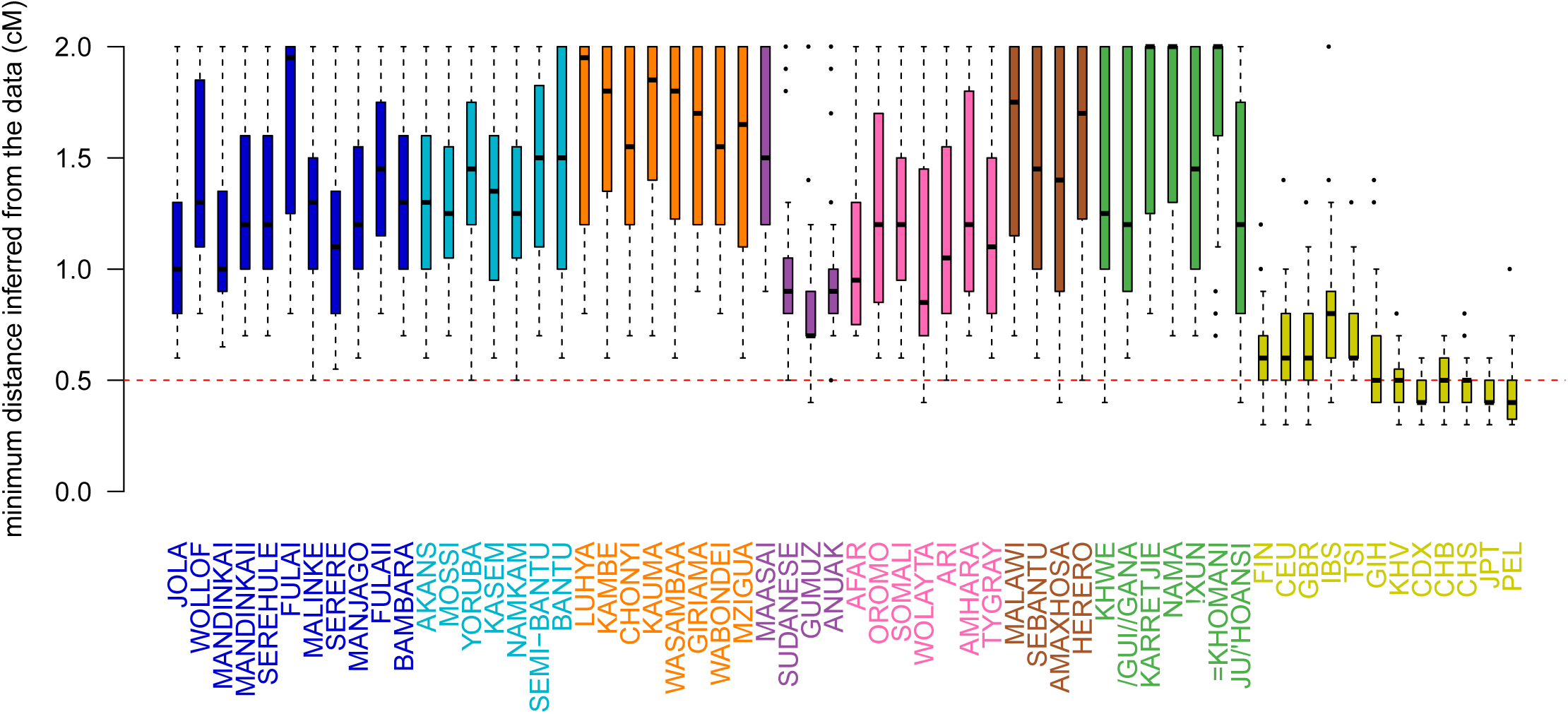
Comparison of the minimum distance to begin computing admixture LD. For each of the 48 African populations as a target, we used ALDER to compute the minimum distance over which short-range LD is shared with each of the 47 other African and 12 Eurasian reference populations. Here we show boxplots showing the distribution of minimum inferred genetic distances (y-axis) over which LD is shared for each of the reference populations separately (x-axis). We performed two analyses using weighted LD, one using these values of the minimum distance inferred from the data, and another where this distance was forced to be 0.5cM (dotted red line). The comparisons show that all African populations share LD correlations at distances > 0.5cM with all other African populations. Note that ALDER computes LD correlations at distances < 2cM.

**Figure 3-figure supplement 4.**
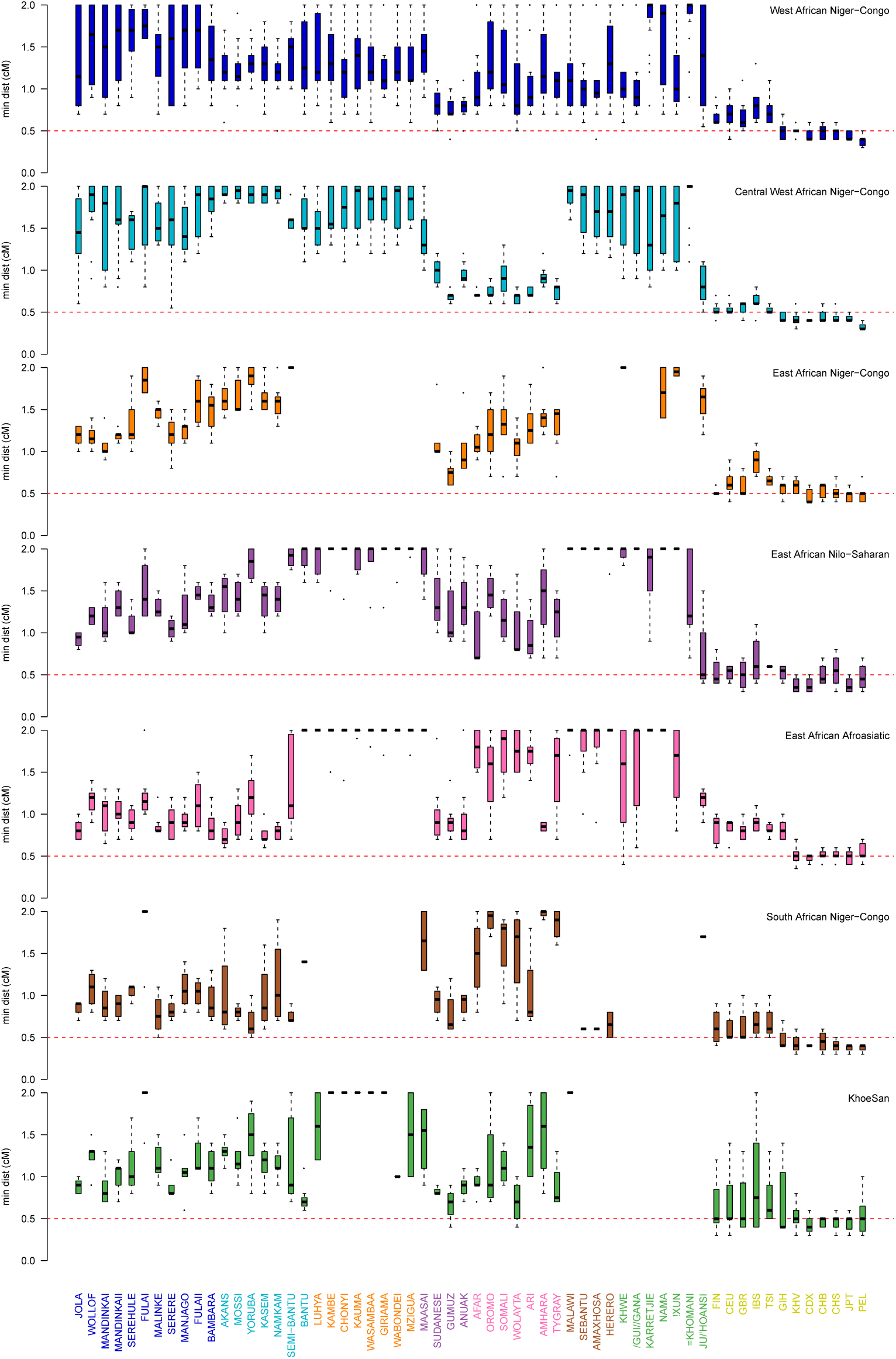
Comparison of the minimum distance to begin computing admixture LD split by region. As in Figure 3-figure supplement 3 except distances are stratified by region. The comparisons show that all African populations share LD correlations at distances > 0.5cM with all other African populations. Note that ALDER computes LD correlations at distances < 2cM.

**Figure 3-figure supplement 5.**
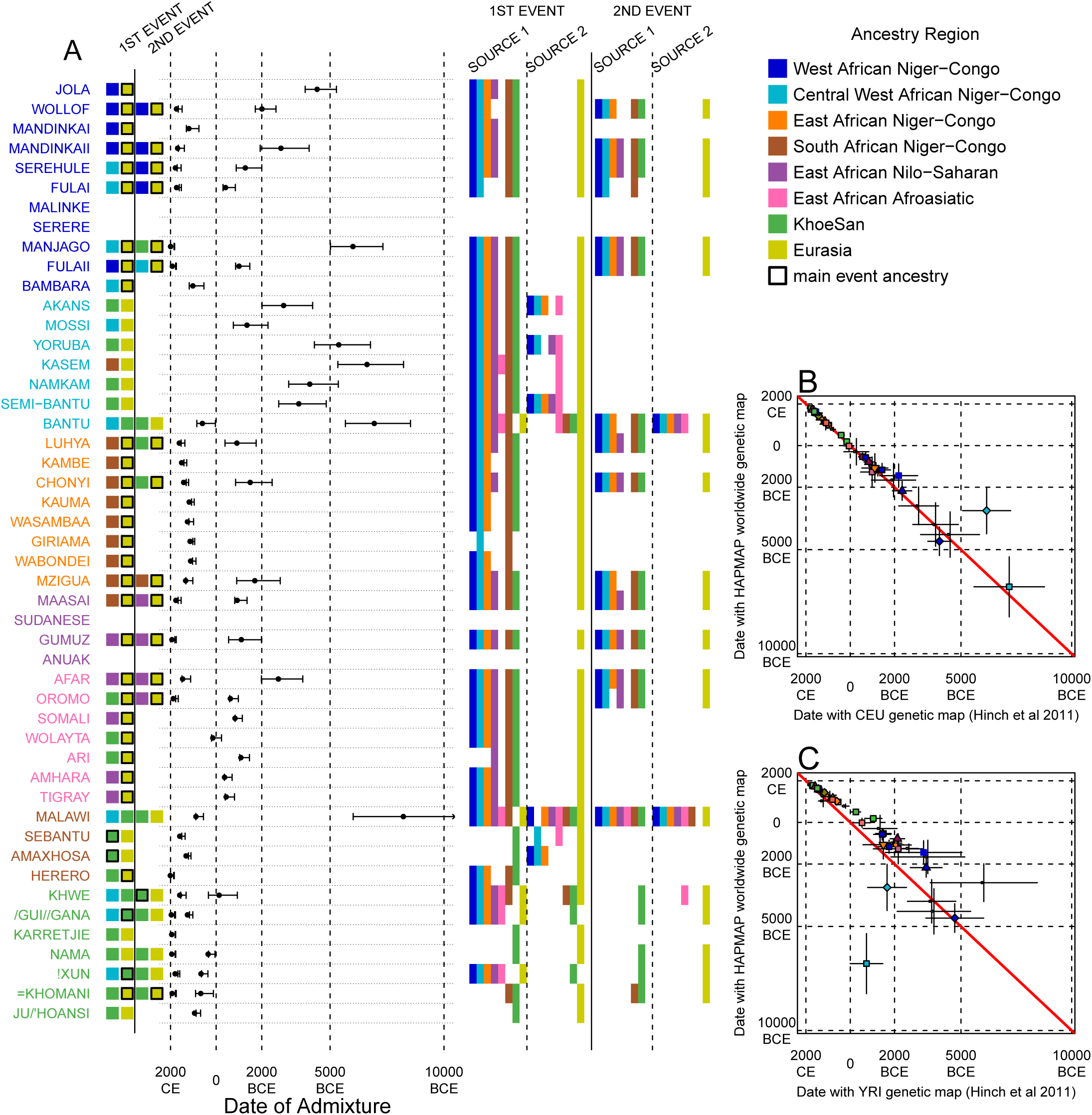
Results of the MALDER analysis computing weighted admixture decay curves from 0.5cM. As in the main analyses, the algorithm was run independently three times with the HAPMAP, YRI, and CEU genetic maps. The main results shown here are from the HAPMAP analysis. For each population, we show the ancestry region identity of the two populations involved in generating the MALDER curves with the greatest amplitudes (which are the closest to the true admixing sources amongst the reference populations) for at most two events. The sources generating the greatest amplitude are highlighted with a black box. Populations are ordered by ancestry of the admixture sources and dates estimates which are shown ± 1 s.e. (B) Comparison of dates of admixture ± 1 s.e. for MALDER dates inferred using the HAPMAP recombination map and a recombination map inferred from European (CEU) individuals from Hinch et al. [2011]. We only show comparisons for dates where the same number of events were inferred using both methods. Point symbols refer to populations and are as in Figure 1. (C) as (B) but comparing with an African (YRI) map.

**Figure 3-figure supplement 6.**
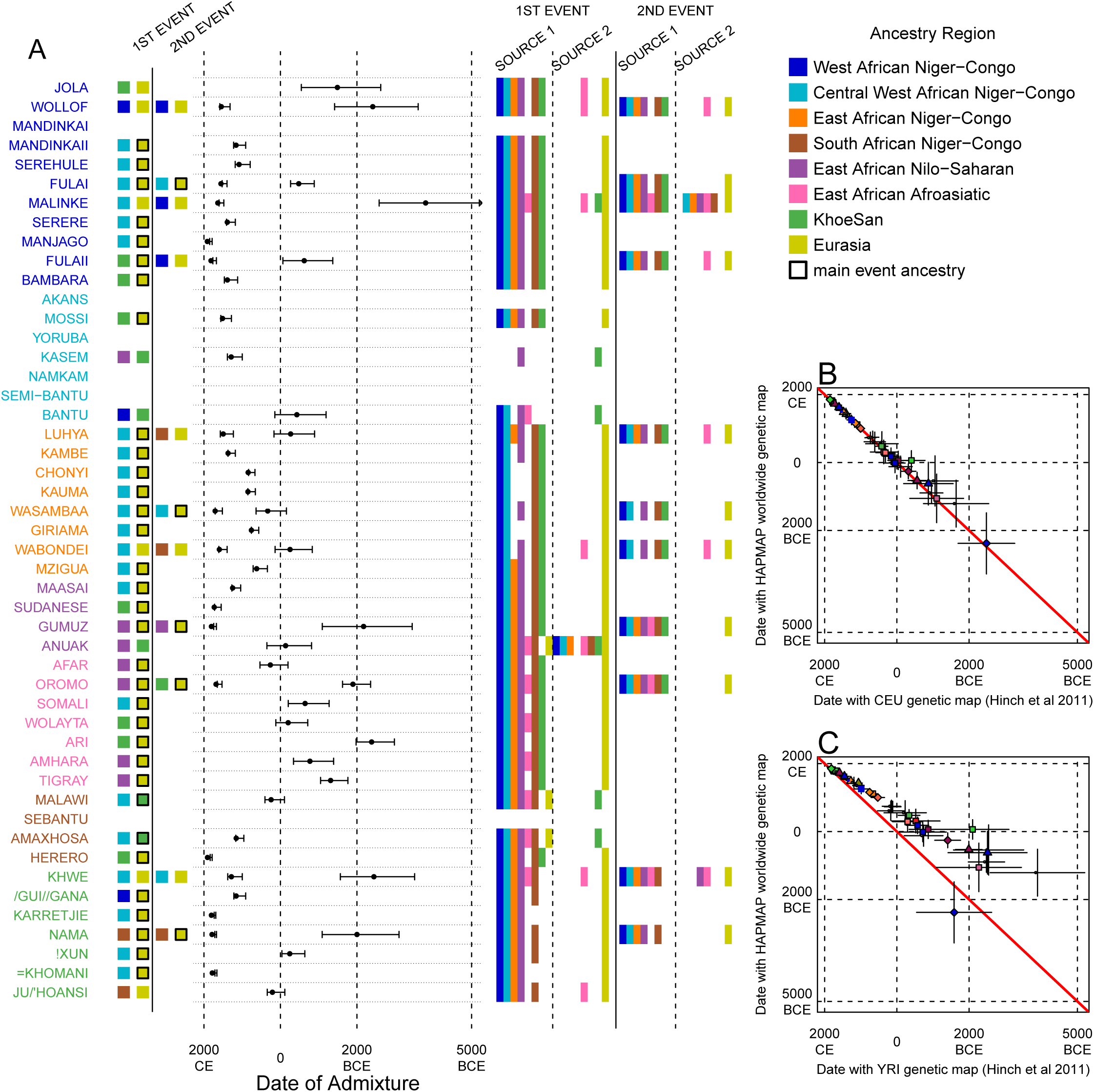
Results of MALDER for all populations using an African specific recombination map. We used MALDER to identify the evidence for multiple waves of admixture in each population. (A) For each population, we show the ancestry region identity of the two populations involved in generating the MALDER curves with the greatest amplitudes (which are the closest to the true admixing sources amongst the reference populations) for at most two events. The sources generating the greatest amplitude are highlighted with a black box. Populations are ordered by ancestry of the admixture sources and dates estimates which are shown ± 1 s.e. (B) Comparison of dates of admixture ± 1 s.e. for MALDER dates inferred using the HAPMAP recombination map and a recombination map inferred from European (CEU) individuals from Hinch et al. [2011]. We only show comparisons for dates where the same number of events were inferred using both methods. Point symbols refer to populations and are as in Figure 1. (C) as (B) but comparing with an African (YRI) map.

**Figure 3-figure supplement 7.**
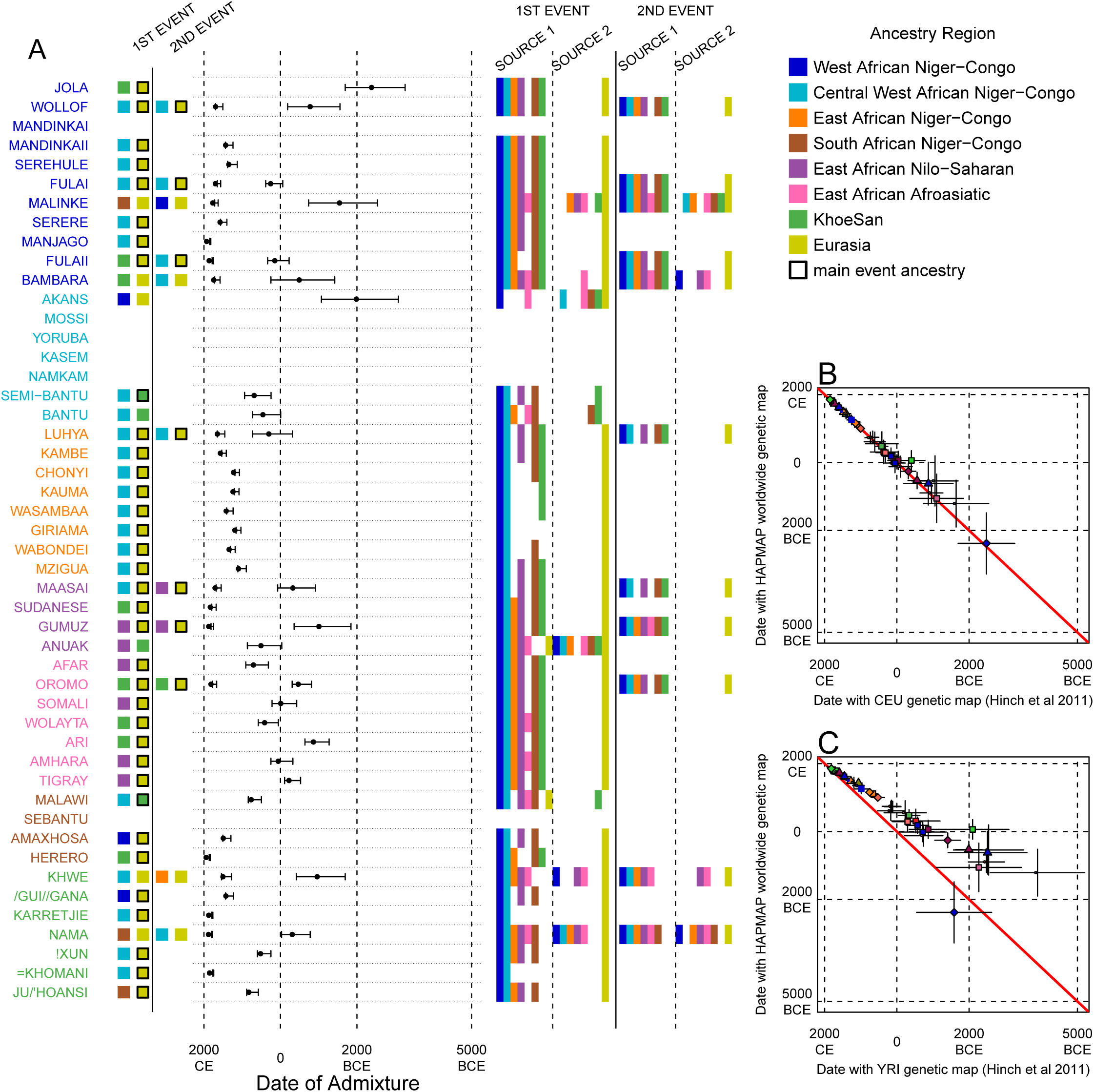
Results of MALDER for all populations using a European specific recombination map. We used MALDER to identify the evidence for multiple waves of admixture in each population. (A) For each population, we show the ancestry region identity of the two populations involved in generating the MALDER curves with the greatest amplitudes (which are the closest to the true admixing sources amongst the reference populations) for at most two events. The sources generating the greatest amplitude are highlighted with a black box. Populations are ordered by ancestry of the admixture sources and dates estimates which are shown ± 1 s.e. (B) Comparison of dates of admixture ± 1 s.e. for MALDER dates inferred using the HAPMAP recombination map and a recombination map inferred from European (CEU) individuals from Hinch et al. [2011]. We only show comparisons for dates where the same number of events were inferred using both methods. Point symbols refer to populations and are as in Figure 1. (C) as (B) but comparing with an African (YRI) map.

**Figure 4-figure supplement 1.**
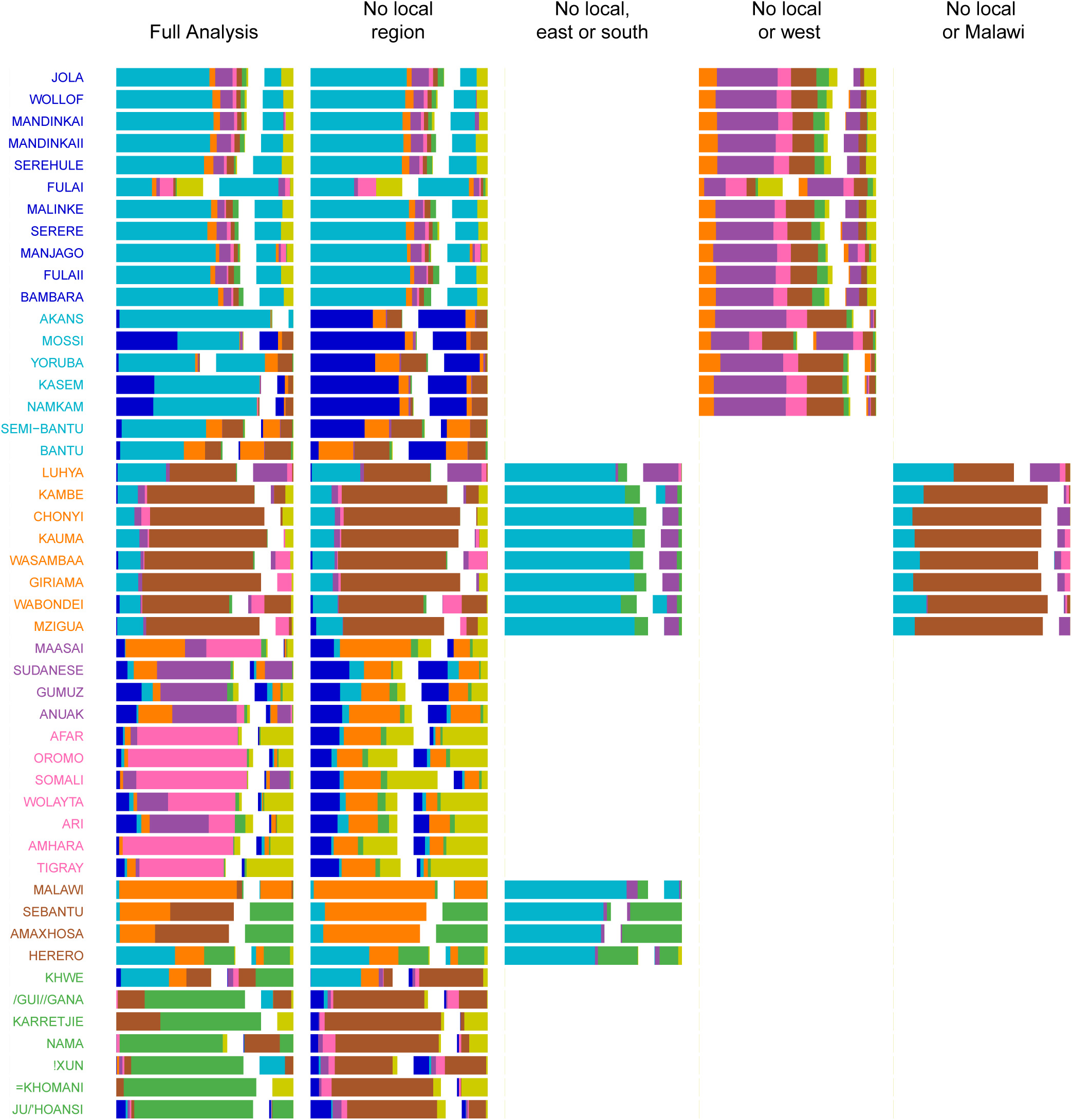
Admixture source inference by GLOBETROTTER after sequentially removing local surrogates from the analysis. In addition to the Full analysis, we show the inferred composition of admixture sources for different, restricted surrogate analyses. Components and y-axis labels are coloured by ancestry region. In each case we show admixture sources inferred by GLOBETROTTER for a single date of admixture.

**Figure 4-figure supplement 2.**
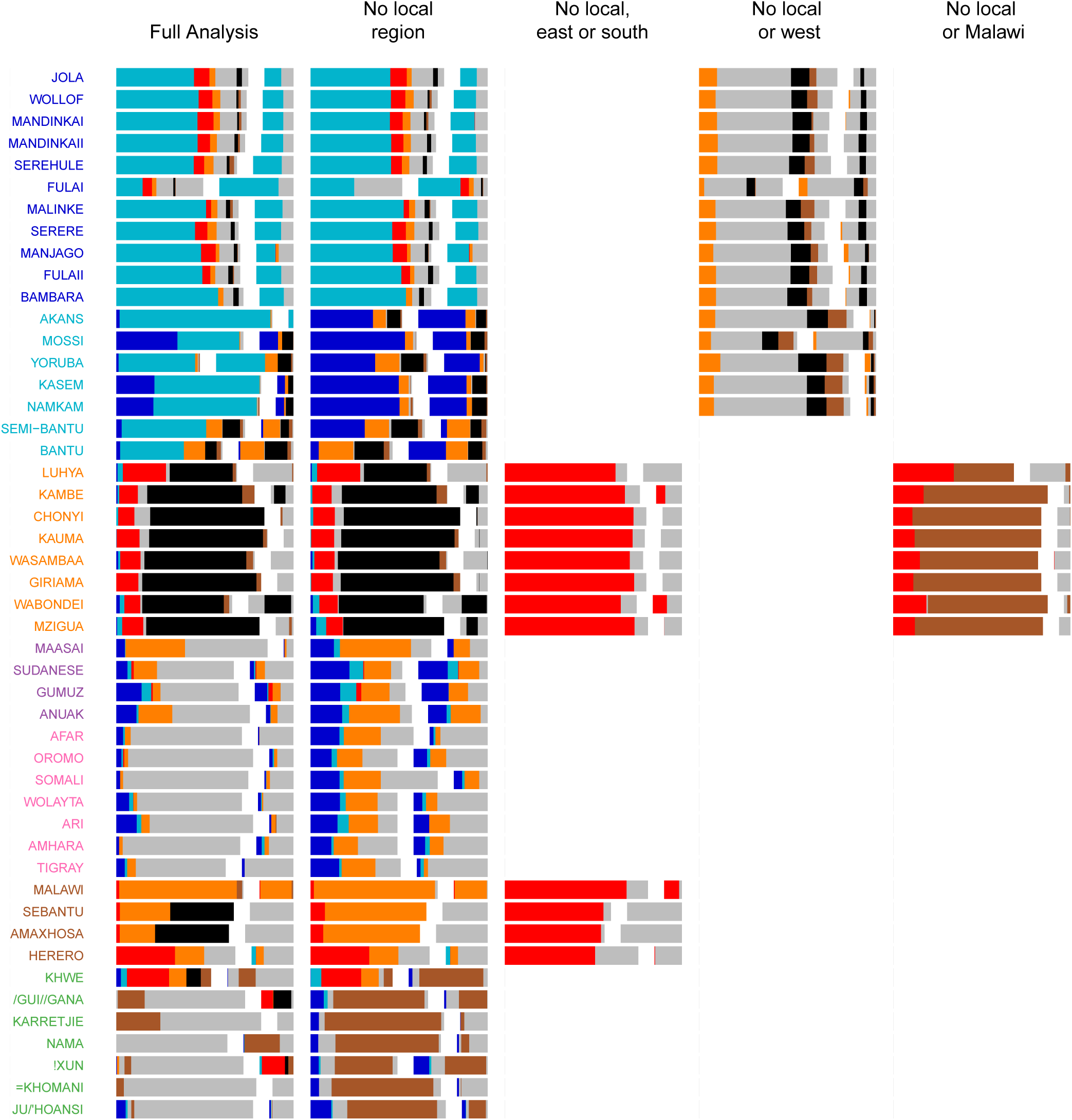
Admixture source inference by GLOBETROTTER after sequentially removing local surrogates from the analysis. The results are the same as Figure 4-figure supplement 1, but only Niger-Congo speaking groups are coloured. We highlight Malawi components in black, and Cameroon (Bantu and Semi-Bantu) in red.

**Figure 6-figure supplement 1.**
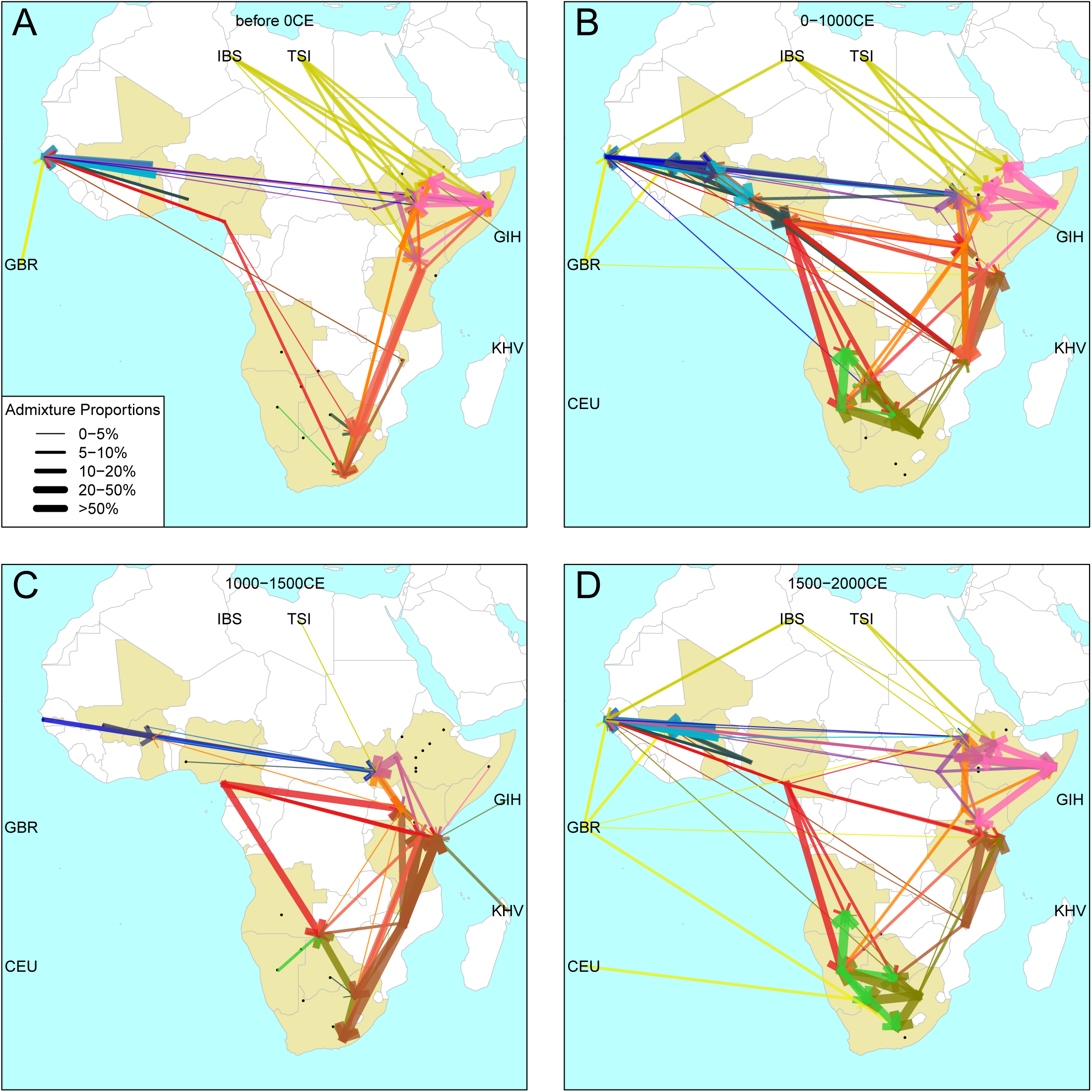
Gene-flow in Africa over the last 2,000 years. Using the results of the GLOBETROTTER analysis we show the connections between different groups in sub-Saharan Africa over time. For each population, we inferred the date of admixture and the composition of the admixing sources. We link each recipient population to its donor components using arrows, the size of which is proportional to the amount it contributes to the admixture event. Arrows are coloured by country of origin, as in Figure 4 in the main text.

### Supplementary File 1 A note on ethnolinguistic groupings

The results of the population genetic analysis shows that population structure is largely the result of ethno-linguistic similarity, which itself is largely but not completely correlated with geographical proximity. These divisions are shown below and referred to in the text, together with the latest Ethnologue classification^‡^ of the languages spoken, where possible.

1. 1st major Niger-Congo speaking group from West Africa: Gambian and Malian ethnic groups

- Niger-Congo, **Mande** {Mandinka, Malinke, Bambara}
- Niger-Congo, Atlantic-Congo, Atlantic, **Northern, Senegambian, Fula-Wolof** {Fula, Wollof}
- Niger-Congo, Atlantic-Congo, Atlantic, **Northern, Senegambian, Serer** {Serere, Serehule}
- Niger-Congo, Atlantic-Congo, Atlantic, **Northern, Bak** {Jola}
2. 2nd major Niger-Congo speaking group from West Africa: Ghana/BF/Nigerian ethnic groups:

- Niger-Congo, Atlantic-Congo, Atlantic, **Volta-Congo**, Kwa{Akan, Yoruba}
- Niger-Congo, Atlantic-Congo, Atlantic, **Volta-Congo, North, Gur** {Mossi, Kasem, Namkam?}
3. A Central and Eastern African Niger-Congo / “Bantoid” speaking group, split into two sub-divisions:

a. North Western

- Niger-Congo, Atlantic-Congo, Atlantic, Volta-Congo, **Benue-Congo, exNarrow-Bantu** {Cameroon: Bantu, Semi-Bantu?}
b. Eastern

- Niger-Congo, Atlantic-Congo, Atlantic, Volta-Congo, **Benue-Congo, Bantoid, Southern, Narrow-Bantu, Central, I** {Masaba-Luhya?}
- Niger-Congo, Atlantic-Congo, Atlantic, Volta-Congo, **Benue-Congo, Bantoid, Southern, Narrow-Bantu, Central, E** {Mijikenda (Kenya)}
- Niger-Congo, Atlantic-Congo, Atlantic, Volta-Congo, **Benue-Congo, Bantoid, Southern, Narrow-Bantu, Central, F-G** {Tanzania}
- Niger-Congo, Atlantic-Congo, Atlantic, Volta-Congo, **Benue-Congo, Bantoid, Southern, Narrow-Bantu, Central, N** {Malawi (Chewa)}
4. Afroasiatic and Nilo-Saharan speakers from the Horn of Africa:

- Afroasiatic, **Cushitic** {Afar, Somali, Oromo}
- Afroasiatic, **Semitic** {Amhara, Tigrayan}
- Afroasiatic, **Omotic** {Ari, Wolayta}
- Nilo-Saharan, **Komuz** {Gumuz}
- Nilo-Saharan, **Eastern Sudanic, Nilotic** {Maasai}
- Nilo-Saharan, **Eastern Sudanic, Nilotic** {Anuak}
- Nilo-Saharan {Sudanese}
5. Khoesan and Bantu speaking groups from Southern Africa

a. Khoesan

- Khoesan, Southern Africa, **Northern** {Ju/’hoansi, !Xun}
- Khoesan, Southern Africa, **Central** {Nama}
- Khoesan, Southern Africa, **Central, Tshu-Khwe, Northwest** {/Gui//Gana, Khwe}
- Khoesan, Southern Africa, **Southern** {=Khomani}
- uncertain {Karretijie}
b. Southern Bantu speakers

- Niger-Congo, Atlantic-Congo, Atlantic, Volta-Congo, **Benue-Congo, Bantoid, Southern, Narrow-Bantu, Central, R** {Herero}
- Niger-Congo, Atlantic-Congo, Atlantic, Volta-Congo, **Benue-Congo, Bantoid, Southern, Narrow-Bantu, Central, S** {Amaxhosa, SEBantu}

### Source Data

**Figure 1-Source Data 1.**
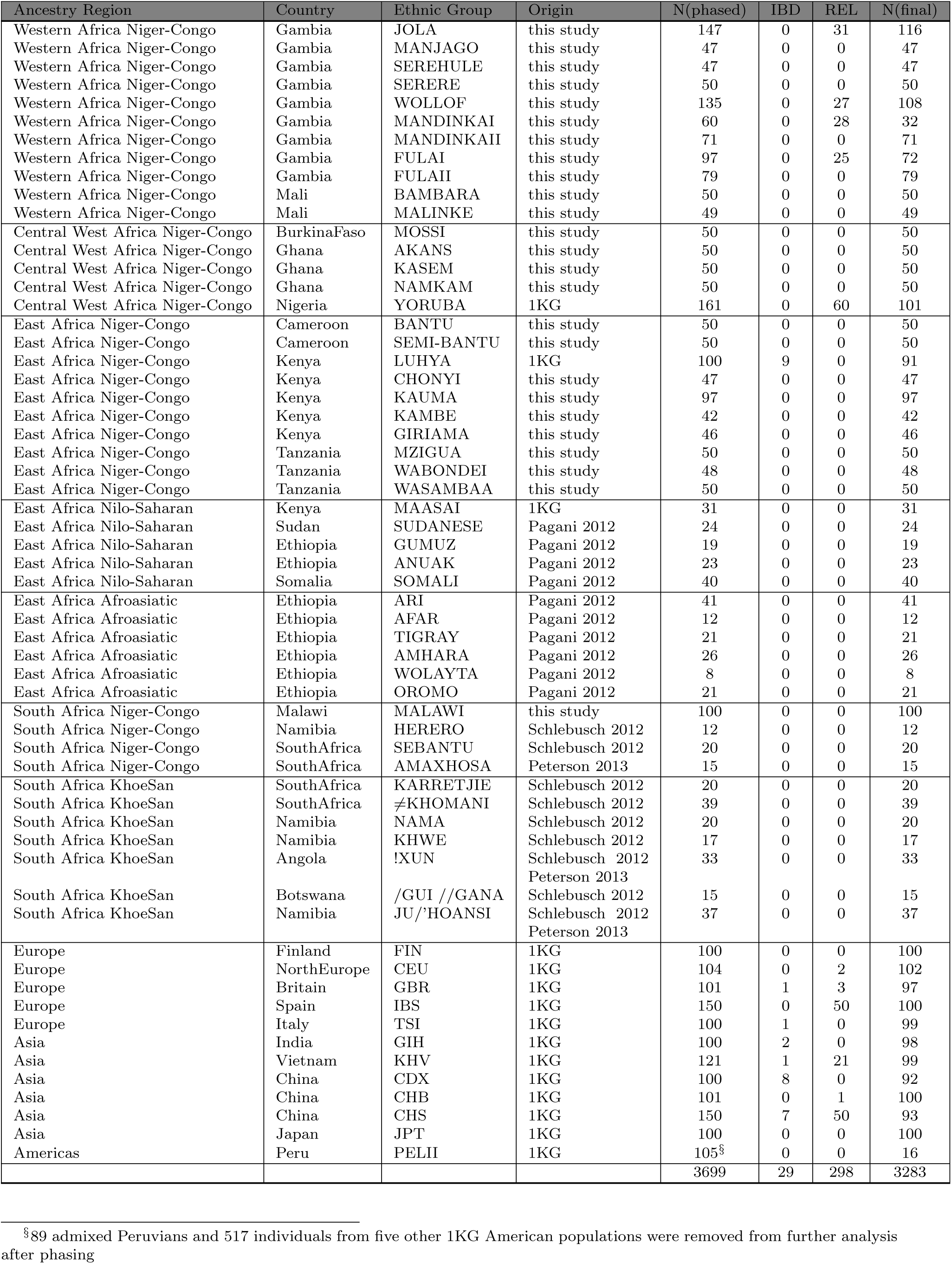
Overview of sampled populations describing the continent, region, numbers of individuals used, and the source of any previously published datasets.

**Figure 2-Source Data 1.**
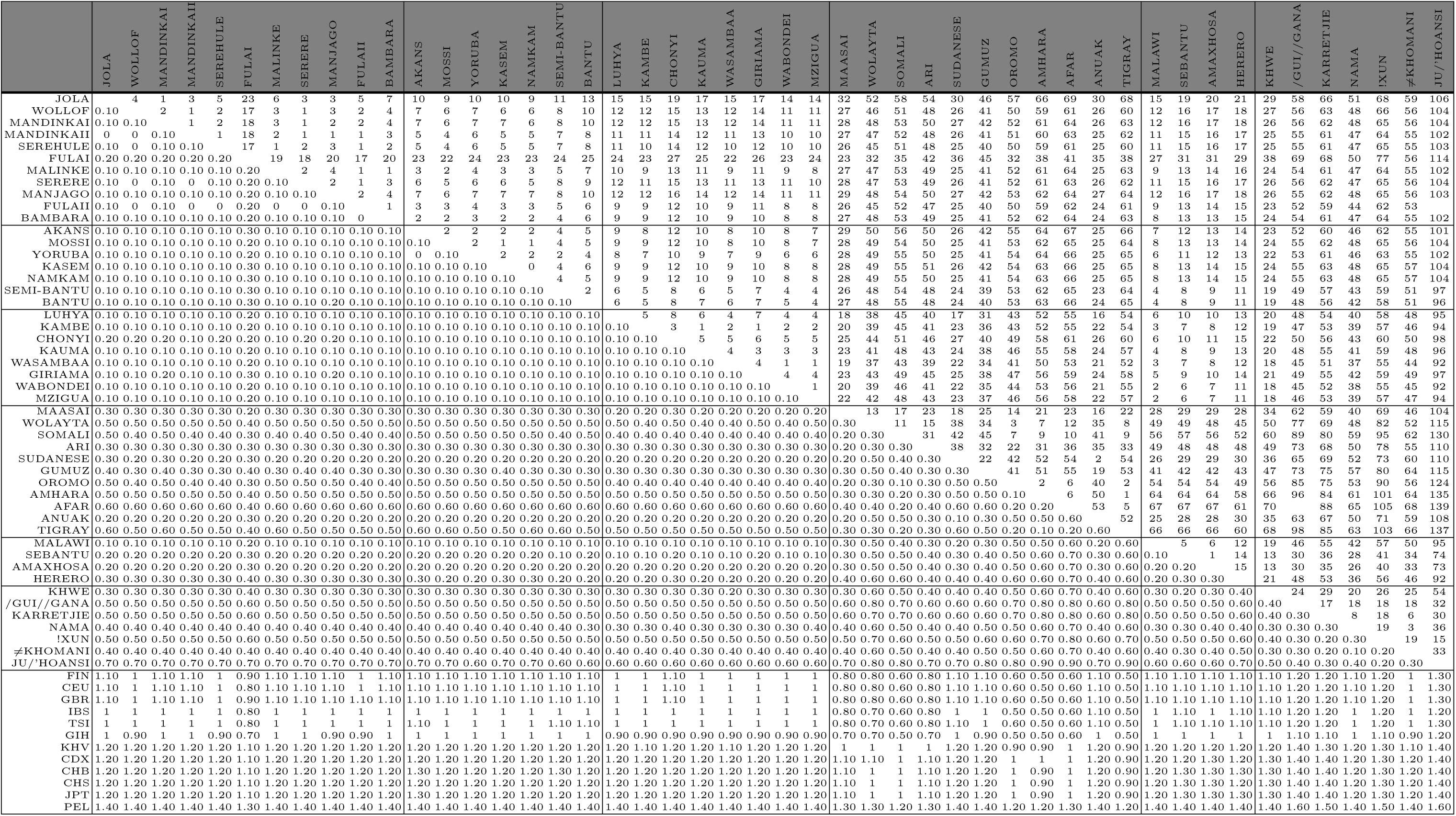
Pairwise *F_ST_* for African populations. We used *smartpca* to compute *F_ST_* for each pair of populations, upper left diagonal, together with standard errors computed using a block jacknife. *F_ST_* has been multiplied by 1000

**Figure 2-Source Data 2.**
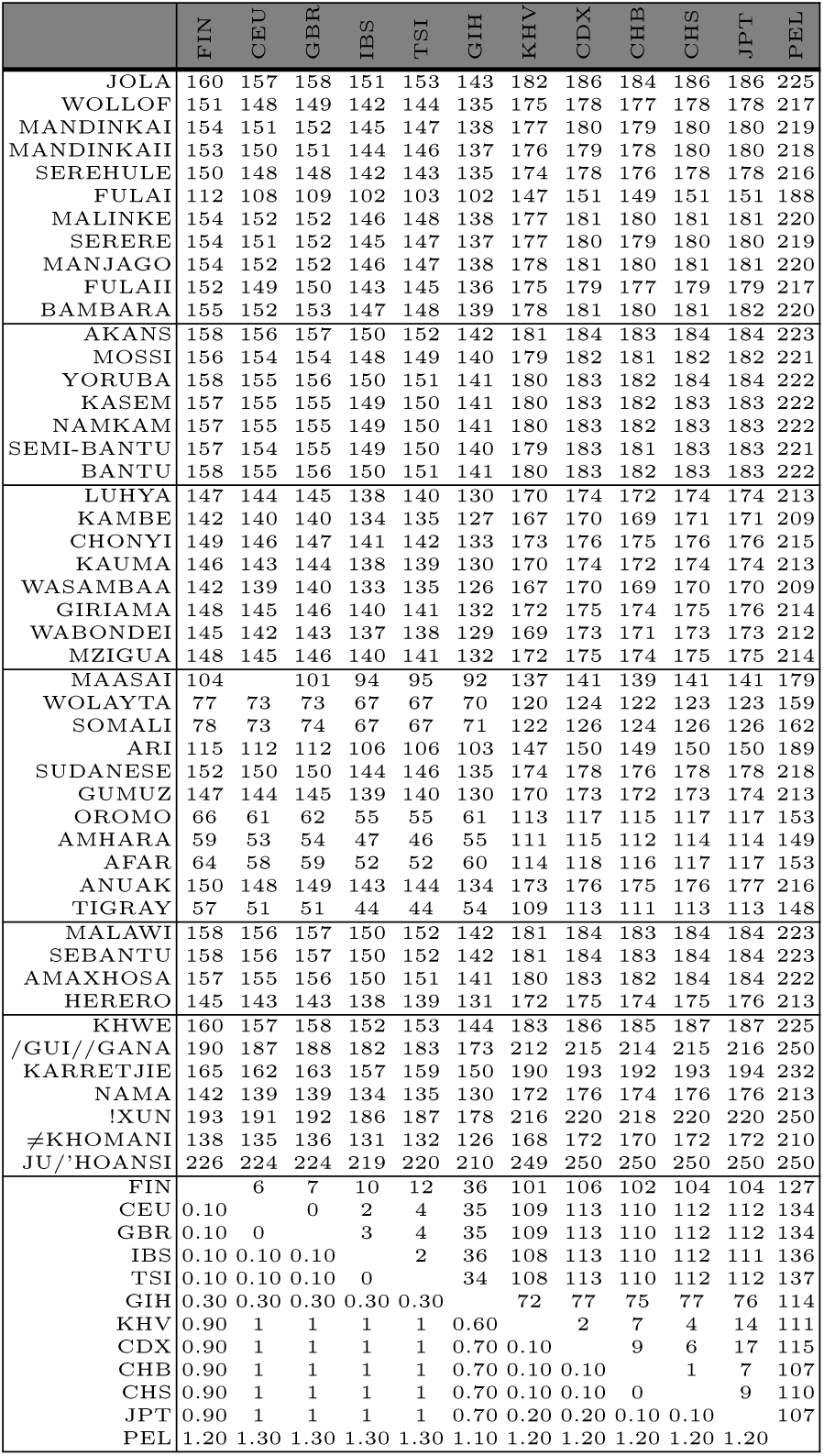
Pairwise *F_ST_* for Eurasian populations. We used smartpca to compute *F_ST_* for each pair of populations, upper left diagonal, together with standard errors computed using a block jacknife. *F_ST_* has been multiplied by 1000

**Figure 2-Source Data 3.**
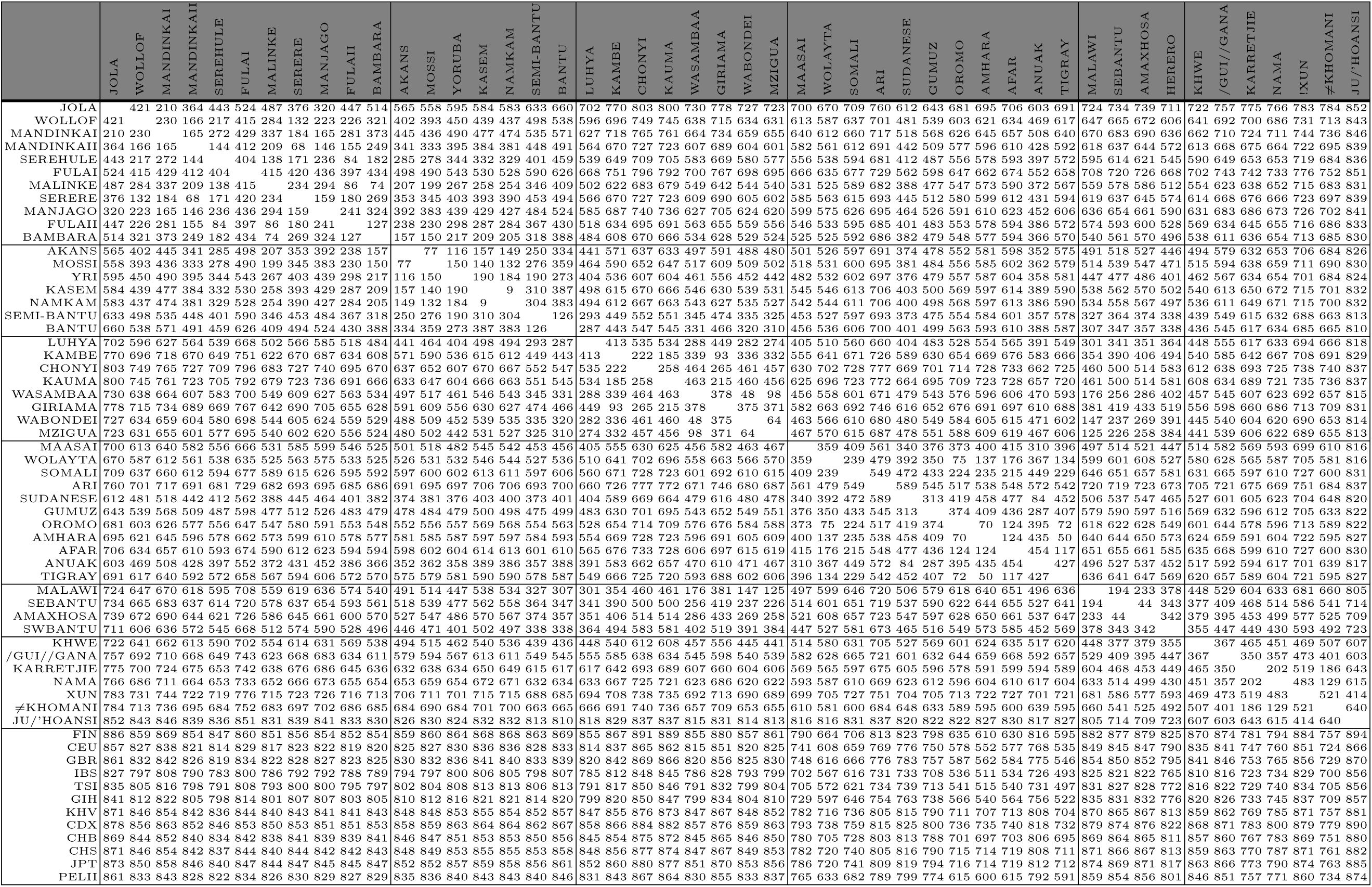
Pairwise *TV D for African populations*. *TV D* has been multiplied by 1000

**Figure 2-Source Data 4.**
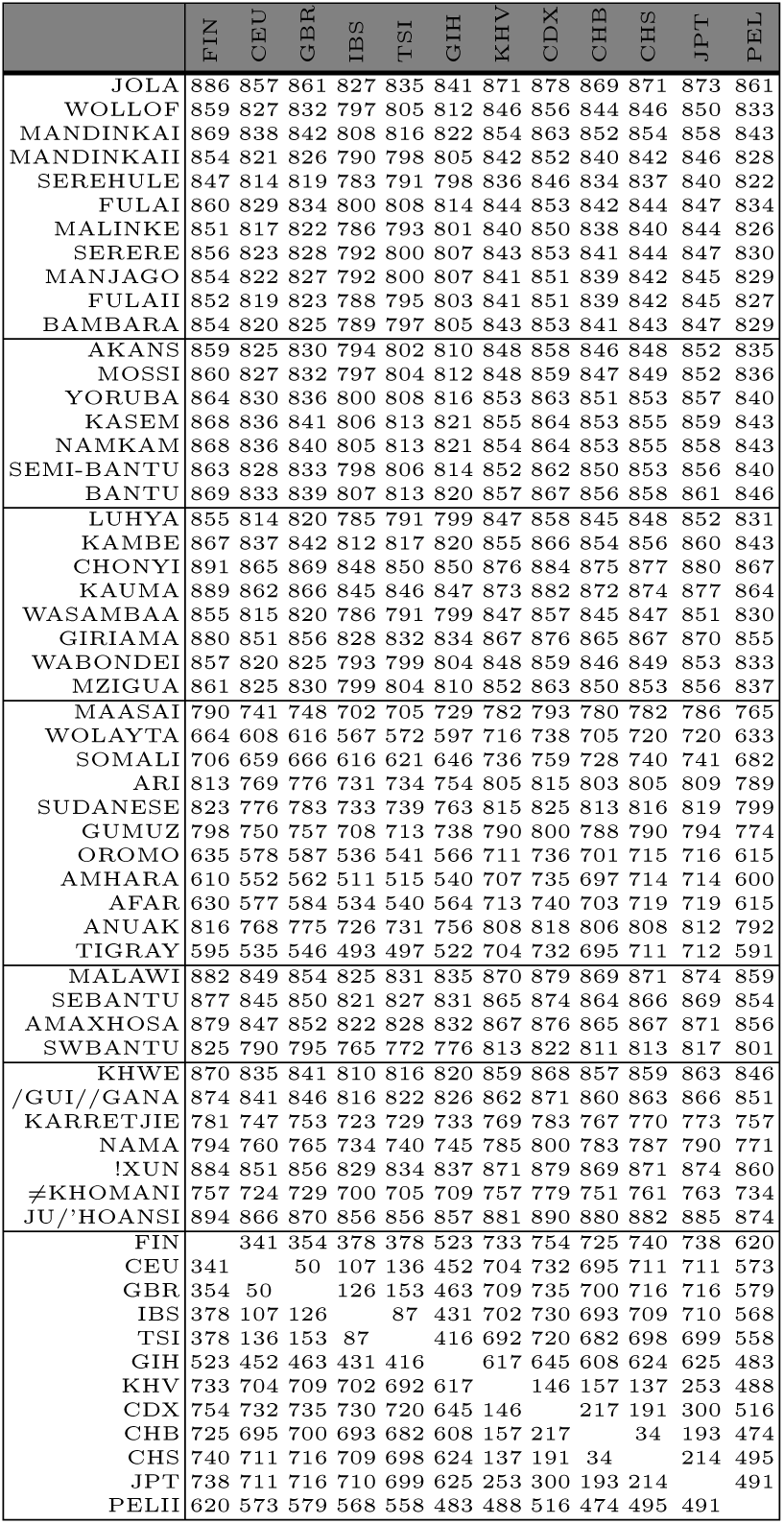
Pairwise *TV D* for Eurasian populations. *TV D* has been multiplied by 1000

**Figure 3-Source Data 1.**
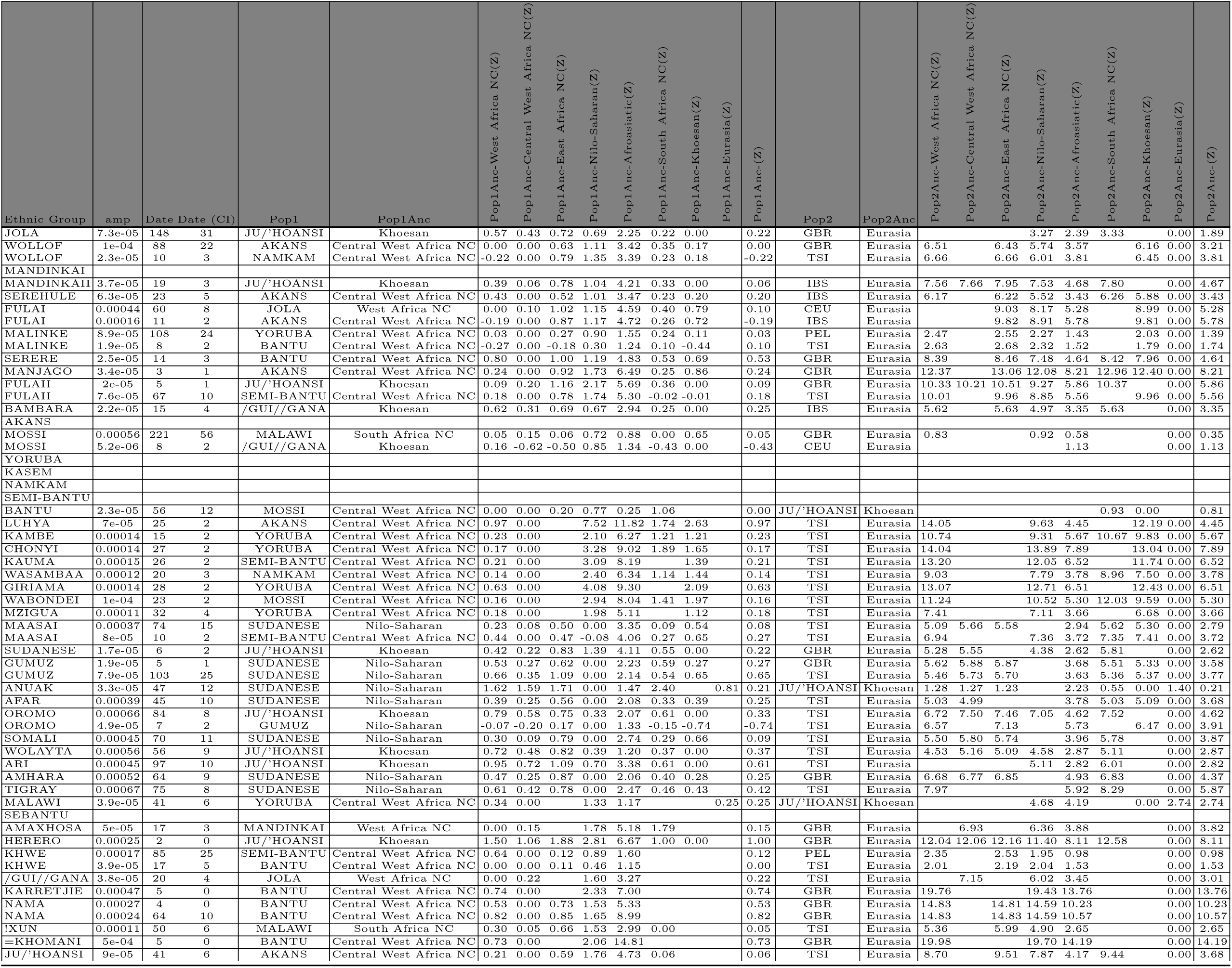
The evidence for multiple waves of admixture in African populations using MALDER and the HAPMAP recombination map. For each event in each ethnic group we show the largest inferred amplitude and date of an admixture event involving two reference populations (Popi and Pop2). We additionally provide the ancestry region identity of the two main reference populations, together with *Z* scores for curve comparisons between this best curve and those containing populations from different ancestry regions. We use a cut-off of *Z* < 2 to decide whether sources from multiple ancestries best describe the admixture source.

**Figure 3-Source Data 2.**
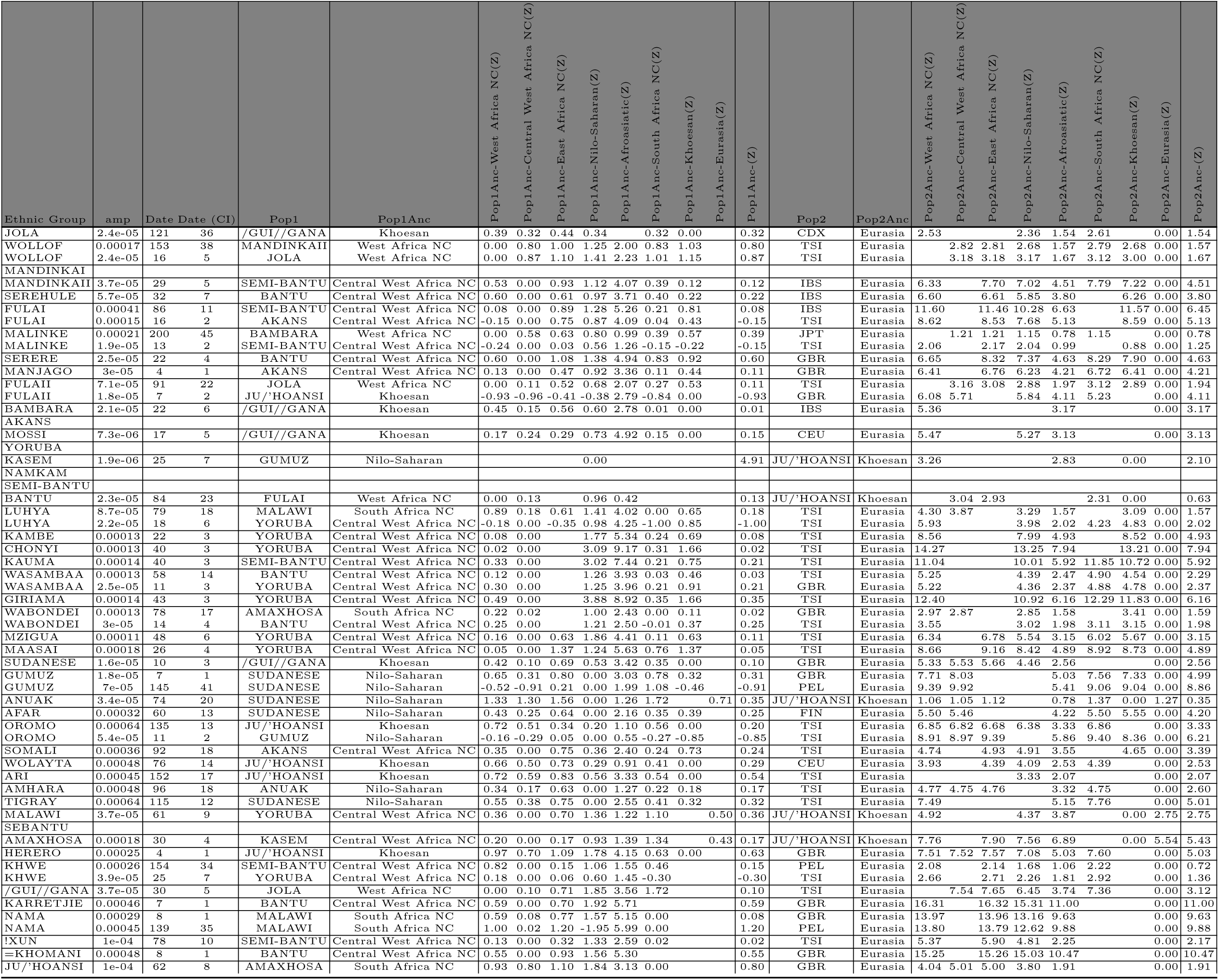
The evidence for multiple waves of admixture in African populations using MALDER and the African recombination map. Columns as in Figure 3-Source Data 1

**Figure 3-Source Data 3.**
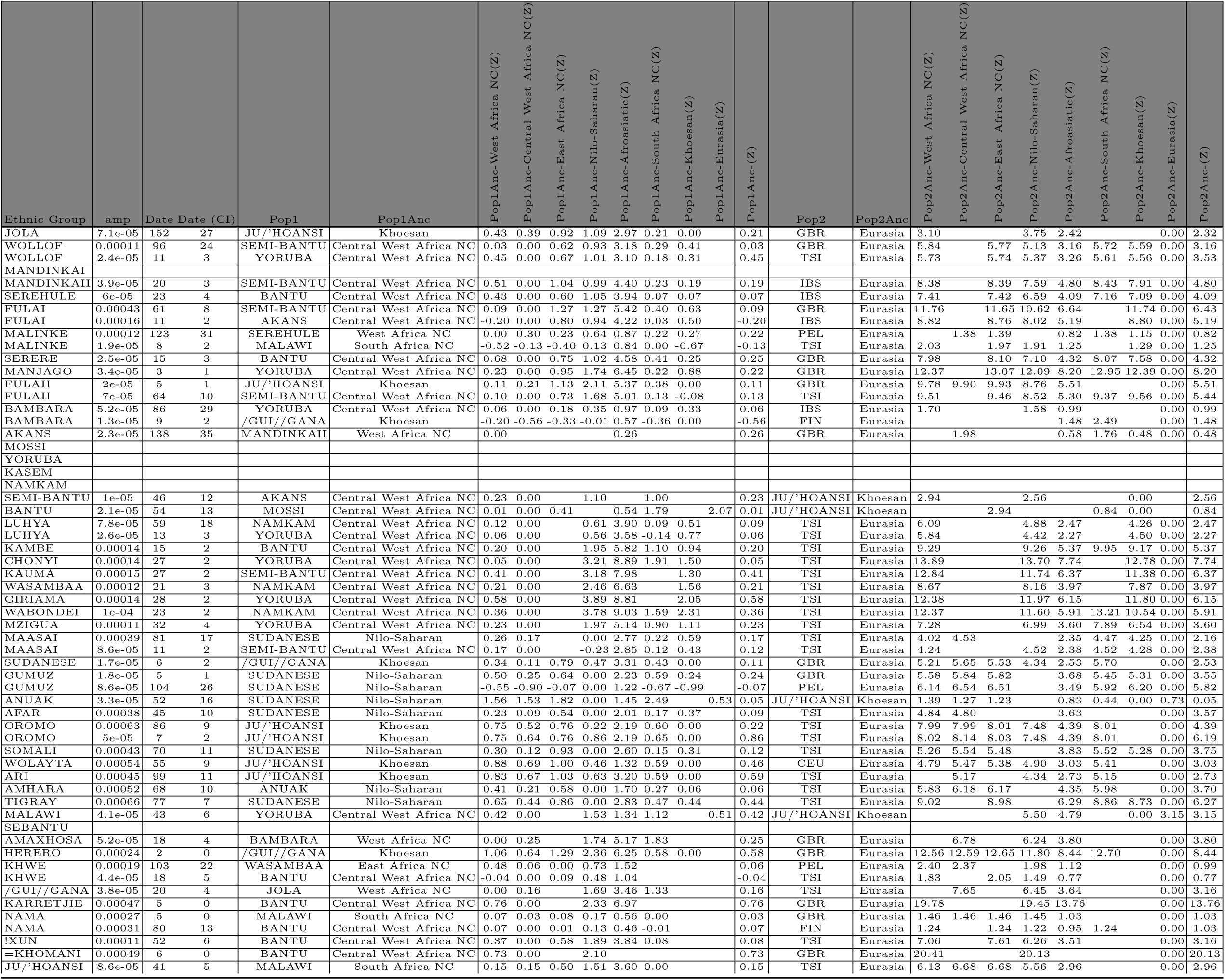
The evidence for multiple waves of admixture in African populations using MALDER and the European recombination map. Columns as in Figure 3-Source Data 1

**Figure 3-Source Data 4.**
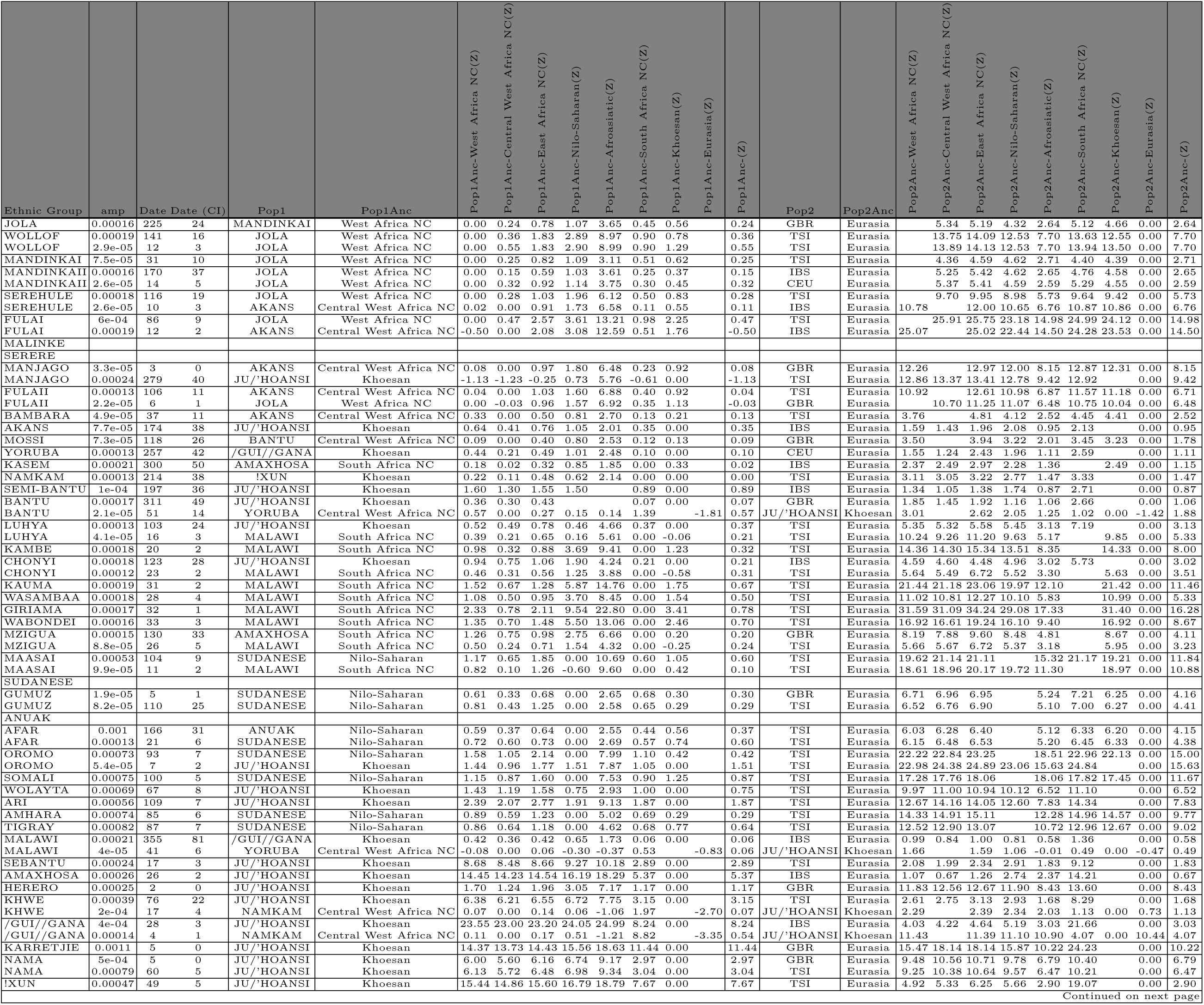
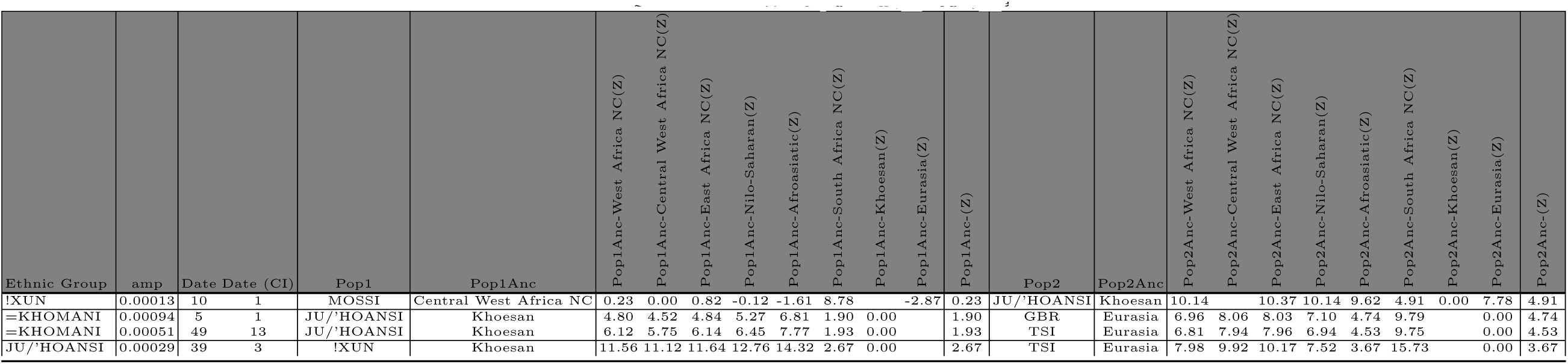
The evidence for multiple waves of admixture in African populations using MALDER and the HAPMAP recombination map and a mindis of 0.5cM. Columns are as in Figure 3-Source Data 1. Here we show the results for the MALDER analysis where we over-ride any short-range LD and define a minimum distance of 0.5cM from which to start computing admixture LD curves

**Figure 4-Source Data 1.**
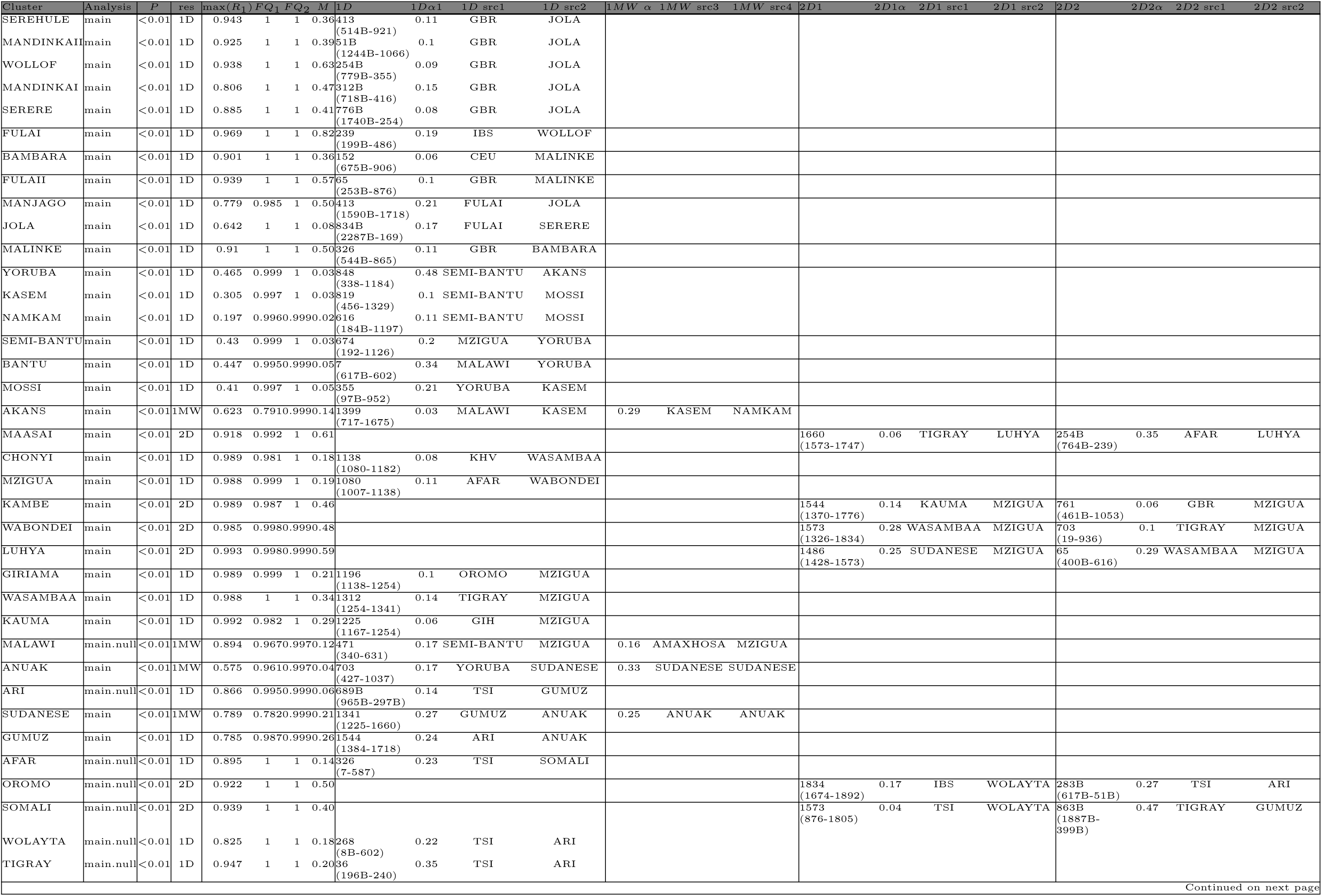
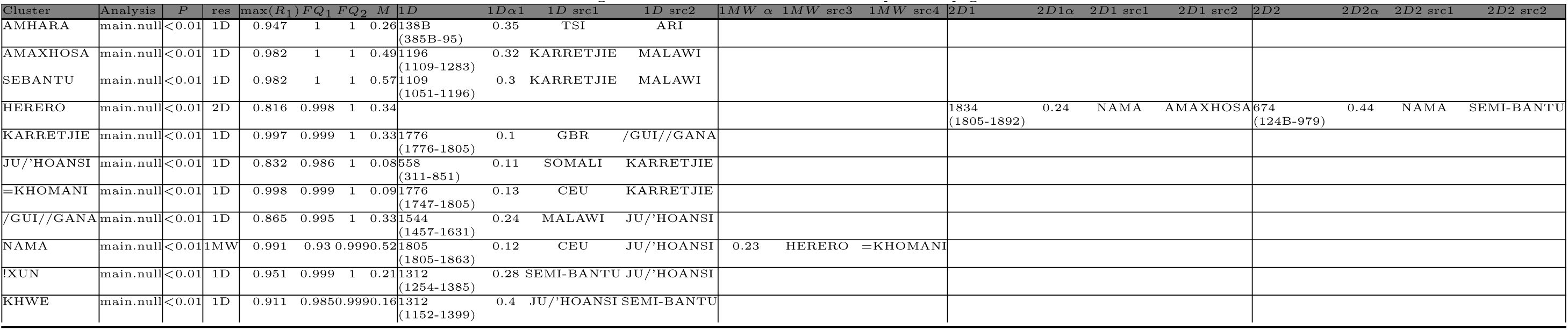
Results of the main GLOBETROTTER analysis. *Analysis* refers to whether the main or masked analysis was used to produce the final result. Admixture *P*-values are based on 100 bootstrap replicates of the NULL procedure. Our resulting inference, *res* can be: 1D (two admixing sources at a single date); 1MW (multiple admixing sources at a single date); 2D (admixture at multiple dates); NA (no-admixture); U (uncertain), max (*P*_1_) refers to the *R*^2^ goodness-of-fit for a single date of admixture, taking the maximum value across all inferred coancestry curves. *FQ*_1_ is the fit of a single admixture event (i.e. the first principal component, reflecting admixture involving two sources) and *FQ*_2_ is the fit of the first two principal components capturing the admixture event(s) (the second component might be thought of as capturing a second, less strongly-signalled event. *M* is the additional *R*^2^ explained by adding a second date versus assuming only a single date of admixture; we use values above 0.35 to infer multiple dates (although see Supplementary Text for details). As well as the final result, for each event we show the inferred dates, αs and best matching sources for ID, 1MW, and 2D inferences. Inferred dates are in years(+ 95% CI; B=BCE, otherwise CE); the proportion of admixture from the minority source (source 1) is represented by *α*. Date confidence intervals are based on 100 bootstrap replicates of the date inference

**Figure 4-Source Data 2.**
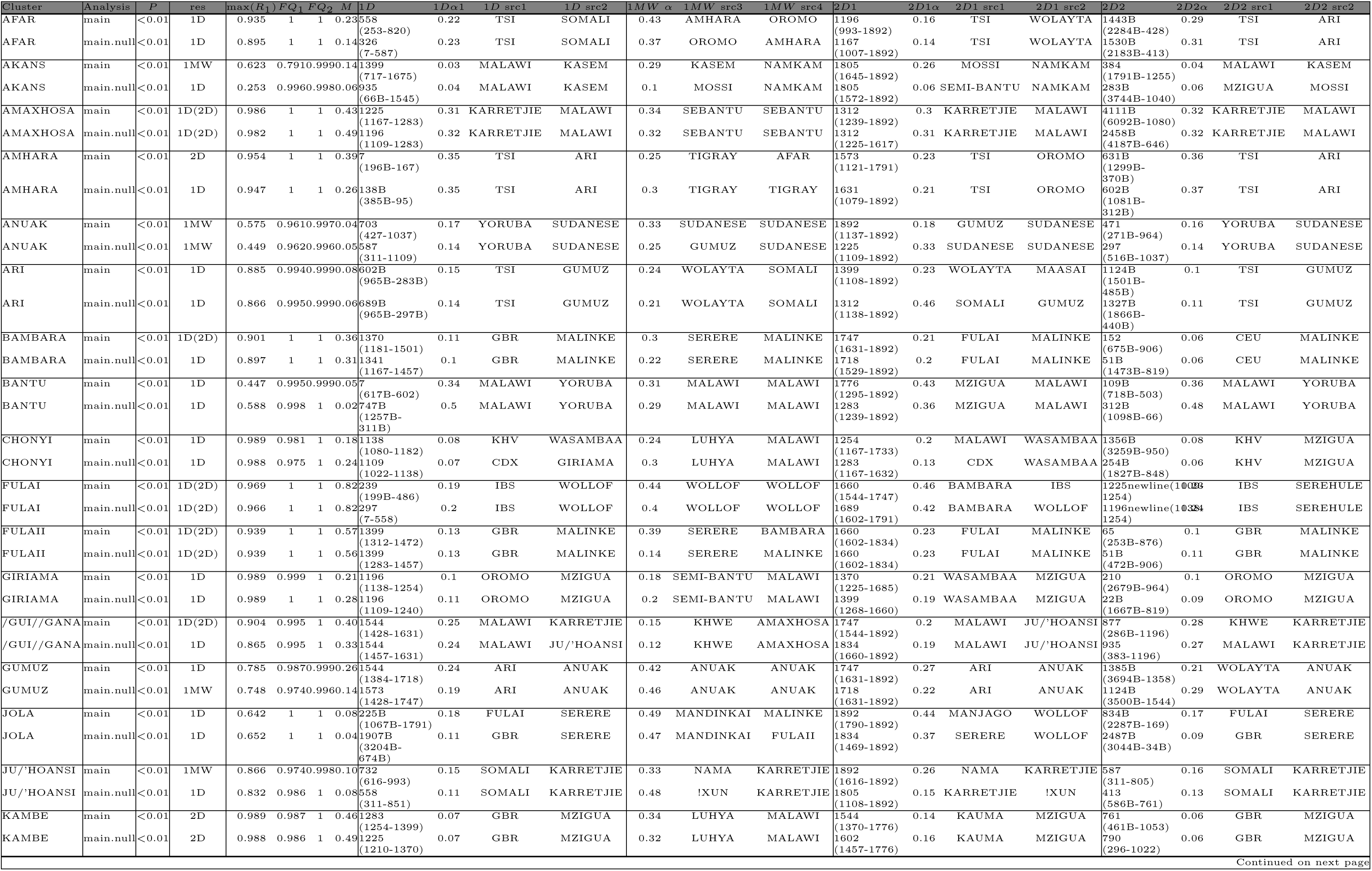
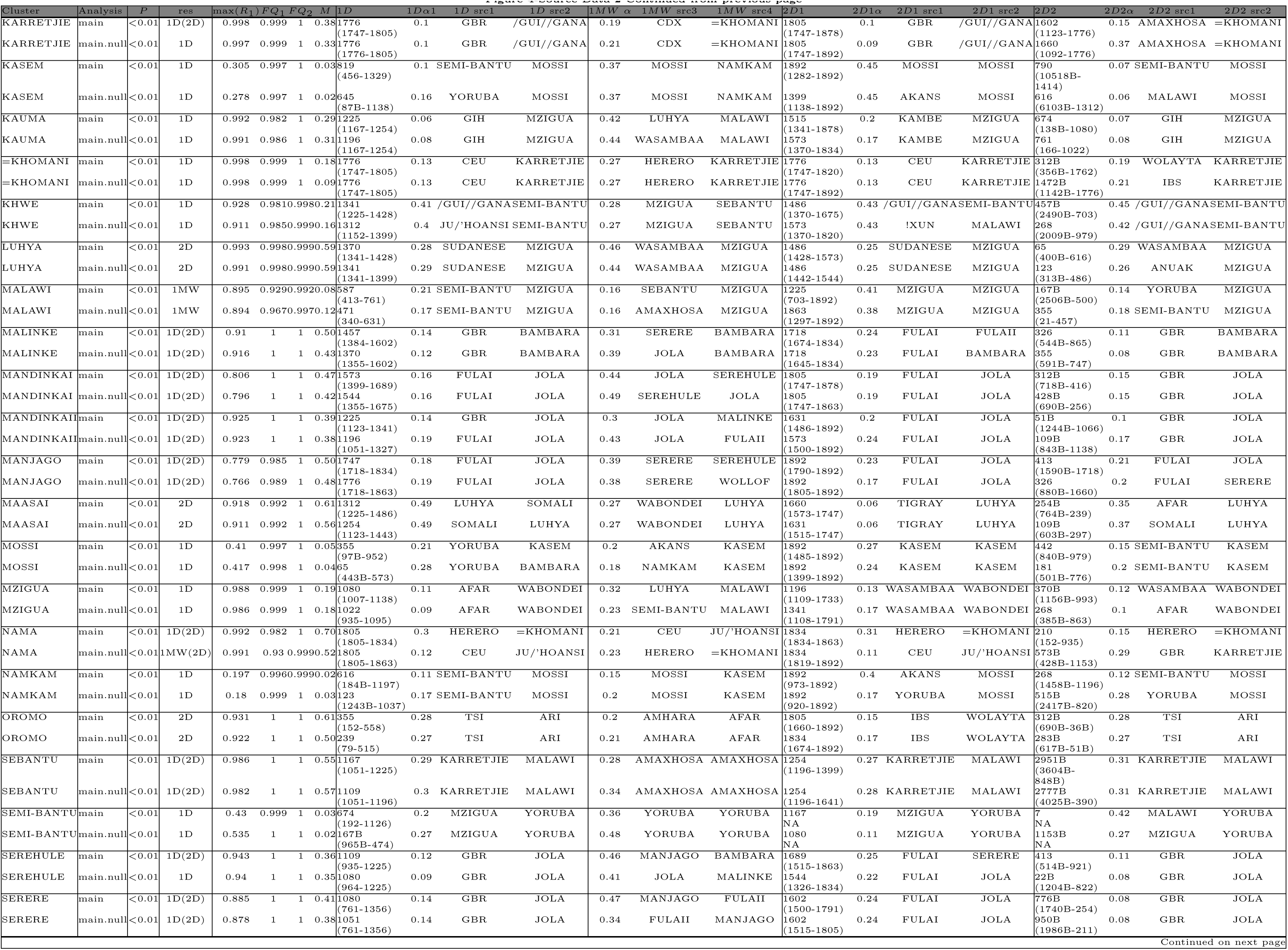
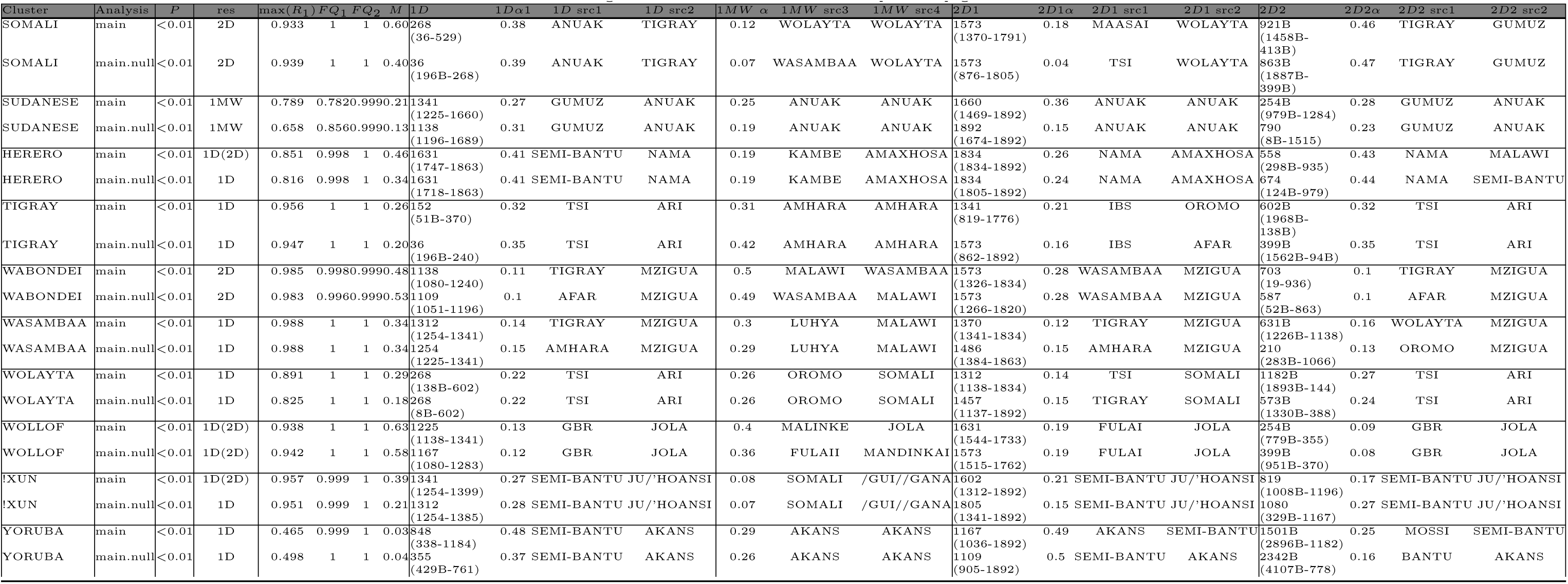
Results of the main GLOBETROTTER analysis. *Analysis* refers to whether the main or masked analysis was used to produce the final result. Admixture *P*-values are based on 100 bootstrap replicates of the NULL procedure. Our resulting inference, *res* can be: ID (two admixing sources at a single date); 1MW (multiple admixing sources at a single date); 2D (admixture at multiple dates); NA (no-admixture); U (uncertain). max(*R*_1_) refers to the *R*_1_ goodness-of-fit for a single date of admixture, taking the maximum value across all inferred coancestry curves. *FQ*_1_ is the fit of a single admixture event (i.e. the first principal component, reflecting admixture involving two sources) and *FQ*_2_ is the fit of the first two principal components capturing the admixture event(s) (the second component might be thought of as capturing a second, less strongly-signalled event. *M* is the additional *R*^2^ explained by adding a second date versus assuming only a single date of admixture; we use values above 0.35 to infer multiple dates (although see Supplementary Text for details). As well as the final result, for each event we show the inferred dates, αs and best matching sources for ID, 1MW, and 2D inferences. Inferred dates are in years(+ 95% CI; B=BCE, otherwise CE); the proportion of admixture from the minority source (source 1) is represented by *α*. Date confidence intervals are based on 100 bootstrap replicates of the date inference

### Supplementary Tables

**Supplementary Table 1.**
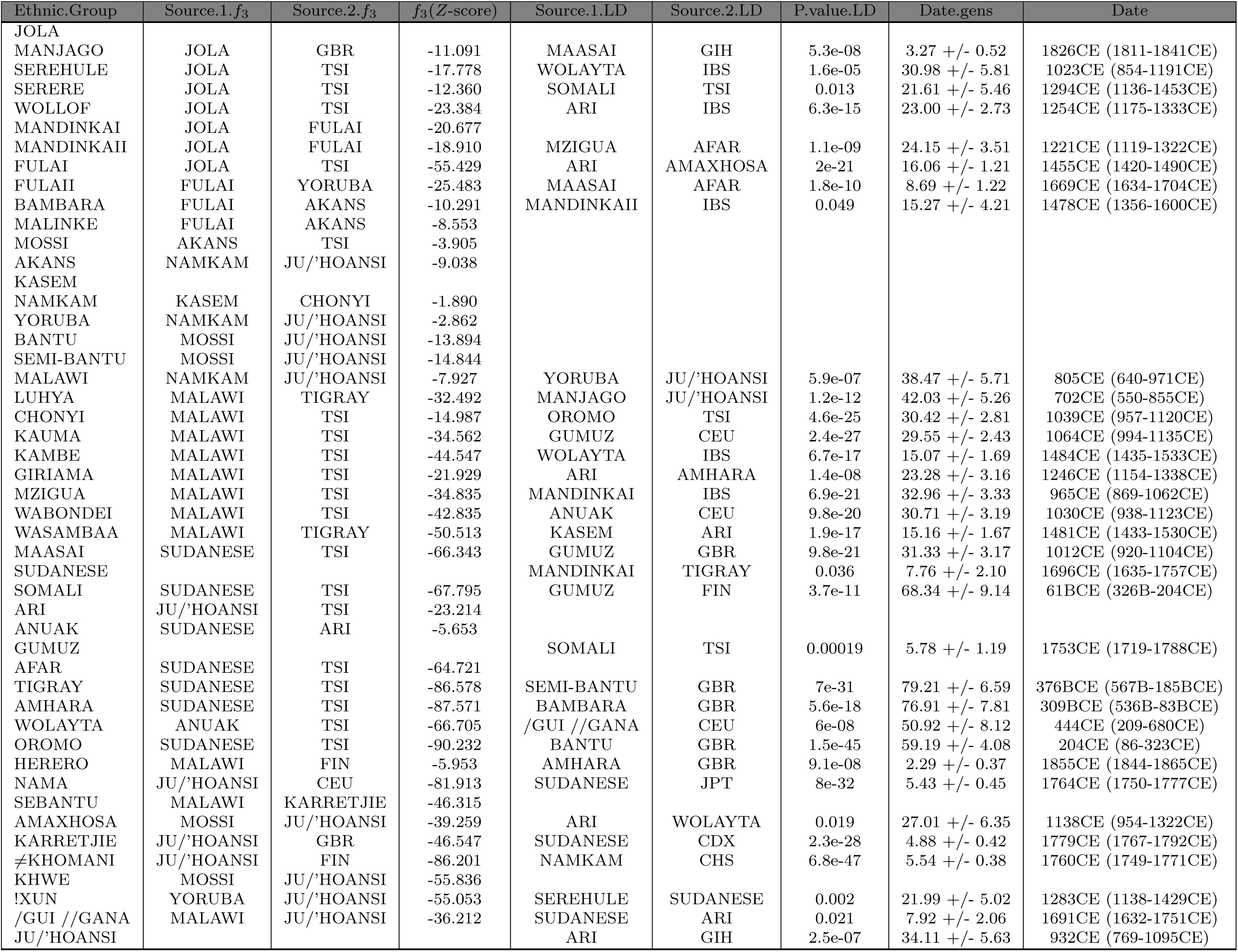
Evidence for admixture across the ethnic groups included in the study using *f*_3_ tests and ALDER. For each group, we report the *f*_3_ test with the most negative value. Source. I.*f*_3_ and Source.2. *f*_3_ refer to the two groups between which gene flow must have occured in the ancestors of the ethnic group under consideration. Groups without results are those where no negative *f*_3_ statistic was found are shown with no results. For the ALDER analysis, we show the two sources of admixture, Source.2.LD and Source.l.LD, with the lowest reported admixture *P*-values. Dates for the ALDER events involving these groups are shown in generations, Date.gens, and in years, Date.

## Footnotes

1 The Gambian Genome Variation Project will sequence a number of full genomes from four Gambian ethnic groups as a basis for improving imputation for future West African specific GWAS

2 data downloaded on 16th October 2013 from ftp://ftp.1000genomes.ebi.ac.uk/vol1/ftp/technical/working/20120131 omni genotypes and intensities/

‡ information accessed on 28th July 2014 at https://www.ethnologue.com/

